# Metastable Dynamics Emerge from Local Excitatory-Inhibitory Homeostasis in the Cortex at Rest

**DOI:** 10.1101/2024.08.24.609501

**Authors:** Francisco Páscoa dos Santos, Paul FMJ Verschure

**Affiliations:** Eodyne Systems SL, 08019, Barcelona, Spain; Universitat Pompeu Fabra, 08018, Barcelona, Spain; Donders Institute for Brain, Cognition and Behavior, Radboud University, 6525 NX, Nijmegen, The Netherlands

**Author notes:** FPS designed and performed the research, analyzed the data, and wrote the manuscript. PFMJV supervised the research and wrote the manuscript. FPS is employed by Eodyne Systems SL. PFMJV is the founder and shareholder of Eodyne Systems S.L., which brings scientifically validated neurorehabilitation and education technologies to society.

**Keywords:** Excitatory-Inhibitory Homeostasis, Cortical Networks, Metastability, Homeostatic Plasticity, Large-Scale Modeling, Complexity

## Abstract

The dynamics of the human cortex are highly metastable, driving the spontaneous exploration of network states. This metastability depends on circuit-level edge-of-bifurcation dynamics, which emerge from firing-rate control through multiple mechanisms of excitatory-inhibitory (E-I) homeostasis. However, it is unclear how these contribute to the metastability of cortical networks. We propose that individual mechanisms of E-I homeostasis contribute uniquely to the emergence of resting-state dynamics and test this hypothesis in a large-scale model of the human cortex. We show that empirical connectivity and dynamics can only be reproduced when accounting for multiple mechanisms of E-I homeostasis. More specifically, while the homeostasis of excitation and inhibition enhances metastability, the regulation of intrinsic excitability ensures moderate synchrony, maximizing functional complexity. Furthermore, the modulation bifurcation modulation by the homeostasis of excitation and intrinsic excitability compensates for strong input fluctuations in connector hubs. Importantly, this only occurs in models accounting for local gamma oscillations, suggesting a relationship between E-I balance, gamma rhythms, and metastable dynamics. Altogether, our results show that cortical networks self-organize toward maximal metastability through the multi-factor homeostasis of E-I balance. Therefore, the benefits of combining multiple homeostatic mechanisms transcend the circuit level, supporting the metastable dynamics of large-scale cortical networks.

---

Complex systems, such as the human brain, show rich collective dynamics beyond the behavior of their units (1–3). One of such collective behaviors in cortical networks is metastability, underlying the spontaneous exploration of the state-space of network configurations (4–10) with a pivotal role in cognitive function (11–14). That said, the key ingredients underlying the emergence of metastable dynamics are not well understood. The connectome can have a pivotal role, either due to its intrinsic inhomogeneity (9) or conduction delays between cortical areas (10, 15), supporting intrinsic activity patterns with high complexity (16–18). However, it has also been suggested that a necessary condition for the emergence of metastability is that the units making up the system are poised near a phase-transition, or bifurcation point (6, 19), underlying the rich patterns of activity observed in the resting-state neocortex (6, 17, 19–24). More specifically, when multiple small-scale phase transitions co-occur, they can self-reinforce and lead to the emergence of collective activity patterns (19). Therefore, while the emergence of metastable dynamics requires the interaction between cortical areas, which is constrained by the connectome, local circuit dynamics must also be key in its genesis. For this reason, we hypothesize that excitatory-inhibitory (E-I) homeostasis, which regulates edge-of-bifurcation dynamics in local cortical circuits (19, 25–29), could be a fundamental principle for understanding the cortex and its collective dynamics (3, 30).

There is extensive evidence that the excitatory and inhibitory inputs received by cortical pyramidal (PY) neurons are subject to a tight balance (31–35), essential for a myriad of operations performed by cortical networks (27, 36–38). For this reason, cortical neurons are equipped with mechanisms of E-I homeostasis, which contribute to the maintenance of E-I balance in the face of perturbations in activity (39–42). These mechanisms include synaptic scaling of excitatory (39, 40, 43) and inhibitory synapses (35, 40, 43, 44) and the adaptation of intrinsic excitability (40, 45, 46). Furthermore, recent results suggest that the conjugation of multiple forms of E-I homeostasis not only ensures stable firing rates but can also regulate the activity of local cortical circuits toward criticality (28, 47), providing a mechanism for the control of edge-of-bifurcation dynamics at the circuit level (29).

In that context, several computational studies have explored how local E-I balance ensures flexible collective dynamics (48, 49) and the emergence of functional connectivity networks (49–51), particularly in the ultra-slow dynamics characteristic of blood-oxygen-level-dependent (BOLD) signals (52). In addition, E-I homeostasis can also be a mechanism of self-organization, driving the recovery of functional connectivity (FC) following lesions to the connectome (53– 55), and the re-emergence of functional properties such as modularity (56). However, a common caveat of such studies is the exclusive focus on the homeostasis of inhibitory synapses, neglecting a possible role for the combined action of multiple mechanisms of homeostasis in the maintenance of circuit dynamics (28, 47). Importantly, our results suggest that multiple mechanisms of homeostasis not only regulate stable firing rates, but also the distance to the bifurcation of local circuits (29), although it is not clear how this shapes large-scale dynamics. Moreso, homeostasis of inhibition alone is not sufficient for the recovery of collective dynamics in lesioned networks (56), suggesting the need for other homeostatic mechanisms.

Here, we aim to investigate the hypothesis that relying on distinct mechanisms of homeostasis at the circuit level is essential to support the emergence of large-scale metastable dynamics in the human cortex. To do this, we study how large-scale models with local E-I homeostasis reproduce empirical FC and FC dynamics (FCD), network dynamics, and the complexity of functional networks. More importantly, we explore the dependence of these collective patterns not only on E-I balance but also on the specific mechanisms employed to maintain it. In this context, our results demonstrate that the only models able to reproduce empirical FC, FCD, network dynamics, and complexity are the ones accounting for the combined homeostasis of excitation, inhibition, and intrinsic excitability. Finally, we also show that networks with these mechanisms of homeostasis can reorganize to recover not only functional networks but also metastable dynamics following focal lesions. Therefore, our results demonstrate that the maintenance of E-I balance not only underlies the emergence of metastable spontaneous dynamics in the cortex, but is also a robust mechanism of self-organization, especially when based on multiple complementary mechanisms of E-I homeostasis.

## Results

To study how E-I homeostasis impacts the resting-state dynamics of the neocortex we constructed a large-scale model considering some of its key features together with local regulation of E-I balance (Figure 1A). The dynamics of cortical areas are modeled with the Wilson-Cowan model of coupled excitatory-inhibitory populations with oscillations in the gamma-band (∼40 Hz). Long-range connectivity and conduction delays between areas follow empirical structural connectivity data derived from diffusion-tensor imaging. Critically, we implement multiple empirically identified mechanisms of local E-I homeostasis (29, 41, 47), which ensure that activity remains close to a target firing rate (*ρ*) by regulating excitatory and inhibitory synapses and the intrinsic excitability of excitatory populations in each cortical area (29) (Figure 1A).

**Fig. 1.**
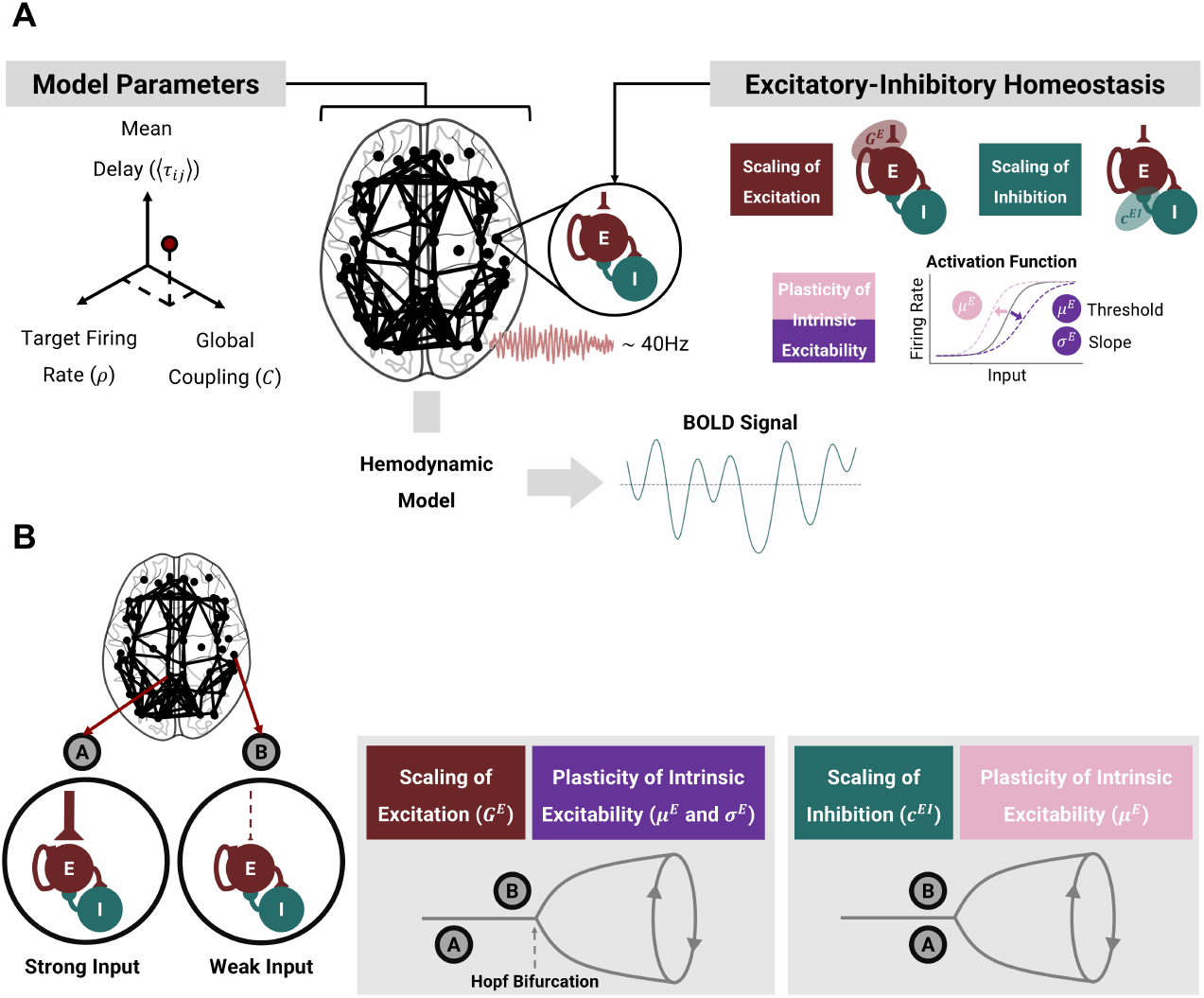
Model Architecture and Local Excitatory-Inhibitory Homeostasis. (A) To simulate the resting-state dynamics of the cortex, we built a large-scale model based on the healthy human connectome. Cortical areas are modeled with the Wilson-Cowan model of coupled excitatory-inhibitory populations, tuned to oscillate at 40 Hz. Based on experimental studies, we implement distinct forms of local E-I homeostasis to maintain the mean excitatory activity at a target firing rate (*ρ*) based on the scaling of excitatory synapses (*G*^*E*^), inhibitory synapses (*c*^*EI*^), or the plasticity of intrinsic excitability, either through the adaptation of the firing threshold (*µ*^*E*^) or the threshold and slope (*µ*^*E*^ and σ^*E*^) of the input-output function of neural populations. In addition, we explore models with different combinations of these homeostatic mechanisms and of the free parameters (global coupling *C*, target firing rate (*ρ*), and mean delay). We extract simulated BOLD signals for further analysis and benchmarking against empirical resting-state recordings. (B) Effects of distinct homeostatic mechanisms on local circuit dynamics of strongly and weakly connected nodes. Certain forms of E-I homeostasis, such as the scaling of excitation and the plasticity of excitability slope (σ^*E*^) and threshold (*µ*^*E*^) modulate the distance to the bifurcation of local cortical circuits, so that areas with a strong average input are poised further away from the bifurcation. Conversely, for the homeostasis of inhibition (*c*^*EI*^) or excitability threshold (*µ*^*E*^), the distance to the bifurcation mostly depends on the target firing rate (*ρ*). For more information, consult (29).

Critically, homeostasis is not implemented dynamically in this work. More specifically, we take advantage of the separation of timescales between the fast circuit dynamics and the ultra-slow homeostatic mechanisms (41, 47) to estimate analytically the parameter values corresponding to the homeostatic fixed point - i.e. the parameters allowing for the maintenance of the target firing rate (*ρ*) across all nodes. This mathematical estimation allows for the models to be initialized with the homeostatic values of each relevant parameter, making the simulations more tractable and saving simulation time. For a more detailed derivation of the methods consult (29). Here, to compute the local parameters of nodes embedded in the large-scale network, we estimate the external input received by each node from the rest of the network, assuming the mean firing rate to be *ρ* in all cortical areas (see Methods).

Importantly, our implementation of the Wilson-Cowan model behaves as a Hopf bifurcation between a fixed point of noisy activity and sustained gamma oscillations. That said, it is relevant to stress that the distinct mechanisms of homeostasis have specific effects on the local dynamics of the model (Figure 1B) (29). More specifically, the scaling of excitation and the plasticity of excitability slope (σ^*E*^) and threshold (*µ*^*E*^) so that circuits with stronger input are poised farther away from the bifurcation than nodes with weaker inputs. Conversely, the plasticity of inhibition (*c*^*EI*^) or the firing threshold (*µ*^*E*^) do not affect the distance to the bifurcation, which depends only on the target firing rate (Figure 1B). Furthermore, when combining multiple homeostatic mechanisms, their effects on edge-of-bifurcation dynamics are also compounded (Figure S1). For a more detailed exploration of the effects of each homeostatic mechanism on node dynamics, refer to (29).

To optimize model performance, we explore different combinations of free parameters, namely the local target firing rate (*ρ*), the global coupling (*C*), which scales the structural connectivity, and the mean conduction delay, which depends on the length of white-matter tracts and the conduction velocity. For each simulation, we first verify the validity of the model by evaluating if the difference between mean local firing rates and the target (*ρ*) does not exceed 1%. We then only retain valid models - i.e. where firing rates can be maintained close to the target - for further analysis. For a detailed exploration of model validity, refer to the Supporting Information (SI Text and Figs S2-11). Finally, we extract simulated BOLD signals from valid networks by using the Balloon-Windkessel hemodynamic model (see Methods) and analyze network dynamics based on the BOLD time series.

### Modulation of Distance to Bifurcation at Circuit Level Optimizes Fit to Empirical Data

We start our analysis by studying how distinct forms of homeostasis, based on the plasticity of excitation, inhibition, and intrinsic excitability (see Methods and Figure 1) shape the performance of our model in reproducing empirical FC and FCD obtained from resting-state BOLD signals, which are strongly modulated by E-I balance (52, 56). We simulate the activity of a large-scale model of the neocortex with different combinations of free parameters (*C, ρ*, and mean delay) and mechanisms of homeostasis (Figure 1). For each simulation, we compare empirical and simulated FC matrices and FCD distributions and extract a cross-feature fitting score to evaluate model performance.

The score is calculated as 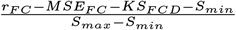, where *r*_*FC*_ is the Pearson’s correlation between empirical and simulated FC matrices, *MSE*_*FC*_ their mean squared error, *KS*_*FCD*_ the Kolmogorov-Smirnov distance between FCD distributions and *S*_*max*_*/S*_*min*_ the maximum/minimum of *r*_*FC*_ − MSE_FC_ − KS_FCD_. With this, we obtain a cross-feature fitting score between 0 and 1, reflecting how well models represent the features of interest (see Methods for more detail). Furthermore, our results remain similar when employing a different method to compute the cross-feature fitting score (Fig S12).

First, it is relevant to point out that the main parameters impacting model performance are *C* and *ρ*, while the mean delay of inter-areal conduction does not have a strong impact, provided it is long enough to avoid widespread pathological synchronization (Figs S13-16 for the full parameter spaces). Therefore we present the performance scores for each combination of *C* and *ρ*, averaged across mean delays (Figure 2A). For each mechanism of homeostasis, we observed a narrow region in the parameter space where empirical resting-state FC and FCD emerge. To compare the optimal performance of each model, we select the combination of *C* and *ρ* yielding the highest fitting score while ensuring that simulations are valid for at least 10 different values of mean delay. Although conduction delays do not strongly impact the fitting score (Figs S13-16), we enforce this constraint to ensure that models are robust to changes in conduction velocity without losing the ability to maintain stable firing rates. That said, we present the distribution of cross-feature fitting scores across mean delays for the optimal point of each model (Figure 2B, Table 1) and examples of activity, FC and FCD distributions from empirical and simulated data at the optimal point of each model (Figure 2C). The best scores are obtained for models with *G*^*E*^ homeostasis (0.855 ± 0.009) and *G*^*E*^ + *c*^*EI*^ homeostasis (0.851 ± 0.019), followed by homeostasis of *G*^*E*^, *c*^*EI*^ and *µ*^*E*^ + σ^*E*^ (i.e. All) (0.849 ± 0.009). The values of *r*_*FC*_, *MSE*_*FC*_, and *KS*_*FCD*_ corresponding to each optimal score can be consulted in Table 1. While the performance of the model with *G*^*E*^ homeostasis was significantly better than the one with all mechanisms of homeostasis (p=0.020; Mann-Whitney u-test, false discovery rate (FDR) correction), the effect size is not substantial (Cohen’s d=0.281). Furthermore, the optimal target firing rates for the three best models (*G*^*E*^: 0.12; *G*^*E*^ + *c*^*EI*^ : 0.1; All: 0.12) are close to the bifurcation between damped and sustained oscillations (Fig S1), in line with theoretical (19, 21, 22) and experimental results suggesting that the neocortex operates at the edge-of-bifurcation.

**Table 1.** Optimal parameters, fitting scores, dynamics, and complexity for each mechanism of E-I homeostasis in comparison to empirical data. Fitting scores correspond to 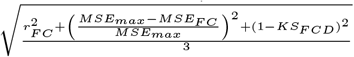, where *r*_*FC*_ is the Pearson’s correlation between empirical and simulated FC matrices, *MSE*_*F C*_ their mean squared error, *KS*_*F CD*_ the Kolmogorov-Smirnov distance between FCD distributions and *S*_*max*_*/S*_*min*_ the maximum/ minimum of *r*_*F C*_ − *MSE*_*F C*_ − *KS*_*F CD*_. Values are presented as mean±sd. Values in bold represent a significant difference from empirical data (p<0.05, Mann-Whitney U-test, FDR correction)

| Homeostatic Mechanism | $C$ | $\rho$ | Score | $r_{FC}$ | $MSE_{FC}$ | $KS_{FCD}$ | Synchrony | Metastability | Complexity |
| --- | --- | --- | --- | --- | --- | --- | --- | --- | --- |
| $G^E$ | 5.99 | 0.12 | 0.86±0.01 | 0.49±0.01 | 0.061±0.010 | 0.16±0.04 | **0.661±0.027** | 0.201±0.009 | **0.640±0.017** |
| $c^{EI}$ | 2.15 | 0.05 | 0.64±0.01 | 0.28±0.02 | 0.158±0.088 | 0.91±0.04 | **0.203±0.021** | **0.100±0.007** | **0.393±0.025** |
| $\mu^E$ | 0.46 | 0.09 | 0.67±0.02 | 0.31±0.02 | 0.139±0.005 | 0.84±0.08 | **0.253±0.017** | **0.116±0.009** | **0.436±0.025** |
| $\mu^E + \sigma^E$ | 3.59 | 0.07 | 0.84±0.01 | 0.43±0.02 | 0.056±0.002 | 0.18±0.03 | 0.507±0.015 | **0.173±0.008** | 0.767±0.021 |
| $G^E + c^{EI}$ | 7.74 | 0.11 | 0.85±0.02 | 0.48±0.04 | 0.088±0.031 | 0.14±0.05 | **0.707±0.041** | 0.203±0.011 | **0.565±0.052** |
| $G^E + c^{EI} + \mu^E$ | 1.00 | 0.10 | 0.82±0.01 | 0.42±0.01 | 0.063±0.002 | 0.28±0.05 | 0.533±0.014 | **0.171±0.007** | **0.797±0.020** |
| All ($G^E + c^{EI} + \mu^E + \sigma^E$) | 3.59 | 0.12 | 0.85±0.01 | 0.47±0.02 | 0.046±0.003 | 0.18±0.04 | 0.580±0.021 | 0.191±0.008 | 0.750±0.020 |
| Empirical | - | - | - | - | - | - | 0.537±0.112 | 0.194±0.025 | 0.754±0.069 |

**Fig. 2.**
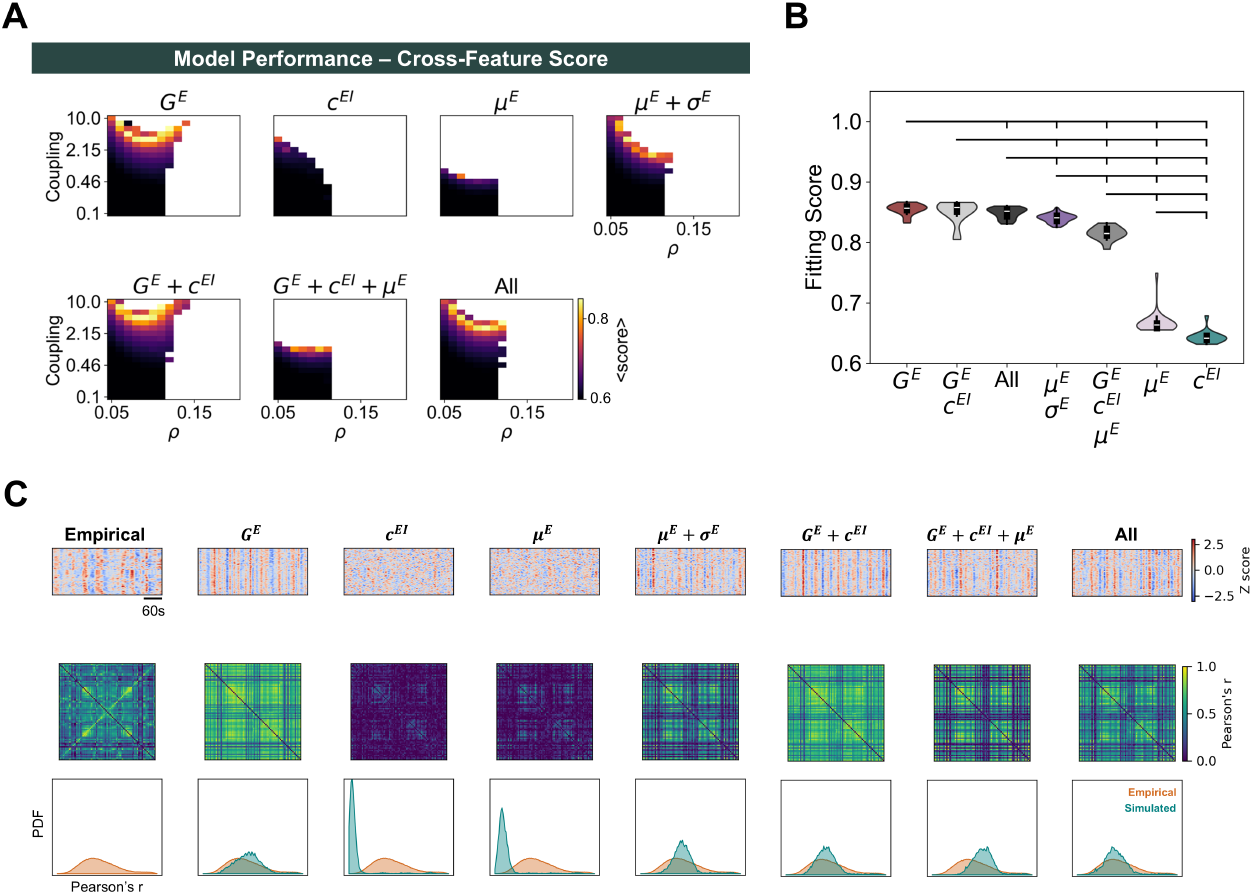
Cross-feature model performance for different homeostatic mechanisms. (A) Parameter spaces representing the cross-feature score for each combination of *C* and *ρ*, averaged across mean delays. Blank spaces represent combinations of *C* and *ρ* for which the homeostatic set point is not valid (i.e. mean firing rates differ from *ρ* by more than 1% in at least one cortical area) (B) Comparison between cross-feature fitting scores at the optimal point of each mechanism of homeostasis. Distributions correspond to simulations with optimal *C* and *ρ* and all values of mean delay yielding a valid network solution. Brackets indicate a significant difference, with p<0.05 from a Mann-Whitney U-test. All p-values were FDR-corrected. (C) Examples of network activity (Top), functional connectivity matrices (middle), and FCD distributions (bottom) from empirical data and optimized models with distinct combinations of homeostatic mechanisms. In each plot, we present 6 minutes of band-pass filtered and Z-scored activity across all cortical areas. Each example from the optimal models represents a simulation with the optimal combination of *C* and *ρ* and a mean delay of 4ms.

Our results indicate that models with E-I homeostasis can reproduce the emergence of empirical networks and their dynamics at the ultra-slow timescales characteristic of resting-state BOLD, depending on the interaction between global coupling (*C*) and target firing rate (*ρ*). In addition, empirical data is best approximated when accounting for mechanisms of homeostasis that modulate the bifurcation point (i.e. the ones that modulate *G*^*E*^ or σ^*E*^) (29), so that areas with stronger inputs are poised farther away from the bifurcation (Figures 1B and S1). This effect compensates for the higher input fluctuations in strongly connected nodes (Fig S17), ensuring that such nodes are not constantly entering the regime of sustained oscillations. Therefore, even though this effect leads to heterogeneous distances to the bifurcation across cortical areas, we argue that our results still align with the edge-of-bifurcation theory of brain dynamics (6, 17, 19–24) when input fluctuations are accounted for. More importantly, this bifurcation modulation, observed in models with the homeostasis of excitation (*G*^*E*^) and excitability slopes (σ^*E*^) (Figures 1B and S1), is essential for the emergence of FC and FCD (Figure 2).

### Resting-State Functional Connectivity Emerges in Networks with Metastable Dynamics

Given that the metastability of large-scale cortical networks is shaped by circuit-level edge-of-bifurcation dynamics (6, 19, 22, 49), controlled by E-I homeostasis (29, 47), it is relevant to explore how distinct mechanisms of homeostasis shape the collective dynamics of the resting-state cortex. To do this, we measure the synchrony and metastability (see Methods) of the simulated BOLD signals of models under different homeostatic mechanisms and combinations of *C* and *ρ*. Again, the conduction delays do not significantly impact either synchrony or metastability (Figs S18-19) and, for this reason, we present results averaged across mean delays.

We observe that synchrony generally follows increases in *C* and *ρ*, reaching a maximum before the networks become unstable (Fig S20). To explore how synchrony is shaped by E-I homeostasis, we compare the optimal models of each mechanism of homeostasis (i.e. best cross-feature fit) with empirical data (Figure 3A, Table 1). While some of the models that best reproduce empirical FC and FCD generally display moderate levels of synchrony, models based on the homeostasis of *G*^*E*^ + *c*^*EI*^ (0.707 ± 0.041) or *G*^*E*^ (0.661 ± 0.027) show significantly higher synchrony than empirical resting-state BOLD dynamics (0.537 ± 0.112) (p<0.001, Mann-Whitney U-test, FDR corrected). In contrast, networks based on the combination of all mechanisms of homeostasis, which reproduce empirical FC and FCD to a similar degree (Figure 2B), show levels of synchrony similar to empirical data (0.580 ± 0.021; p=0.108; Mann-Whitney U-test, FDR corrected).

**Fig. 3.**
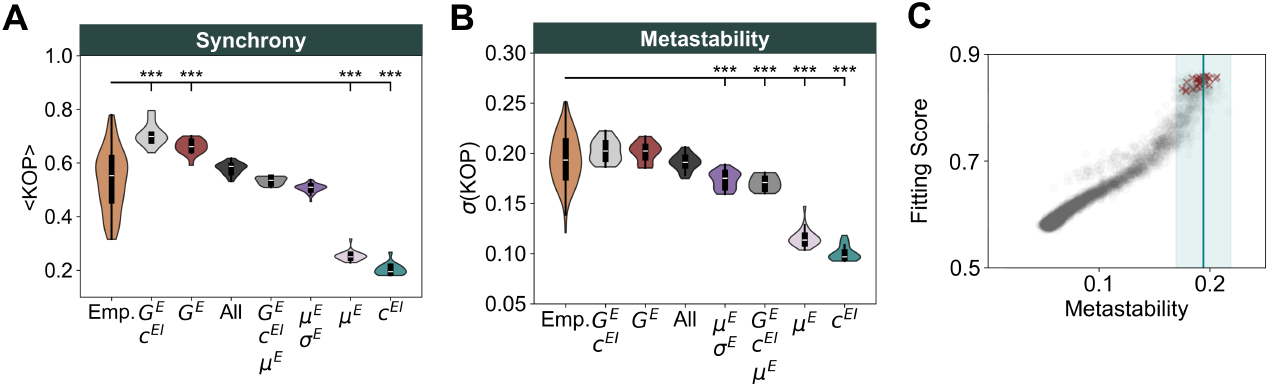
Networks with multiple mechanisms of homeostasis reproduce empirical network dynamics. (A) Comparison between synchrony in empirical data and models at the optimal point (best cross-feature fitting score) for each mechanism of homeostasis. Simulated distributions correspond to models with optimal *C* and *ρ* and all values of mean delay yielding a valid solution. Brackets represent a significant difference between models and empirical data from a Mann-Whitney U-test (^*^ p<0.05; ^**^ p<0.01; ^***^ p<0.001). All p-values were FDR-corrected. (B) Same as (A), for metastability. (C) Relationship between fitting score and metastability in the model with homeostasis of excitation, inhibition, and intrinsic excitability (*µ*^*E*^ and σ^*E*^). Gray dots show the results of all simulations while red crosses represent simulations at the optimal point (*C* = 3.59, *ρ* = 0.12). The vertical bar and shaded area display the mean and standard deviation of metastability in empirical BOLD signals.

Regarding metastability, our results reveal a region of maximal metastability determined by the interaction between *C* and *ρ* (Fig S21), coinciding with the region of optimal fit to empirical data (Figure 2A). Furthermore, the mechanisms of homeostasis that best fit empirical FC and FCD also show levels of metastability close to empirical values (0.194 ± 0.025) (Figure 3B, Table 1). More specifically, we found enhanced metastability in models with homeostasis of *G*^*E*^ + *c*^*EI*^ (0.203 ± 0.011), *G*^*E*^ (0.201 ± 0.009), and the combination of all mechanisms of E-I homeostasis (0.191 ± 0.008), with no significant difference from empirical values (*G*^*E*^ + *c*^*EI*^ : p=0.254; *G*^*E*^: p=0.254; All: p=0.521; Mann-Whitney U-test, FDR corrected). Importantly, empirical levels of both synchrony and metastability can only be reproduced when including all mechanisms of homeostasis (Figure 3 and S22). In addition, we found a pronounced positive correlation between the metastability of models and their fitting scores, not only in the model with all mechanisms of homeostasis (Figure 3C) but as a general principle (Fig S23). Therefore, our results confirm previous studies proposing that the resting-state dynamics of cortical networks might correspond to a regime of maximal metastability (22, 49, 57).

In summary, our analysis of network dynamics suggests that empirical FC networks and dynamics emerge more prominently in networks with moderate synchrony and high metastability, which can only be replicated when combining three mechanisms of E-I homeostasis (*G*^*E*^, *c*^*EI*^ and *µ*^*E*^ + σ^*E*^). More importantly, our results show that, through the homeostatic regulation of firing rates, cortical networks self-organize toward a regime of maximal metastability where the empirical resting-state networks emerge.

### Functional complexity is shaped by E-I balance, metastability, and connectome topology

The complexity of spontaneous activity in the human brain is strongly related to the topology of its structural networks (16–18). However, the architecture of the connectome is likely not the only source of complexity in cortical networks. Given the relevance of E-I homeostasis at both the local and global scales, determining how local circuits engage in collective dynamics, E-I balance might be key in the emergence of complex functional interactions between cortical areas. Therefore, we study how different mechanisms of E-I homeostasis impact FC complexity, using the methodology presented in (18) (see Methods).

Our results on FC complexity suggest that, for all mechanisms of homeostasis, it is maximized in the region of the parameter space corresponding to heightened metastability and optimal fit to empirical FC and FCD (Figs S24 and S25). Comparing complexity in the models that best fit empirical FC and FCD data (Figure 4A, Table 1), our results indicate that the combined homeostasis of *G*^*E*^, *c*^*EI*^ and *µ*^*E*^ (0.797 ± 0.020) maximizes FC complexity, although the resulting values are significantly higher than in empirical data (p=0.014; Mann-Whitney U-test, FDR corrected). Conversely, some of the homeostatic mechanisms with the best fit to empirical FC and FCD are associated with levels of complexity significantly lower than empirical values (*G*^*E*^: 0.640 ± 0.017, p<0.001; *G*^*E*^ + *c*^*EI*^ : 0.565 ± 0.052, p<0.001; Mann-Whitney U-test, FDR-corrected). However, models relying on the combination of all forms of homeostasis reach levels of FC complexity (0.750 ± 0.020; p=0.026) consistent with empirical data. Therefore, accounting for multiple mechanisms of homeostasis is necessary not only to reproduce synchrony and metastability but also functional complexity. Importantly, we found a strong correlation between metastability and complexity, not only for the model with all mechanisms of homeostasis (Figure 4B) but also across models (Fig S23). This suggests that heightened metastability of resting-state brain dynamics (22, 49) results in highly complex activity patterns that reflect on the variability of FC. More importantly, this requires the combined action of multiple mechanisms of E-I homeostasis at the circuit level.

**Fig. 4.**
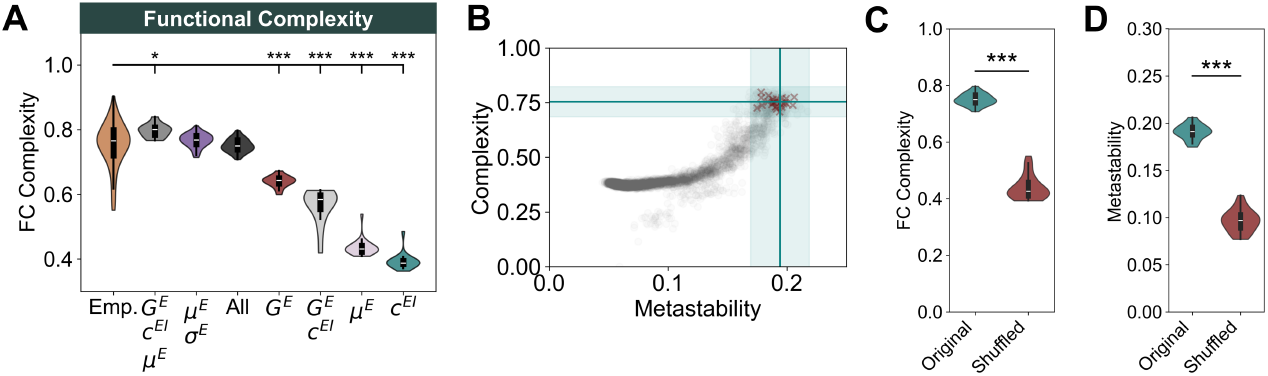
Functional complexity reflects E-I homeostasis, metastability, and connectome topology. (A) Comparison between functional complexity in empirical data and models at the optimal point (best cross-feature fit) for each mechanism of homeostasis. Distributions correspond to models with optimal *C* and *ρ* and all values of mean delay yielding a valid system. Brackets represent a significant difference between models and empirical data from a Mann-Whitney U-test (^*^ p<0.05; ^**^ p<0.01; ^***^ p<0.001). All p-values were FDR-corrected. (B) Relationship between complexity and metastability in the model with homeostasis of excitation, inhibition, and intrinsic excitability (*µ*^*E*^ and σ^*E*^). Gray dots show the results of all simulations while red crosses represent simulations at the optimal point (*C* = 3.59, *ρ* = 0.12). The horizontal and vertical bars and shaded areas display the mean and standard deviation of complexity and metastability, respectively, in empirical BOLD signals. (C) Comparison between FC complexity in models with the original and shuffled structural connectomes and homeostasis of *G*^*E*^, *c*^*EI*^, *µ*^*E*^, and σ^*E*^. Simulations were performed with *C* = 3.59 and *ρ* = 0.12 and all mean delays yielding valid solutions. Asterisks represent the significance of a Mann-Whitney U-test between samples (^***^ p<0.001). (D) Same as C, for metastability.

Having investigated the relationship between E-I homeostasis and complexity compare complexity in models with the empirical and shuffled structural connectomes to assert the relevance of network topology (16–18). Since shuffling preserves the distribution of edge weights, the complexity of the original and shuffled structural matrices is the same (Fig S26). That said, our results show that the complexity of FC in models with the empirical connectome is significantly higher than those with its shuffled counterpart (Original: 0.750 ± 0.020; Shuffled: 0.439 ± 0.042; p<0.001; Mann-Whitney U-test) (Figure 4c). Therefore, in line with (16–18), the complexity of functional interactions also depends on the topology of structural connectivity. However, we point out that, given the relationship between metastability and FC complexity, this result could be an effect of decreased metastability in models with a shuffled connectome (Figure 4d). Therefore, our results suggest that while E-I homeostasis maximizes metastability and complexity by regulating edge-of-bifurcation dynamics in local cortical circuits, the role of the structural connectome in constraining their interactions should not be overlooked.

In summary, our results indicate that the processes underlying the emergence of metastable dynamics also maximize the complexity of spontaneous activity in the cortex. More importantly, our results suggest that multiple mechanisms of homeostasis (modulating excitatory and inhibitory synapses and intrinsic excitability through *µ*^*E*^ and σ^*E*^) are required to reproduce the various dynamical signatures of cortical networks (Table 1). Therefore, we propose that the role of multiple mechanisms of E-I homeostasis does extend beyond the local circuit level (42, 47), shaping the collective dynamics of large-scale cortical networks.

### Effects of E-I Homeostasis on Collective Dynamics Depend on Fast Local Rhythms

The version of the Wilson-Cowan implemented to simulate the local dynamics of cortical areas is based on the fast dynamics of gamma oscillations (∼40Hz). These rhythms are generated by the recurrent loop between E-I populations, with fast inhibition provided by PV interneurons and fast excitation through *α*-amino-3-hydroxy-5-methyl-4-isoxazolepropionic (AMPA) receptors (58–60). While the regulation of AMPA receptors has an important participation in the homeostasis of excitatory synapses [REFS], experimental results show that the slower N-methyl-D-aspartate (NMDA) receptors are also modulated to maintain the homeostasis of firing rates [REFS]. Therefore, it is relevant to investigate if the effects of local E-I homeostasis on the emergence of FC, FCD, and collective metastable dynamics are limited to its impact on fast gamma rhythms, related to AMPA receptors, or can also be observed when accounting for the slower NMDA dynamics.

To investigate this, we implemented a large-scale model with local dynamics based on the reduced Wong-Wang model (50), where excitatory synapses have slower time constants based on the timescales of NMDA receptors (Figure 5A). First, following the framework of (29), previously implemented for the Wilson-Cowan model, we derive the analytical expressions to calculate the parameter values at the homeostatic fixed point for each empirically derived mechanism of homeostasis. For more details on the derivation, refer to the Supplementary Methods and Fig S27. Importantly, since the timescale of excitatory synapses (100 ms) is substantially slower than the timescale of inhibition (10 ms), the Wong-Wang model does not have a Hopf-bifurcation. Therefore, the local dynamics dependent on NMDA receptors remain in the stable regime for all tested combinations of external input and target firing rate, regardless of the combination of homeostatic mechanisms (Fig S28), as opposed to the strong bifurcation modulation observed for the Wilson-Cowan model (Figures 1B and S1).

**Fig. 5.**
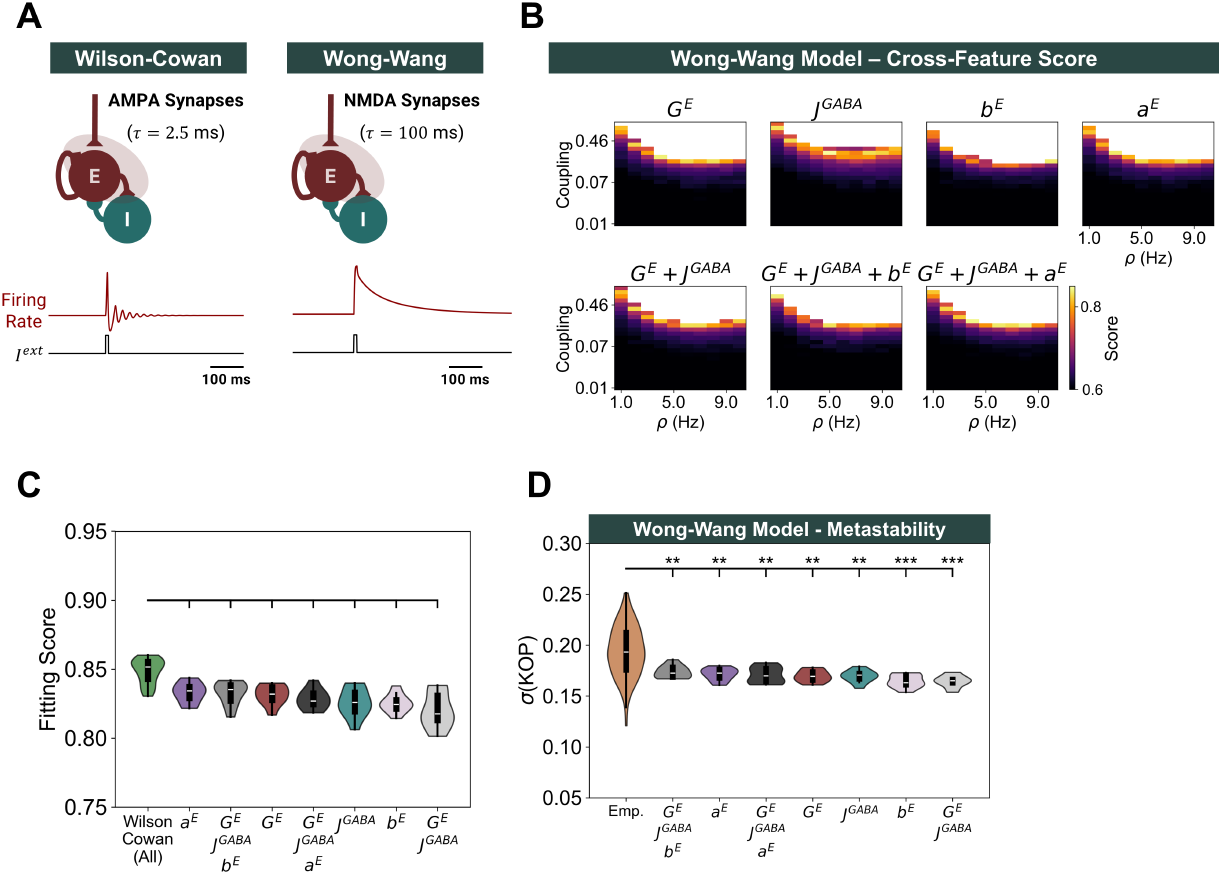
Effects of E-I Homeostasis in Models with Slow Local Dynamics. (A) Response of the Wilson-Cowan (Left) and Wong-Wang (Right) models to perturbations in external inputs. While the Wilson-Cowan model quickly returns to the equilibrium, exhibiting transient gamma oscillations, the Wong-Wang model takes longer to recover due to the slower dynamics of NMDA synapses. (B) Parameter spaces representing the cross-feature score of models based on Wong-Wang dynamics for different combinations of *C* and *ρ*. Blank spaces represent combinations of *C* and *ρ* for which the homeostatic set point is not valid (i.e. mean firing rates differ from *ρ* by more than 15% in at least one cortical area) (C) Comparison between cross-feature fitting scores at the optimal point of each mechanism of homeostasis and the Wilson-Cowan model with homeostasis of *G*^*E*^, *c*^*EI*^, *µ*^*E*^ and σ^*E*^. Distributions correspond to the result of 10 simulations with optimal *C* and *ρ*. Brackets indicate a significant difference, with p<0.05 from a Mann-Whitney U-test. All p-values were FDR-corrected. D) Comparison between metastability in empirical data and models at the optimal point (best cross-feature fitting score) for each mechanism of homeostasis. Values correspond to the results of 10 simulations with optimal *C* and *ρ*. Brackets represent a significant difference between models and empirical data from a Mann-Whitney U-test (^*^ p<0.05; ^**^ p<0.01; ^***^ p<0.001). All p-values were FDR-corrected.

With that in mind, we explore the performance of large-scale models based on Wong-Wang local dynamics by studying the dynamics of networks with different combinations of target firing rate (*ρ*) and global coupling (*C*). In line with the implementation of (50), we do not account for inter-areal conduction delays, due to their negligible impact on the slow NMDA dynamics. Furthermore, following (50), we consider a simulation valid if the firing rate is within 15% of the target, as opposed to the more stringent condition of 1% imposed on the Wilson-Cowan model. For each combination of parameters, we analyze how the model reproduces empirical FC and FCD by computing the cross-fitting scores. Our results indicate that the score depends on the interaction between *C* and *ρ*, similarly to the Wilson-Cowan model (Figure 5B). The optimal values of *C* and *ρ* and the respective scores can be consulted in Table S1. When comparing results at the optimal point for each model, our results show that there is no significant difference between the fitting scores of models with distinct mechanisms of homeostasis (Figure 5C). This is due to the reduced impact of E-I homeostasis on the local NDMA dynamics (Fig S28), when compared to its effect on the dynamics of faster gamma rhythms, dependent on AMPA receptors (Figures 1B and S1) (29). More importantly, the performance of any of the networks based on the Wong-Wang model is significantly lower than the performance of the Wilson-Cowan model with the homeostasis of excitation (*G*^*E*^), inhibition (*c*^*EI*^) and excitability slope (σ^*E*^) and threshold (*µ*^*E*^) (Figure 5C). In line with this result, even though we also found a strong relationship between metastability and the emergence of FC and FCD when using the Wong-Wang model (Fig S29), networks based on slow NMDA dynamics were not able to reach the levels of metastability observed in empirical data (Figure 5D).

That considered, our results suggest that the resting-state collective dynamics of the cortex, measured with BOLD fMRI, more strongly reflect the interaction between gamma rhythms generated by fast excitation and inhibition than the slower dynamics of NDMA receptors. Therefore, given the strong impact of E-I homeostasis on local gamma dynamics, the collective dynamics of the cortex at rest are strongly dependent on the mechanisms employed for the homeostasis of firing rates.

### Flexible Network Dynamics Emerge Independently of Noise

Synaptic transmission is kn, we propose that own to be noisy in brain networks (61). This variability can have a relevant role in spontaneous brain dynamics (61, 62), underlying phenomena such as stochastic resonance (24, 63, 64). In addition, systems with multiple weakly stable attractor states display noise-driven transitions between attractors, corresponding to multistable dynamics (20, 24), which can be confounded with metastability (65). In metastable systems, collective dynamics emerge as a consequence of the interaction between network topology and node dynamics, largely independently of noise. However, it is not yet clear if resting-state brain activity is more reflective of a metastable or multistable system (24, 65). For that reason, we explore the behavior of our network under different levels of noise (see Methods). Given that the model with all mechanisms of homeostasis better reproduces empirical dynamics, connectivity, and complexity, we take the optimal model with all forms of homeostasis (*C* = 3.59, *ρ* = 0.12) and simulate BOLD signals under varying levels of noise (see Methods). We then analyze model performance, collective dynamics, and FC complexity to assess how they are influenced by the variability of neuronal noise. Since we did not observe an interaction between noise and conduction delays (Fig S30), we present the average results across mean delays (Figure 6). Our results demonstrate that there is no significant effect of noise on model performance, collective dynamics, and FC complexity for lower noise levels. Conversely, as the variance of noise increases beyond these levels, model performance deteriorates. This effect is to be expected, given that the magnitude of noise becomes comparable to the intrinsic fluctuations in node dynamics (node activity is constrained between 0 and 1 by the sigmoid activation function). Furthermore, we observe a small peak in synchrony at levels of noise close to 0.1, which could suggest stochastic resonance (63), in line with studies showing noise-induced synchronization due to the structure of the human connectome (66). However, given that this peak does not correspond to the emergence of empirically relevant dynamics and there are no significant effects on the remaining network features, we do not explore this further.

**Fig. 6.**
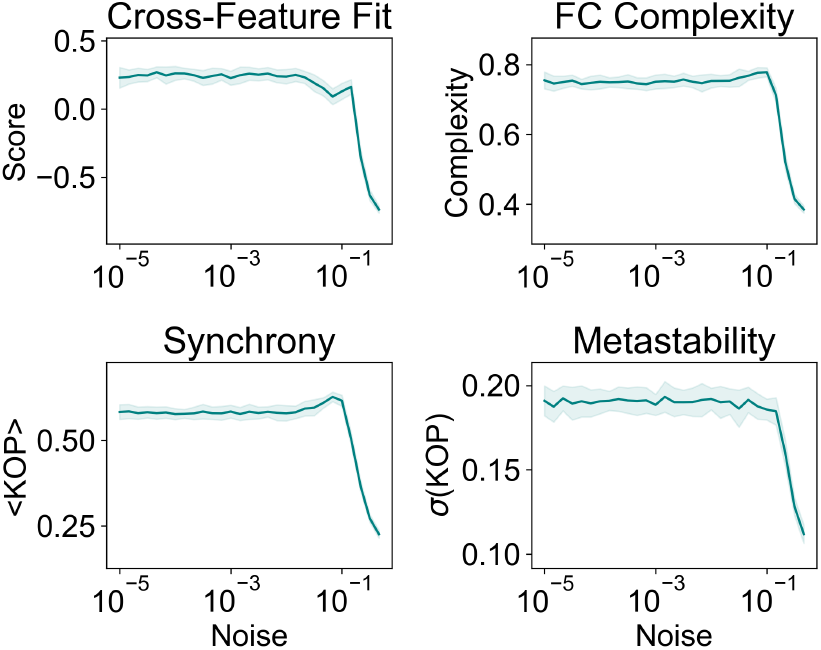
Changes in network features with the variance of noise. Values relate to models with *C* = 3.59 and *ρ* = 0.12 and all mean delays between 0 and 40 ms corresponding to valid network solutions. Solid lines represent the average and shaded areas the standard deviation across mean delays.

That said, given that model performance and dynamics are largely independent of noise, our results favor the emergence of metastable dynamics, as opposed to multistability (24, 65). For this reason, we consider that the collective dynamics of our models are most likely a product of the emergence of metastability, shaped by edge-of-bifurcation dynamics at the circuit level (regulated by E-I homeostasis) and the topology of inter-areal interactions (constrained by the structural connectome).

### Multiple Homeostatic Mechanisms Drive the Re-emergence of Functional Networks and Collective Dynamics after Structural Lesions

Our results establish the key role of local E-I balance and its homeostasis for the emergence of empirically valid large-scale networks and dynamics in cortical networks. E-I homeostasis can also be a significant driver of the reemergence of large-scale FC properties in the lesioned cortex (53–56). However, our earlier results (56) suggest that scaling of inhibition, the homeostatic mechanism most commonly implemented in large-scale models (48–52), is insufficient by itself for the recovery of metastable dynamics. Therefore, we investigate if the conjugation of multiple forms of homeostasis identified here can potentiate the re-emergence of not only FC but also network dynamics following structural lesions. To do this, we apply structural lesions in the optimal model: including homeostasis of excitation (*G*^*E*^), inhibition (*c*^*EI*^) and intrinsic excitability (*µ*^*E*^ and σ^*E*^) with *C* = 3.59, *ρ* = 0.12 and mean delay = 40 ms, corresponding to a conduction velocity (3.9m/s) close to the values reported in previous studies (67, 68). We extract activity before the lesion (healthy), after the lesion (acute), and after node parameters are adapted to restore local E-I balance (chronic). For more details on the simulation protocol and analysis (56) refer to the Methods and SI Methods. All p-values presented in this section relate to the result of the Wilcoxon-Ranked sum test and are FDR-corrected.

Starting with FC, we measure the Euclidean distance between healthy FC and FC in the acute and chronic periods (Figure 7A). We observed that, even though FC patterns are disrupted in the acute period (10.751±4.900) significant recovery toward pre-lesion FC configurations (5.800±1.573; p<0.001) occurs in the chronic period. Consistent with stroke literature (69), the model displays an acute decrease in their correlation between structural and functional networks (−10.97±13.34%; p<0.001) (Figure 7B). However, this disruption was recovered in the chronic period through the action of E-I homeostasis (1.33±6.10%; p=0.059). In addition, we observe an acute reduction of modularity (−6.00±10.39%; p<0.001) (Figure 7C), consistent with that observed in stroke patients (70, 71). In line with previous results (56, 71), the recovery of E-I balance drives the re-emergence of modularity toward pre-lesion levels in the chronic period (2.41±12.44%; p=0.257). Therefore, our model can replicate the effects of lesions on FC and its subsequent recovery through focal E-I homeostasis. We further explore the effects of lesions on functional complexity (Figure 7D), revealing a minimal decrease in the acute period (−1.86±6.21%; p=0.046) and a return to baseline after recovery of E-I balance (−0.62±3.81%; p=0.240). Therefore, our results suggest that, as opposed to the FC-SC relation and FC modularity, FC complexity may be minimally disrupted by lesioning the connectome.

**Fig. 7.**
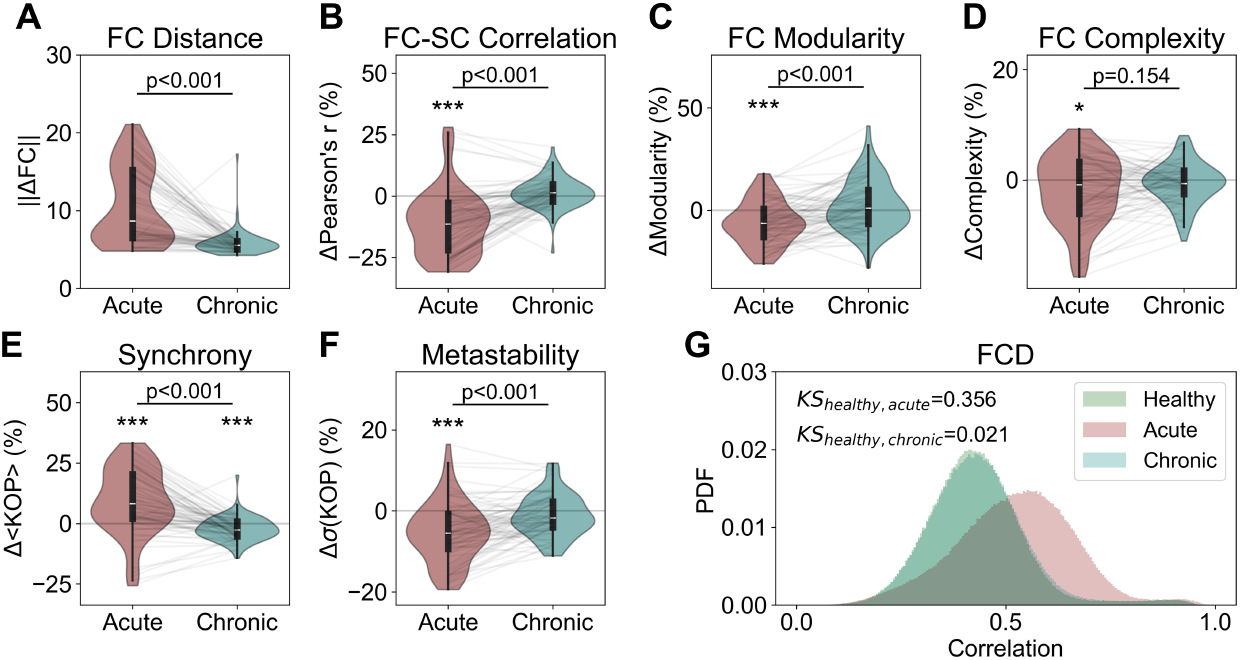
Disruption and recovery of network properties after focal lesion in models with homeostasis of excitation (*G*^*E*^), inhibition (*c*^*EI*^) and intrinsic excitability (*µ*^*E*^ + σ^*E*^) **(A)** Distance from baseline of FC matrices in acute (post-lesion) and chronic (post-lesion with homeostatic recovery of E-I balance) simulations (B) Change from baseline (in percentage) of FC-SC correlation in the acute and chronic simulations (C) Same as B, for FC Modularity (D) Same as B, for FC complexity (E) Same as B, for synchrony (F) Same as B, for metastability (G) Distributions of FCD correlations in all stages of our lesion simulations. In addition, we present the Kolmogorov-Smirnov distance between distributions at baseline and acute/chronic simulations. In all plots, p-values show the significance of the Wilcoxon Ranked-sum test (^*^ p<0.05; ^**^ p<0.01; ^***^ p<0.001). All p-values were FDR-corrected.

Beyond reproducing the effects of stroke, and subsequent recovery, on functional connectivity (53, 56, 69, 70, 72), we explore how lesions disrupt network dynamics. Starting with synchrony, our results indicate that the system tends toward hypersynchrony in the acute period (9.31±13.44%; p<0.001) (Figure 7E), in line with literature on the effects of brain injury (73). More importantly, synchrony is recovered toward pre-lesion levels in the chronic period (p<0.001), although a small, but significant, difference from baseline remains (−2.19±4.88%; p<0.001). Regarding metastability, we observe a significant decrease post-lesion (−5.30±7.32%; p<0.01) (Figure 7F), followed by a significant recovery (p<0.001) to pre-lesion levels in the chronic period (−1.08±5.17%; p=0.060). Finally, given the relationship between metastability and FCD (22, 24), we analyze FCD distributions in the simulated Healthy, Acute, and Chronic periods. In line with (56), we observe an acute disruption, with the mean FCD correlation values shifting from 0.429 in the Healthy period to 0.522 post-lesion (KS distance=0.356) (Figure 7F). Importantly the combination of multiple forms of homeostasis can drive the recovery of FCD distributions to pre-lesion levels (mean correlation=0.434; KS distance=0.021), as opposed to the plasticity of inhibition alone, as attested by our previous results (56).

Finally, we have also performed the same analysis steps in models with the *G*^*E*^ or *G*^*E*^ + *c*^*EI*^ homeostasis, showing that such models are either unable to reproduce the studied post-stroke disruptions in FC and network dynamics, or such disruptions were not recovered through E-I homeostasis (Figures S31-32).

In summary, the conjugation of multiple mechanisms of homeostasis not only supports metastable dynamics and empirical FC networks but can also drive their re-emergence following lesions to the connectome, which is not the case in models with homeostasis of excitation or inhibition only. Therefore, the multi-factor homeostasis of E-I balance can also be considered a robust means of self-organization in large-scale cortical networks.

## Discussion

Building on the observation that multiple mechanisms of E-I homeostasis are essential for the regulation of local circuit dynamics (47), we extend this hypothesis to large-scale networks, exploring how distinct homeostatic mechanisms impact the global dynamics and functional networks of the human cortex. Our results demonstrate that the homeostatic regulation of local E-I balance supports the emergence of resting-state FC and FC dynamics, particularly when involving forms of homeostasis that modulate the distance to the bifurcation between damped and sustained oscillations (29) (Figures 1B and S1). In addition, we show that the resulting heterogeneity in the distance to the bifurcation across cortical areas supports the emergence of empirical FC and FCD by compensating for heterogeneities in the magnitude of input fluctuations. We further demonstrate that the optimal working points of models generally correspond to moderate levels of synchrony and heightened metastability, confirming that cortical networks might operate in a regime of enhanced metastability. More importantly, empirical levels of synchrony and metastability can only be replicated by models that incorporate multiple mechanisms of homeostasis. Furthermore, we study the complexity of large-scale functional interactions, stressing that it not only depends on the topology of structural networks (16–18) but also on local E-I balance. Importantly, empirical levels of complexity were more closely approximated by the model with all forms of homeostasis enabled. Altogether, we confirm the hypothesis that the benefits of the interaction between multiple forms of E-I homeostasis extend beyond the local circuit level (40–43, 47), contributing to the regulation of large-scale dynamics toward a regime of maximal metastability and complexity, which underlies the FC networks observed in the resting-state cortex. Finally, we also demonstrate that the combined action of multiple homeostatic mechanisms can drive the re-emergence of not only FC but also network dynamics following focal lesions to the connectome (56), evidencing the robustness of E-I homeostasis as a self-organization process in cortical networks. Importantly, our study provides a framework to tie several relevant features of cortical organization and dynamics across spatial scales. More specifically, the regulation of E-I balance in cortical neurons allows for the regulation of edge-of-bifurcation dynamics at the circuit level which, in turn, underlies the self-organization of spontaneous large-scale dynamics toward a regime of high metastability, underlying the resting-state activity of the human cortex.

To our knowledge, we are the first to explore the combined action of diverse mechanisms of E-I homeostasis in large-scale models, given that most studies focus on the homeostasis of inhibition (48–52, 56). At the circuit level, the synergy between the synaptic scaling of excitation and inhibition and the plasticity of intrinsic excitability are essential for the maintenance of not only stable firing rates but also edge-of-bifurcation dynamics (i.e. criticality) (42, 47, 74). Our results add to this perspective by suggesting that these benefits extend to larger scales, supporting network dynamics and FC networks. More specifically, our results show that networks with synaptic scaling of excitation and inhibition reach heightened levels of global metastability, characteristic of the cortex at rest (6, 10, 19, 22, 24). Regarding the plasticity of intrinsic excitability, it is not yet clear from experimental data if it acts through the modulation of firing thresholds only (*µ*^*E*^) (40) or the concerted adaptation of both the threshold and slope of neuronal excitability (*µ*^*E*^ +σ^*E*^) (45, 46, 75). Here, we suggest that the second option is more plausible given its contribution to ensuring the moderate levels of synchrony observed in empirical data, while still allowing for highly metastable dynamics. In this line, it should be highlighted that recent results suggest that, while the plasticity of intrinsic excitability might not be mobilized in the adult cortex, being instead pivotal during development (46). However, it has also been suggested that homeostasis of intrinsic excitability can also be mobilized following substantial perturbations that cannot be compensated for by the homeostasis of excitation and inhibition (41). This aligns with evidence from the post-stroke brain, where processes typical of the developmental phases, such as neurogenesis, are also recruited for recovery (76). Therefore, we argue that the diverse mechanisms of homeostasis have specific contributions to cortical dynamics at both the circuit and global scales which may play a role in both the healthy and diseased brain.

In light of our results, it is relevant to reflect on how exactly local E-I balance contributes to the large-scale dynamics of the cortex. Here, we focus on the edge-of-bifurcation theory for cortical dynamics (6, 19–24). Under this perspective, local circuits at the edge-of-bifurcation can undergo spontaneous phase transitions, which, when coincident across cortical areas, can self-reinforce, leading to the emergence of the collective behaviors typical of the cortex (19). In our model, these collective dynamics are strongly regulated by the maintenance of stable mean firing rates (*ρ*) through E-I homeostasis (29). For example, increases in global coupling (*C*), which may push node dynamics beyond the bifurcation, can be compensated by lowering *ρ*. For this reason, model performance and dynamics are heavily dependent on the interaction between *C* and *ρ*. On the other hand, the heterogeneity in the in-degrees of the connectome not only leads to stronger inputs in highly connected nodes, but also higher input fluctuations (Fig S17). Therefore, if all cortical areas were to be equally distant from the bifurcation, the highly connected ones would be more likely to engage in sustained oscillations due to strong input fluctuations that push them beyond the bifurcation. However, this effect can be compensated for by forms of homeostasis (i.e. excitation and excitability slopes) that maintain stable firing rates by scaling the distance to the bifurcation (Figs 1B and S1) (29), poising highly connected nodes farther away from the bifurcation point. This adjustment is critical to ensure that cortical areas exhibit edge-of-bifurcation dynamics regardless of the variability of their inputs. More so, this aligns with previous studies showing that distance to criticality varies across cortical areas (77) and that global metastable dynamics are better reproduced in models with heterogeneous distances to the bifurcation (22). More so, recent computational studies investigating brain dynamics during different states of consciousness are in line with our findings that strongly connected areas are poised further away from the bifurcation, at least in the awake cortex (78). Importantly, we demonstrate that this heterogeneity, related to differences in connectivity, can emerge naturally from the maintenance of stable population firing rates via multiple forms of homeostasis. In this line, we suggest that future studies investigate how the heterogeneity in distance-to-bifurcation across cortical areas (22, 77, 78) relates to their structural connectivity and microcircuitry (79) or to the interaction between E-I homeostasis and states of reduced consciousness (78).

Our results indicate that empirical FC dynamics emerge in a state of maximal metastability (Fig S23), confirming that FCD reflects the exploration of the state space of network configurations potentiated by metastable brain dynamics (9, 10, 19, 22, 24). Furthermore, it has been argued that metastability is generally derived from abstract Hopf-bifurcation models, with no direct relationship to physiological processes (24). However, we point out that previous studies, although not quantifying metastability directly, have demonstrated the emergence of large-scale wave patterns consistent with metastable dynamics in models similar to ours (80). Here, we demonstrate that these theoretical concepts are directly applicable in a model of the neocortex where the generation of oscillations is based on E-I interactions and bifurcation control implemented by E-I homeostasis. This confirms that the concept of metastability is compatible with both the architecture of the human cortex and its processes of self-organization. Importantly, our results also show that the collective dynamics of the human cortex most likely reflect metastability and not multistability (65), making a further argument in favor of the hypothesis that metastability is one of the key dynamical features of large-scale cortical networks (6, 9, 10, 24).

Our analysis further shows a relationship between metastable dynamics and the complexity of functional interactions. In complex systems, the interactions between units are dictated by both the structural scaffold and circuit dynamics (81). In the cortex, the architecture of the connectome plays an important role in the emergence of complex spontaneous activity (16, 17) and its graph properties are required to support the complexity observed in empirical FC networks (18). By exploring the relationship between complexity and metastability, we demonstrate that FC complexity not only reflects the topology of the connectome, but is also tightly related to metastable dynamics (Figures 4B and S23), suggesting it reflects a balanced exploration of network states (9, 10). When averaged over long time scales, this exploration leads to a distribution of FC weights that is closer to uniformity, revealing a dynamic balance between integration and segregation. For this reason, we propose that the complexity of functional interactions can be another signature of local E-I balance, which allows for the emergence of metastable collective dynamics (6, 19, 21, 22) through the regulation of local edge-of-bifurcation dynamics (29). That said, networks with homeostasis of excitation (*G*^*E*^) or excitation and inhibition (*G*^*E*^ + *c*^*EI*^), associated with high metastability (Figure 3B), display levels of complexity considerably lower than found in empirical data (Figure 4B). However, such models also show overly synchronized dynamics (Figure 3A). Conversely, when the plasticity of intrinsic excitability (*µ*^*E*^ + σ^*E*^) is introduced, synchrony is maintained at empirical levels and complexity enhanced. Therefore, we propose that the plasticity of intrinsic excitability has an important role in maintaining complex interactions by avoiding hyper-synchrony while still allowing for highly metastable dynamics.

We not only demonstrate the association between local E-I balance and collective metastable dynamics, but also the relevant role of gamma oscillations as a connecting mechanism between the local and global scales. To do this, we compare the dynamics in models with local circuits based on either fast AMPA (i.e. Wilson-Cowan) or slow NMDA (i.e. Wong-Wang) synapses, both equipped with local E-I homeostasis. The first relevant conclusion from this exploration is that, in models with slow excitation, as long as mean firing rates are maintained locally, collective dynamics are not heavily shaped by which homeostatic mechanisms are in place (Figure 5C). We suggest that this occurs because the Wong-Wang model is not capable of generating oscillations and, thus, the bifurcation modulation of some homeostatic mechanisms, discussed in (29), is not observed in the Wong-Wang model (Fig S28). Conversely, in the Wilson-Cowan model, this effect is essential to compensate for heterogeneities in input fluctuations and, therefore, the combination of local homeostatic mechanisms strongly impact global dynamics. For this reason, the way multiple homeostatic mechanisms shape collective dynamics is strongly tied to the regulation of edge-of-bifurcation oscillations and not just the maintenance of firing rates. More importantly, our results also show that models with slow excitation cannot reach the metastability levels observed in empirical resting-state data (Figure 5D). Here, we propose that the relevance of gamma rhythms for the collective dynamics of our model has two main explanations. The first, and more obvious, is that we base our measurements on the dynamics of BOLD signals, which have been associated with fluctuations in gamma power and synchrony in both experimental (82–84) and computational (52, 63, 85) studies. Therefore, it is natural that models that generate gamma rhythms would be more equipped to reproduce the metastable dynamics of BOLD signals. However, based on previous theories for the generation of spontaneous cortical dynamics (2, 6, 19, 30), we advance a further explanation. At the local level, E-I balance is maintained by regulating not only excitation but also the fast GABAergic synapses from fast-spiking interneurons (35, 41, 47, 86). Since gamma oscillations are also generated by the recurrent loop between pyramidal neurons and fast-spiking interneurons (59, 60, 87, 88), E-I homeostasis should have strong effects on gamma oscillations. Previously, we have proposed that one of such effects is the modulation of edge-of-bifurcation oscillations (29), in line with empirical studies showing the role of multiple homeostatic mechanisms in the regulation of both firing rates and circuit-level criticality (47). That said, given that edge-of-bifurcation oscillations have been shown to support the emergence of metastable activity patterns in the cortex (6, 19, 22, 80), our results support the idea that gamma oscillations can be a necessary binding factor between local E-I balance and global metastable dynamics (19, 30). For this reason, we suggest that gamma rhythms should necessarily be accounted for in large-scale models built to study the relationship between local and global dynamics in cortical networks. That said, since the combination of multiple homeostatic mechanisms is pivotal to ensure that gamma frequencies remain within physiological bounds (29), we suggest that large-scale modeling studies should also go beyond the plasticity of inhibition as the sole mechanism of homeostasis when studying E-I balance.

To further validate our model, we also examine the role of E-I homeostasis in the self-organization of the cortex following focal lesions, following the framework of (56). We replicate disruptions in FC properties such as SC-FC coupling (69) and modularity (71), characteristic of the post-stroke brain, which are then recovered through cortical re-organization, driven by standard mechanisms of homeostatic plasticity, in line with previous results (56). However, the model in (56) was equipped with the plasticity of inhibition only, which proved insufficient for the recovery of metastable dynamics and FCD. Here, we demonstrate that networks with multiple mechanisms of E-I homeostasis (excitatory and inhibitory synapses and intrinsic excitability) can not only replicate the disruptions and subsequent recovery of macroscale FC properties, but also return metastability and FCD to pre-lesion levels through the restoration of E-I balance (Figure 6). That said, not all mechanisms of homeostasis may be necessary for the recovery of network dynamics. Therefore, we apply the same lesion protocol in networks with either *G*^*E*^ or *G*^*E*^ + *c*^*EI*^ homeostasis (Figs S31-32). Critically, those are either unable to recover network dynamics or do not reproduce empirical post-stroke deficits in FC (69–71). Therefore, while the homeostasis of inhibition might strongly contribute to the recovery of FC networks, as demonstrated in (56), the plasticity of excitation and intrinsic excitability is critical for ensuring that network dynamics can be recovered following damage to structural networks.

Our results show that, by imposing stronger constraints on the local circuitry of cortical networks, E-I homeostasis potentiates the flexible exploration of macroscale network states. While this effect appears paradoxical, it is in line with the concept of “constraints that deconstrain” (89). This idea stems from the work of Kirschner and Gerhart (90), who suggest that some constraining mechanisms are preserved through evolution because they deconstrain other processes that are advantageous for the organisms. In neuroscience, this concept has been applied to explain how the layered architecture of brain networks (91, 92) ensures both robust motor control and high adaptability in behavior (89). We propose that the tight control of E-I balance by multiple mechanisms of homeostasis is another example of a “constraint that deconstrains”, by supporting the emergence of metastability which endows the cortex with the flexibility needed to explore its state-space during rest (6–10, 19).

There are, however, some limitations to point out in our model. The first is that we do not explore homeostatic plasticity at the level of inhibitory neurons. Recent studies have shown that the plasticity of excitatory connections (28) or intrinsic excitability (42) in parvalbumin-positive (PV) interneurons can also contribute to E-I homeostasis. Therefore, future models should consider the implementation of E-I homeostasis in inhibitory neurons, delving into the interaction between the homeostatic set-points of excitatory and inhibitory populations. Another limitation relates to the fact that, while synaptic scaling of excitation is ubiquitous across cortical layers (40, 43, 44, 46, 93), other forms of plasticity, such as the scaling of inhibition, might be absent from deeper layers (40, 43, 46, 93, 94). Therefore, we suggest that E-I homeostasis should be explored in large-scale models accounting for the laminar structure of the neocortex (95, 96). Furthermore, we implement a homogeneous target firing rate across the network, in line with previous modeling studies (48– 51). While *in vivo* studies of E-I homeostasis are mostly based on the activity of visual or somatosensory cortices (28, 93), different cortical areas might be regulated toward area-specific set points, possibly following hierarchical gradients of cortical organization (79). For this reason, future models should explore the possibility of not only implementing heterogeneous target firing rates, but also the interaction between cortical hierarchy, structural heterogeneities (97) and the homeostasis of E-I balance. In addition, our model can also be used in conjunction with a more data-driven approach, as employed by (97), to optimize model parameters, investigating how this interacts with local homeostasis of E-I balance.

Finally, we tune the Wilson-Cowan model to generate gamma oscillations, which have been related to fluctuations in BOLD signals (21, 52, 63, 82) and transient rhythms across frequency bands (15). While there is empirical evidence for gamma resonance through the recurrent interactions between PY and PV neurons (58, 60), some studies suggest that cortical networks also generate alpha oscillations in deeper layers (98, 99). Therefore, the extension of our framework to the laminar architecture of the cortex (95, 96) might also provide new insight into how E-I balance interacts with different cortical rhythms, fast and slow inhibition (29) and how it shapes cross-frequency interactions.

To conclude, our results demonstrate the critical role of firing-rate homeostasis, which regulates edge-of-bifurcation dynamics at the circuit level, as a key piece underlying the self-organization of resting-state cortical activity toward a regime of heightened metastability where resting-state networks emerge. More importantly, we show that this reflects the joint action of multiple homeostatic mechanisms employed to maintain E-I balance, suggesting that their influence extends beyond the neuronal and circuit level, underlying the rich spontaneous dynamics of cortical networks.

## Materials and Methods

### Structural Connectivity Data

To approximate the anatomical connections between brain areas, we use a normative connectome from 32 healthy participants (mean age 31.5 years old ±8.6, 14 females) generated as part of the Human Connectome Project (HCP - https://www.humanconnectome.org). In this work, we focus on the cortical regions of the Anatomic Automatic Labeling (AAL) atlas, yielding a 78×78 matrix. Tract lengths between brain regions were also extracted to be used in the computation of conduction delays between network nodes. For more detailed information refer to (52).

### BOLD fMRI Data

To analyze network dynamics and FC properties in empirical data, we use resting-state blood-oxygenation-level-dependent (BOLD) time series from 99 healthy unrelated subjects from the HCP dataset. Data was processed using the pipeline described in (52), yielding 99 time series of size 78 areas x 1200 TR, roughly corresponding to 14.5 minutes of data for each subject (TR=0.72 s).

### Large-Scale Model

To model the activity of individual cortical regions, comprising excitatory (*r*^*E*^) and inhibitory (*r*^*I*^) neural masses, we make use of the Wilson-Cowan model (100, 101):

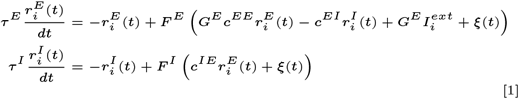

Where *c*^*EE*^ represents the recurrent excitatory coupling, *c*^*EI*^ the coupling from inhibitory to excitatory populations and *c*^*IE*^ the excitatory coupling unto inhibitory populations. *G*^*E*^ is a parameter introduced in (29), allowing for the scaling of all excitatory inputs to the excitatory neural mass (i.e. recurrent excitation and *I*^*ext*^). The default values of each parameter can be consulted in Table S1. Unless stated otherwise, *ξ*(*t*) represents Gaussian noise with mean 0 and variance 0.01. Here, 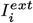 describes the incoming input to each node *i* from the rest of the network and can be written as:

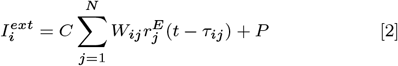

where *C* is a scaling factor referred to as global coupling, *W*_*ij*_ is the structural connectivity matrix and *τ*_*ij*_ represents the conduction delay between nodes *i* and *j*. Here, similarly to (51, 52, 56), *P* represents a background input imparted on the nodes and is set at 0.31 so that the default uncoupled node is close to the Hopf bifurcation. The time constants *τ* ^*E*^ and *τ* ^*I*^ were tuned so that the uncoupled node oscillates at ∼40 Hz. Furthermore, we define *F*^*E/I*^ (*x*) as the input-output function of the excitatory and inhibitory populations, respectively, using the following equation:

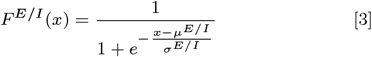

### Excitatory-Inhibitory Homeostasis

As opposed to the common approach to E-I homeostasis (48–52, 56), we do not implement dynamical homeostasis. Instead, we follow the framework developed in (29) to mathematically estimate the parameter values that ensure the maintenance of mean excitatory activity at a target firing rate (*ρ*) for different levels of input (*I*^*ext*^) (28, 93), in line with the governing rules of E-I homeostasis (41). That said, to compute the homeostatic parameters of nodes embedded in a network, it is necessary to provide an estimate of *I*^*ext*^ for each node. To do this, we rely on the fact that E-I homeostasis works in timescales much slower than neural dynamics (41, 42, 47), ensuring timescale separation. Therefore, to compute the homeostatic parameters, we consider the average external input received by each node at the homeostatic set point, when the average activity across the network corresponds to *ρ*, leading to the following expression:

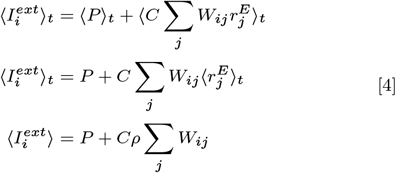

Here, we explore the behavior of models with the following distinct mechanisms of E-I homeostasis, based on experimental results (39–41, 43, 45–47) and described in detail in (29):

- *G*^*E*^ homeostasis: synaptic scaling of excitatory synapses in PY neurons.
- *c*^*EI*^ homeostasis: synaptic scaling of inhibitory synapses in PY neurons.
- *µ*^*E*^ homeostasis: plasticity of the intrinsic excitability of PY neurons, modulating the “firing threshold” (i.e. *µ*^*E*^) of *F*^*E*^(*x*).
- *µ*^*E*^ + σ^*E*^ homeostasis: plasticity of the intrinsic excitability of PY neurons, modulating both the “firing threshold” (*µ*^*E*^) and “slope” (σ^*E*^) of *F*^*E*^(*x*) in a synergistic manner.
- *G*^*E*^ + *c*^*EI*^ homeostasis: synaptic scaling of excitatory and inhibitory synapses in PY neurons.
- *G*^*E*^ +*c*^*EI*^ +*µ*^*E*^ homeostasis: synaptic scaling of excitatory and inhibitory synapses and intrinsic excitability of PY neurons, implemented at the level of *µ*^*E*^.
- *G*^*E*^ +*c*^*EI*^ +*µ*^*E*^ +σ^*E*^ homeostasis: synaptic scaling of excitatory and inhibitory synapses and intrinsic excitability of PY neurons, implemented at the level of *µ*^*E*^ and σ^*E*^.

The equations to compute the set-point parameters corresponding to each form of homeostasis can be consulted in SI Methods.

### Hemodynamic Model

From the raw activity of the excitatory populations, we extract simulated BOLD signals by using a forward hemodynamic model (102, 103), described in detail in (56). This model represents the coupling between model activity and blood vessel diameter, which affects blood flow, blood volume, and deoxyhemoglobin content. After passing model activity through the hemodynamic model, the output is downsampled to a sampling period of 0.72s to align simulated signals with empirical data.

### Network Validity

For each type of homeostasis and combination of hyper-parameters (*C, ρ*, and mean delay), we evaluate if the homeostatic set point - i.e. a mean firing rate of *ρ* for all nodes - can be maintained across the network. To do this, we run 15-second simulations and compute the mean firing rate of each node in the last 10 seconds of the simulation. We consider the given network fixed point to be invalid if there is any node in the network for which the mean firing rate differs from *ρ* by more than 1% (i.e.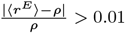). The dynamical behavior of invalid models is examined in detail in SI Text. While it is possible to evaluate the stability of a network analytically (see (104)), the characteristic polynomial of our model (from which the eigenvalues can be extracted) does not have a closed-form solution due to the highly non-linear nature of the Wilson-Cowan equations. For this reason, we base our exploration of model validity on the analysis of the numerical solution of the system.

### Model Optimization

We perform model optimization by exploring all combinations of the global coupling (*C*), mean delay, and target firing rate (*ρ*) within the following ranges of variation: *C* : [0, 10], Mean Delay: [0, 40] ms and *ρ*: [0.05, 0.2]. Within these ranges, we selected 19 logarithmically spaced values for *C*, 16 values for *ρ* in steps of 0.01, and 41 mean delays in steps of 1 ms. For each simulation, we initialize the model with the homeostatic parameters corresponding to the combination of *C* and *ρ* and record model activity for 30 minutes. To evaluate model performance against empirical data, we use the following properties of FC.

- **Static FC**: 78×78 matrix of correlations between BOLD time series across all network nodes. Modeled FC matrices were compared with group-averaged empirical FC by computing the correlation coefficient (*r*_*FC*_) and mean squared error (*MSE*_*FC*_) between their upper triangular elements.
- **FC Dynamics (FCD)**: matrix of correlations between the upper-triangular part of FC matrices computed in windows of 80 samples with 80% overlap (22, 105). Model results are compared to empirical data by computing the Kolmogorov-Smirnov distance (*KS*_*F CD*_) between the distributions of values in the respective FCD matrices.

BOLD signals are filtered between 0.008 and 0.08 Hz before computing FC and FCD. In addition, we compute a cross-feature fitting score as 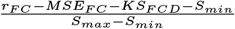 to evaluate the performance of models in simultaneously reproducing static and dynamic properties of empirical FC (52, 56). *S*_*max*_ = 1 represents the maximum value of *r*_*FC*_ − *MSE*_*FC*_ − *KS*_*F CD*_ (1-0-0) while *S*_*min*_ = − 4.0024 represents its minimum (−1 - 2.0024 - 1). The value of 2.0024 represents the maximum possible MSE between an FC matrix and our empirical FC matrix, considering that FC can only vary between -1 and 1. With this expression, we obtain a cross-feature fitting score normalized between 0 and 1, indicating how well models represent FC and FCD.

All simulations were performed by solving the system’s equations numerically using the Euler method with an integration time step of 0.2 ms.

### Synchrony and Metastability

To evaluate the magnitude of synchrony and the degree of dynamic switching between network states (i.e. metastability), we compute the Kuramoto Order Parameter (KOP) (106, 107) of BOLD timeseries, filtered between 0.008 and Hz. More specifically, the KOP (*R*(*t*) in Eq 5) represents the degree of phase alignment within a set of coupled oscillators at a given point in time and can be calculated as:

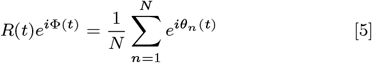

where *θ*_*n*_(*t*) represents the instantaneous Hilbert phase of node *n* at time *t*. Synchrony and metastability are defined, respectively, as the mean and standard deviation of *R*(*t*) over time.

### Functional Complexity

Previous studies have demonstrated that the architecture of structural networks has a significant impact on the complexity of the functional interactions between nodes (16–18). To explore this, we measure the complexity of simulated FC matrices using the method introduced in (18). In short, we quantify complexity by measuring how strongly the distribution of FC weights deviates from an equivalent uniform distribution using the following formula:

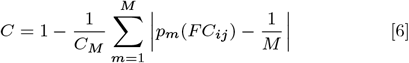

where *M* represents the number of bins used to evaluate the FC distribution. The term 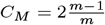 is added for normalization, corresponding to the deviation o^*m*^f a Dirac-delta function from the uniform distribution. Therefore, FC complexity is 0 when all entries of the FC matrix are the same and 1 when they are uniformly distributed. While there are other methods to evaluate the complexity of a matrix, (18) demonstrate that their formulation is more robust to changes in bin sizes, compared to other methods quantifying the entropy or variance of FC matrices. Here, we use a bin size of 0.05.

### Replication of Analysis Using the Wong-Wang Model

To investigate the effects of E-I homeostasis on slow local dynamics we employ the reduced Wong-Wang model of cortical populations (50). In this model, instead of simulating fast AMPA synapses, excitatory synapses have a considerably slower decay time (100 ms), consistent with the timescale of NMDA synapses. The model dynamics are described as follows:

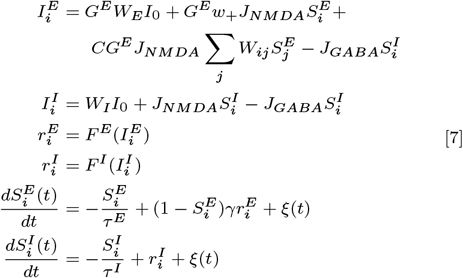

where 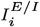 represents the total input to excitatory or inhibitory neuronal population *i*, 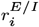 its firing rate and 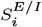 an average synaptic gating variabl^*i*^e. In this case, we hav^*i*^e also added the parameter *G*^*E*^, which allows for the simultaneous modulation of all excitatory synapses unto the excitatory neuronal population. The function *ξ*(*t*) represents additive Gaussian noise with mean 0 and variance 0.01 (nA). *H*^*E/I*^ (*t*) represents the input-output function of neuronal populations and is written as:

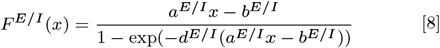

The parameter values, adapted from (50) and their physiological meaning are described in detail in Table S3. Importantly, this model does not consider the dynamics of AMPA synapses and *τ* ^*E*^ = 100*ms* (the time constant of excitatory synapses) represents the slow decay time of NMDA (50). Therefore, in line with (50) and given the focus of the Wong-Wang model on slower dynamics, inter-areal conduction delays are neglected in this model.

Similarly to the Wilson-Cowan model, we derived the equations allowing for the mathematical estimation of the steady-state value of each model parameter corresponding to the different empirically-based mechanisms of homeostasis. More specifically, we investigate the following forms of E-I homeostasis:

- *G*^*E*^ homeostasis: synaptic scaling of excitatory synapses in PY neurons.
- *J*_*GABA*_ homeostasis: synaptic scaling of inhibitory synapses in PY neurons.
- *b*^*E*^ homeostasis: plasticity of the intrinsic excitability of PY neurons, modulating the “firing threshold” (i.e. *b*^*E*^) of *F*^*E*^(*x*).
- *a*^*E*^ homeostasis: plasticity of the intrinsic excitability of PY neurons, the “slope” of (σ^*E*^) of *F*^*E*^(*x*) in a synergistic manner. In this case, since changing *a*^*E*^ has a minimal impact on the value of *F*^*E*^(*x*) at 0, we employ this form of homeostasis to simulate the plasticity of intrinsic excitability described in (45, 46, 75). For more details, refer to Fig S9.
- *G*^*E*^ + *J*_*GABA*_ homeostasis: synaptic scaling of excitatory and inhibitory synapses in PY neurons.
- *G*^*E*^ + *J*_*GABA*_ + *b*^*E*^ homeostasis: synaptic scaling of excitatory and inhibitory synapses and intrinsic excitability of PY neurons, implemented at the level of *b*^*E*^.
- *G*^*E*^ +*J*_*GABA*_ +*a*^*E*^ homeostasis: synaptic scaling of excitatory and inhibitory synapses and intrinsic excitability of PY neurons, implemented at the level of *a*^*E*^.

For more detail on the derivation of the mathematical expressions and the effects of E-I homeostasis at the circuit level, refer to the Supplementary Methods.

Similarly to the procedure employed for the Wilson-Cowan model, we perform model optimization by running one simulation for each combination of the global coupling (*C*) and target firing rate (*ρ*) within the following ranges of variation: *C* : [0, 1] and *ρ*: [1, 10] Hz. Within these ranges, we selected 25 logarithmically spaced values for *C* and 10 values for *ρ*. For each simulation, we initialize the model with the homeostatic parameters corresponding to the combination of *C* and *ρ* and record model activity for 30 minutes. In (50), the target firing rate is fixed at 3.0631 Hz the model is considered to have reached the homeostatic fixed point when the mean firing rate is between 2.63–3.55 Hz, corresponding to an interval of ∼15% around the target. Therefore, here we consider a simulation valid when the error between mean firing rates and the target is smaller than 15% for all nodes in the network.

Finally, we detect the optimal combination of *C* and *ρ* maximizing the fitting score for each form of homeostasis. To be able to statistically compare models, we run 10 simulations with each combination of optimal parameters and compute the respective cross-feature fitting score and metastability.

### Effects of Noise

To explore the impact of noise in our model, we run simulations of the network with homeostasis of *G*^*E*^, *c*^*EI*^ and *µ*^*E*^ + σ^*E*^ with optimal parameters (*C* = 3.59, *ρ* = 0.12) and across all mean delays yielding valid simulations. For each simulation, we vary the variance of Gaussian noise (*ξ*(*t*)), corresponding to 29 logarithmically spaced values between 10^−5^ and 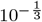. We then analyze model performance, synchrony, metastability, and functional complexity across all models with varying levels of noise.

### Lesion Simulation Protocol

Following the framework applied in (56), we investigate the effects of focal lesions in model dynamics and functional interactions by following the following protocol. Using the model with all mechanisms of homeostasis at the optimal point (*C* = 3.59; *ρ* = 0.12), we compute the homeostatic parameters by using the methodology from (29) (SI Methods). From the balanced model, we extract 30 minutes of BOLD signal corresponding to activity in the pre-lesion (i.e. healthy) period. Then, we apply a structural lesion by removing all connections to and from a chosen node and extract 30 minutes of activity corresponding to the early acute period of stroke recovery. We then compute the new homeostatic parameters, allowing for the restoration of E-I balance in the lesioned connectome, using the pre-lesion homeostatic values as 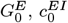 and 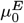 (SI Methods for the equations). From the re-balanced model, we extract 30 minutes of model activity corresponding to the chronic period. We follow this protocol for all possible single-node lesions and extract the following metrics from the healthy, acute, and chronic simulations: difference of FC from baseline, FC-SC correlation, FC modularity, FC complexity, Synchrony, Metastability, and FCD distributions. For a more detailed description of how each feature is computed, refer to SI methods. While, in the main text, we present the effects of lesions on models with the combination of all mechanisms of homeostasis, we also report the effects of lesions on models with *G*^*E*^ and *G*^*E*^ + *c*^*EI*^ homeostasis (Figs S27-28).

### Code

All simulations and analyses were run using in-house scripts in Python. The code can be consulted in https://gitlab.com/francpsantos/wc network homeostasis.

## ACKNOWLEDGMENTS

FPS is supported by the European Commission through the euSNN project (Erasmus+ MSCA-ITN ETN H2020dID 860563). PFMJV is supported by AISN (Horizon Europe, 101057655), EBRAINS-HEALTH (Horizon Europe, 101058516), and PHRASE (European Innovation Council, 101058240) and the euSNN project (Erasmus+ MSCAITN ETN H2020 860563) and ReHyb (Horizon2020 871767).

## Supporting Information for

### Supporting Information Text

#### Methods

##### Computation of Homeostatic Parameters

In (1) we develop the analytical methodology to compute the value of different parameters of the Wilson-Cowan model allowing for the maintenance of stable activity under various mechanisms of E-I homeostasis. While we present the mathematical expressions here, refer to (1) for a detailed derivation and empirical basis for each mechanism of homeostasis. In all equations below, *r*^*E*^ represents the fixed point *r*^*E*^ corresponding to the target activity actively maintained by cortical networks, here represented by the Wilson-Cowan model.

###### Excitatory Synapses unto Pyramidal Populations

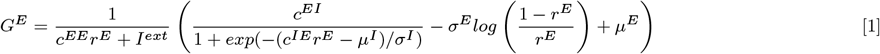

###### Inhibitory Synapses unto Pyramidal Populations

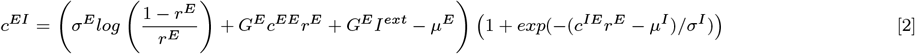

###### Intrinsic Excitability of Excitatory Populations

Homeostatic plasticity of intrinsic excitability, which modulates the parameters of the activation function *F* ^*E*^(*x*), can be implemented in two manners, considering empirical data (1). The first is at the level of the firing threshold of excitatory populations, *µ*^*E*^:

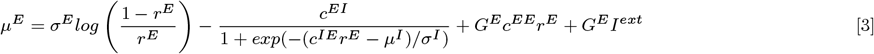

In the second case, E-I homeostasis modulates both the firing threshold *µ*^*E*^ and the sensitivity of the activation function σ^*E*^ in a coordinated manner so that the value of *F*^*E*^(0), or the baseline firing rate under no external input, is maintained. To do this, we consider that both parameters can be related to each other as σ^*E*^ = *Kµ*^*E*^, where *K* is a constant. For more details on the derivation, refer to (1). Here, we use the default parameters to compute *K*, so that 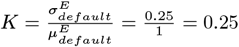. That said, the values of both parameters can be obtained through the following expression:

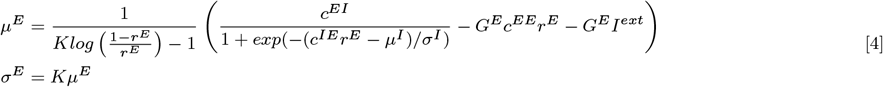

###### Synaptic Scaling of Excitatory and Inhibitory Synapses

Considering that plasticity of excitation and inhibition operate under the same timescale, it is possible to estimate the steady-state values of both parameters given the initial values 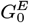 and 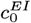. For a detailed derivation of the expressions and the extension of the framework to the case of different timescales, please consult (1).

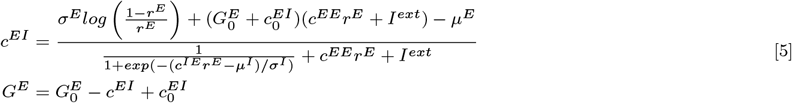

###### Synaptic Scaling of Excitation and Inhibition and Plasticity of Intrinsic Excitability

Following the previous implementations, we implement homeostasis of excitation and inhibition in conjunction with the two forms of homeostasis of intrinsic excitability. First, for plasticity of *µ*^*E*^, we have:

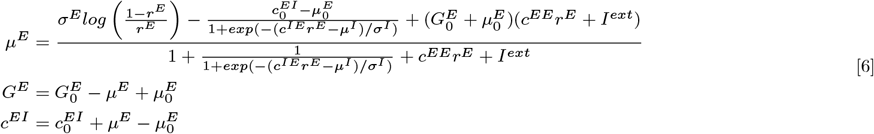

Similarly, for plasticity of *µ*^*E*^ and σ^*E*^, we obtain:

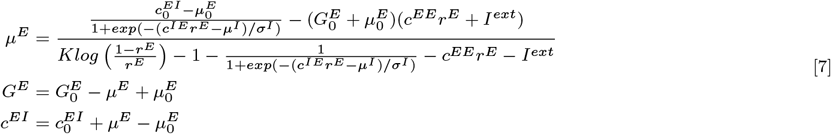

###### Computation of Homeostatic Parameters as a Function of the Target Firing Rate ρ

When the fixed point of the Wilson-Cowan model corresponds to a stable fixed point or spiral, the mean activity of the excitatory population is precisely equal to the fixed-point *r*^*E*^. However, in systems in a limit-cycle regime, the average firing rate differs from 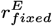. For this reason, in (1), we derive a method to obtain an estimate of the steady-state model parameters as a function of *ρ* and *I*^*ext*^, based on the following iterative algorithm, with *ϵ* = 10^−10^. For more details, refer to (1).

1. Define upper and lower limits for 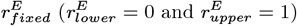
2. For 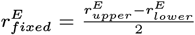, compute the homeostatic parameters of the system under *I*^*ext*^ using the equations in the previous sections with 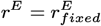.
3. Solve the system numerically for 5 seconds and compute the average firing rate ⟨*r*^*E*^⟩ from the last 2 seconds of activity.
4. If |⟨*r*^*E*^⟩ − *ρ*| < *ϵ*, take 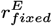 as the fixed point corresponding to *ρ* and save the homeostatic model parameters.
5. Else, if ⟨*r*^*E*^⟩ *> ρ*, restart the procedure from step 1, updating 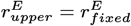. Conversely, if ⟨*r*^*E*^⟩ < *ρ*, restart from step 1 with 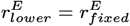.

##### Excitatory-Inhibitory Homeostasis in the Wong-Wang Model

###### Derivation of Homeostatic Expressions for the Wong-Wang Model

To derive the expressions allowing for the computation of parameters at the homeostatic fixed point we start with the equations for the fixed points of the model, when 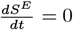 and 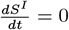:

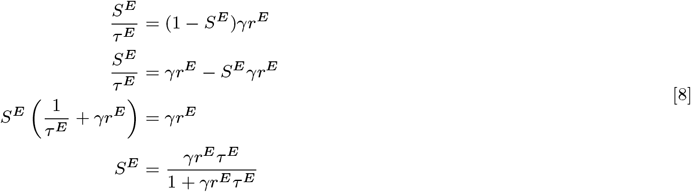

and

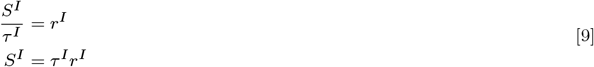

These equations can then be used together with the expressions for *I*^*E*^, *I*^*I*^, *r*^*E*^, and *r*^*I*^ for the mathematical estimation of the parameter values at the fixed point.

###### Synaptic Scaling of Excitation

Similarly to the Wilson-Cowan model, the excitatory synapses unto the excitatory population can be modulated through the parameter *G*^*E*^. To compute the value of *G*^*E*^ at the fixed point, as a function of the external input (*I*^*ext*^) and the target firing rate 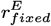 we start by computing the value of 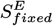 can be computed from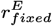 using expression 8.

Then, to compute the value of 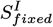 we first substitute *S*^*I*^ by *τ* ^*I*^ *r*^*I*^ (Equation 9), obtaining a function that can be solved for *I*^*I*^ to obtain its fixed point value:

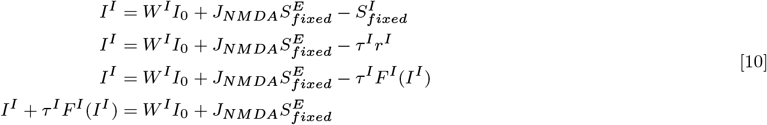

The resulting value of 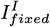 can then be substituted in equation 9 to obtain 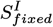.

Then, 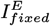 can be calculated by solving the following expression for *I*^*E*^:

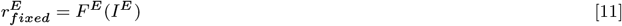

Finally, we re-organize the equation for *I*^*E*^, obtaining the following expression:

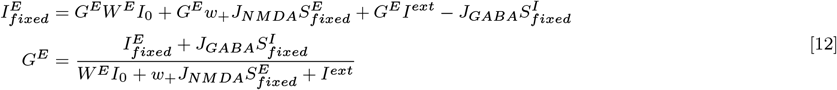

where the values of 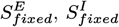, and 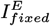 can be inserted to calculate the value of *G*^*E*^.

###### Synaptic Scaling of Inhibition

Similarly to the synaptic scaling of excitation, the value of *J*_*GABA*_ corresponding to the homeostatic fixed point can be estimated from the expression for *I*^*E*^ as follows:

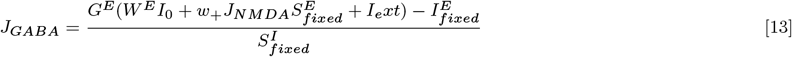

###### Plasticity of Intrinsic Excitability of Excitatory Populations

In the Wong-Wang model, the equivalent of the firing threshold of the input-output function is the parameter *b*^*E*^. The effect of changing *b*^*E*^ on *F* ^*E*^(*x*) can be visualized in Figure S27.

In this case, we start by obtaining 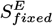 from expression 8 and then solve the following equation for *S*^*I*^ to obtain 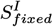:

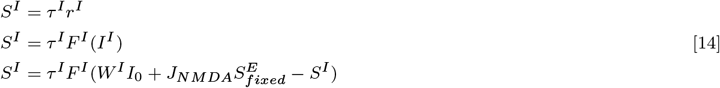

which can then be used, together with 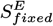 to compute 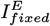.

Finally, the homeostatic value of *b*^*E*^ at the fixed point can be obtained by solving the following expression for *b*^*E*^:

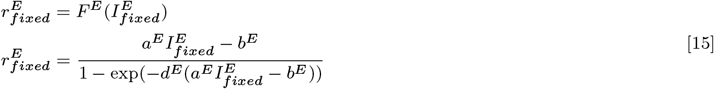

Similarly, the plasticity of excitability slopes can be approximated by modulation of the parameter *a*^*E*^. In this case, since *F* (*x*) is not a sigmoid function (Figure S27) and approaches *a*^*E*^*x* −*b*^*E*^ as *x* increases, the effect described in (2–4), whereby the slope is modulated without changing the output of the function when the input is equal to 0, can be approximated by varying *a*^*E*^ only. Not only that, since 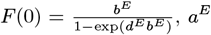 can be changed without affect the value of *F* (*x*) at 0. We present a visual demonstration of this effect in Figure S27.

That said, the homeostatic value of *a*^*E*^ can be computed by solving expression 15 for *a*^*E*^ instead of *b*^*E*^.

###### Synaptic Scaling of Excitatory and Inhibitory Synapses

Similarly to the procedure applied in (1) for the Wilson-Cowan model, we consider that the homeostasis of excitation and inhibition operate under the same timescale. Therefore, in line with (1), we consider that:

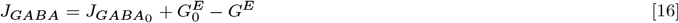

where 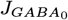 and 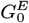 are the default values presented in Table S3.

Then, we substitute *J*_*GABA*_ with equation 16 in 12, obtaining:

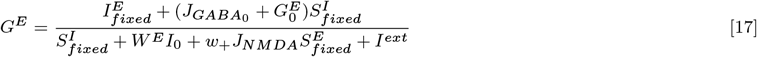

###### Synaptic Scaling of Excitatory and Inhibitory Synapses and Plasticity of Intrinsic Excitability

In line with the methodology implemented for the Wilson-Cowan model, we consider the homeostasis of excitation and inhibition in conjunction with the two forms of plasticity of intrinsic excitability. In this case, since the order of magnitude of *J*_*GABA*_ and *G*^*E*^ is much lower than *b*^*E*^ or *a*^*E*^ (see Table S3), to consider similar time scales for all homeostatic mechanisms we instead assume that their **relative** variation is the same, as opposed to absolute variation as done in (1). Therefore, when implementing the homeostasis of *b*^*E*^ in conjunction with the two forms of synaptic scaling we have:

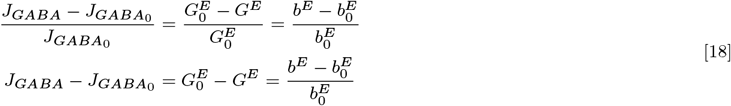

since 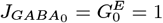, which simplifies the expression.

That said, *G*^*E*^ and *J*_*GABA*_ can be substituted in the expression for *I*^*E*^, yielding:

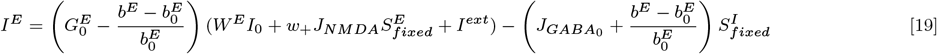

The value of *b*^*E*^ can then be obtained by solving equation 15 for *b*^*E*^.

Similarly, for *a*^*E*^, we have

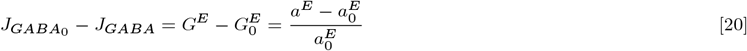

which can be substituted in the *I*^*E*^ equation, yielding:

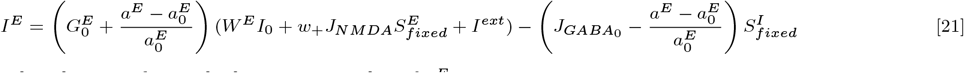

which can be used with 15 to obtain the homeostatic value of *a*^*E*^.

###### Modulation of Local Circuit Dynamics by E-I Homeostasis in the Wong-Wang Model

To analyze the effects of E-I homeostasis at the local circuit level in the Wong-Wang Model we explore the behavior of the model under different combinations of target firing rate 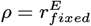 and external input *I*^*ext*^ and distinct mechanisms of E-I homeostasis. For each combination of *ρ* and *I*^*ext*^, we compute the homeostatic values of the parameters of interest, following the expressions in the previous section, and follow up with linear stability analysis to investigate circuit dynamics (5, 6). To do that, it is first necessary to compute the Jacobian *J* of the system:

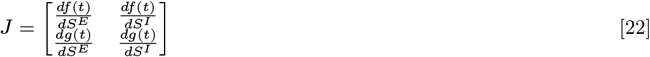

where 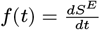 and 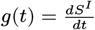.

The entries of the Jacobian can be calculated as follows:

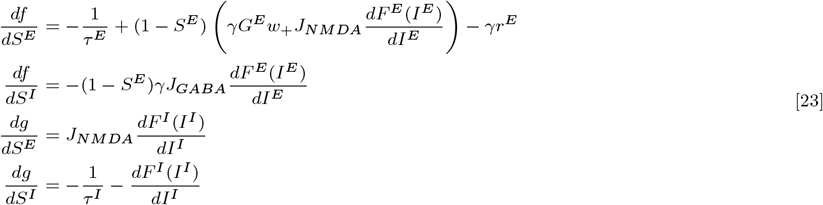

where

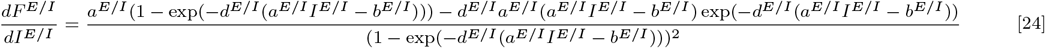

Then, the local dynamical regimes can be evaluated by analyzing the trace and determinant of the Jacobian (5, 6). We present the results of this analysis for all homeostatic mechanisms in Figure S28. In the case of the Wong-Wang model, it can be observed that E-I homeostasis does not have the same effect on local dynamics as for the Wilson-Cowan model (e.g. Figure S1). More specifically, there is no Hopf-bifurcation in the Wong-Wang model and, thus, regardless of the effects of homeostasis, the local circuits remain in a stable regime. Even though the plasticity of *J*^*GABA*^ can lead to the emergence of a stable spiral attractor (Figure S28B), with transient rhythms in response to perturbations, the dynamics still correspond to a stable attractor and, therefore, this transition is not considered a bifurcation.

#### Analysis of FC Properties in Lesion Simulations

To analyze the impact of structural lesions on FC and the subsequent recovery through E-I homeostasis, we follow the procedure introduced in (7). To analyze changes in FC patterns, we make use of FC distance, which roughly quantifies the magnitude of disruption, following (7, 8). Furthermore, to measure disruptions in the macroscale architecture of FC, we measure the correlation between structural and functional connectivity (9) and modularity (10, 11), both of which are impacted in stroke patients. Furthermore, the recovery of modularity is thought to be relevant for cognitive function (11). Below, we describe how each of these metrics is computed from simulated FC matrices. The simulation protocol is described in the Methods section of the main text.

#### FC Distance

To measure the dissimilarity between FC matrices at baseline, acute, and chronic periods, we follow (8), defining FC distance as the Frobenius norm of the difference between two given matrices.

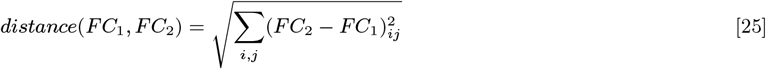

#### FC-SC Correlation

Given the results of (9), showing a decoupling between functional and structural connectivity in stroke patients, correlating with motor function, we test this biomarker at baseline, acute, and chronic periods, by computing the Pearson’s correlation coefficient between the upper triangles of FC and SC matrices.

#### Modularity

Modularity measures the degree to which a network follows a modular structure, with dense connections within functional clusters and sparser ones between them. Modularity (*Q*) was calculated using the formula defined in (12):

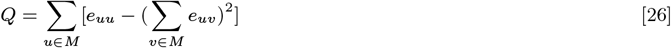

where *M* is a set of non-overlapping modules (groups of nodes) in the network and *e*_*uv*_ is the proportion of edges in the network that connect nodes in module *u* with nodes in module *v*. Similarly to previous studies, (7, 11), we define modules *a priori* to avoid biasing the modularity measure by using modularity maximization to detect community structure. That said, modules were from the empirical FC data, by using a clustering algorithm described in (7), resulting in 6 clusters. Following the original formulation of modularity (12), FC matrices were transformed into unweighted graphs by applying a density threshold, through which only a percentage of the strongest connections are kept and their weights set to 1. Lesioned nodes were removed from the network before computing modularity, similarly to (7, 11).

#### Model Validity for Different Mechanisms of Homeostasis

For each type of homeostasis, we run short simulations (15 seconds) with all combinations of parameters and evaluate the deviation of the mean node activity from the target firing rate *ρ*. If any node deviates by more than 1%, we consider the fixed point invalid.

To start, we illustrate the impact of each parameter on model validity for models under *G*^*E*^ homeostasis and discuss the principles underlying the ability of networks to maintain target firing rates in different regions of the parameter space represented by the combination of the free parameters (*C, rho* and mean delay). We start by using models with *G*^*E*^ homeostasis to illustrate the different behaviors of our model (Figure S2). However, the general principles apply to other types of homeostasis, as will be illustrated later. The respective parameter spaces can be consulted in Figure S7.

The first main conclusion is that, as the target firing rate *ρ* increases, the models become increasingly unable to maintain firing rates at the target and, for any value of *ρ* higher than 0.15, no combination of parameters resulted in a valid system. This result is easily interpreted. As more nodes enter the limit cycle regime (S1) the presence of sustained oscillations leads to a tendency of nodes to synchronize in a self-reinforcing manner, generating run-away activity and resulting in the instability of the fixed point solution. Therefore, to guarantee the stability of models, the target firing rate *ρ* should be close to the threshold after which some nodes enter the limit cycle regime, corresponding to *ρ* = 0.12.

To delve into this topic in more detail, we examine the behavior of the model when *ρ* = 0.14. While most of the parameter space corresponds to an invalid solution of the system, systems can maintain mean firing rates if the global coupling is high enough (Figure S2). To study this effect, we simulated models with the same *ρ* (0.12) and mean delay (10 ms), but with low (*C* = 0.75) and high (*C* = 7.5) global coupling (Figures S3 and S4, respectively). For each, we ran simulations with the predicted homeostatic *G*^*E*^, no noise, and different initial conditions for node activity *r*^*E*^ (0, *ρ* or 2*ρ*). In addition, we ran simulations with dynamical *G*^*E*^ homeostasis (obeying 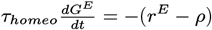) and different homeostatic timescales, to ensure that the instabilities were inherent to the model and not a consequence of errors in the estimation of homeostatic parameters. Starting with the lower global coupling (*C* = 0.75) (Figure S3), we observe that, regardless of the initial conditions, the model with predicted *G*^*E*^ tends toward global synchronization and, since long-range connections are excitatory, the oscillations self-reinforce and quickly reach activity levels close to saturation. How fast this happens depends on the initial conditions. Can this be fixed by implementing explicit homeostatic plasticity of *G*^*E*^, instead of using the predicted weights? Our results indicate that such models, having the same tendency to synchronize, will exhibit short periods of high activity departing from *ρ*, which are then quenched by homeostatic plasticity. In this case, the speed of this quenching depends on the timescale of homeostatic plasticity (Figure S3). For this reason, we consider these models to still be invalid, indicating that the inability to maintain mean firing rates is an inherent property of systems with this combination of parameters and not due to errors in the estimation of parameters. Indeed, when observing the behavior of individual nodes in the homeostatic fixed points (Figure S3), we see that 52 of them are in the limit cycle regime, which explains the tendency of the model for over-synchronizing.

But why are models with *ρ* = 0.14 when the global coupling is increased? In Figure S4, we show one such case where, regardless of the initial conditions, all nodes in the model converge toward the target firing rate. Following the principles introduced before, we show that the homeostatic solution to this system corresponds to only one node in the limit cycle regime. This phenomenon occurs due to how model dynamics are shaped by *G*^*E*^ homeostasis (Figure S4). Shortly, for a given value of *ρ*, as the external current is increased, the models transition from a limit cycle to a stable spiral regime. Therefore, given that *I*^*ext*^ scales with the global coupling, increasing the global coupling will bring nodes out of the limit cycle regime, contradicting the tendency to hypersynchrony. This effect is only present for some types of homeostasis (Figure S1) which has implications for the validity of models. Conversely, in models with *µ*^*E*^ or *c*^*EI*^ homeostasis, when *ρ* ≥ 0.12 all nodes are necessarily in the limit cycle regime and, therefore, have a greater tendency toward instability.

That said, it should be pointed out that, for target firing rates lower than 0.12, the general effect of increasing the global coupling is the opposite, disrupting the ability of models to maintain mean firing rates at the target. However, in this case, the underlying phenomena are distinct. In Figure S5 we present the behavior of the model in one such case, where the model settles in different fixed points depending on the initial conditions. If the initial *r*^*E*^ is higher than the target firing rate, the model will settle in a stable configuration where mean activity is higher than *ρ*. Conversely, the opposite happens when the model is initialized with *r*^*E*^ = 0. More importantly, the target fixed point of the system is only stable when the initial conditions correspond to *r*^*E*^ = *ρ* across all nodes. These results indicate that, for this combination of parameters, the model is bistable, with the target corresponding to the saddle point of the system, thus being unstable. Accordingly, when implementing explicit homeostatic plasticity (Figure S5), the system will constantly switch between the two stable attractors. Since the target firing rate is different from *ρ* in both, homeostatic plasticity will be permanently frustrated, leading to the constant switching between attractors. For this reason, even though the model can be stable in this case, the stable solutions do not correspond to the target firing rate and, therefore, we consider this to represent an invalid solution of the system. While we performed this analysis in the model with *G*^*E*^ homeostasis, the same principles apply, for example, to models with *G*^*E*^ + *c*^*EI*^ homeostasis (Figure S10).

Our results so far suggest that the main parameters shaping the ability of models to maintain the target firing rate are the global coupling and *ρ*. However, there are instances where the mean delay also plays a significant role. As an example, we take *C* = 4.5 and *ρ* = 0.11 (close to the bifurcation after which nodes with weaker inputs start entering the limit cycle regime) and examine model activity for 4 different values of mean delay (Figure S6). Starting with the model with no delays (mean delay=0), we observe that the system tends toward hyper-synchrony, similar to what was described for high values of *ρ*. As the mean delay is increased to higher values, this tendency disappears and the model can settle in the target fixed point. Here, our results demonstrate the role of delayed interactions in avoiding widespread synchronization, through the disruption of phase-relations between oscillating systems. However, we observe a range of mean delays centered around 26 milliseconds, for which the model can also be invalid (Figure S6). In this case, the target fixed point is only weakly stable, since, when approached from one direction, dynamics settle at *ρ*. However, if the initial conditions are higher than the target, some of the nodes engage in sustained oscillations potentiated by their recurrent interactions. While this case is not as severe as the model with no delays, it still represents a solution of the system that does not correspond to the target *ρ*. But why does this happen specifically for this range of delays? For this combination of *C* and *ρ*, most of the nodes have a natural frequency of oscillation between 45 and 60 Hz (Figure S6). More importantly, the nodes with the lower input, which are the ones closer to the limit cycle regime (S1), oscillate at around 45 Hz, corresponding to a period of ∼22.5 ms. Therefore, when the mean delay is close to this value, nodes in the system can still synchronize, albeit with a phase shift close to a full cycle, which is sufficient to throw the system outside of the desired equilibrium. Accordingly, as the mean delay is further increased, the models become valid again (Figure S6). While we performed this analysis in the model with *G*^*E*^ homeostasis, the same principles apply, for example, to models with *c*^*EI*^ (Figure S8) or *G*^*E*^ + *c*^*EI*^ (Figure S9) homeostasis.

Finally, for each mechanism of homeostasis, we plot the percentage of valid simulations as a function of each free parameter (*C, ρ*, and mean delay) and the percentage across all simulations (Figure S11). Our results suggest that, across the various mechanisms of homeostasis, the main parameters shaping model validity are the global coupling *C* and target firing rate *ρ*. However, validity is affected in models with homeostasis of inhibition (*c*^*EI*^) (19.26% valid simulations), firing threshold (*µ*^*E*^) (14.80%), or the combined homeostasis of *G*^*E*^, *c*^*EI*^ and *µ*^*E*^ (20.47%), all of which strongly rely on the modulation of “additive/subtractive” model parameters (1). Conversely, homeostasis mechanisms involving excitatory synapses (*G*^*E*^) or the slope of neural excitability (σ^*E*^) (i.e. “multiplicative parameters”) enhance model validity (Figure S11D). We suggest that this results from the modulation of the bifurcation point toward higher *ρ* in response to increases in external inputs, which is characteristic of these mechanisms of homeostasis (1) (Fig S15). More specifically, when *C* is increased, amplifying not only the magnitude of external inputs, but also their fluctuations (Fig S16), nodes can compensate for the stronger input fluctuations by being farther away from the bifurcation. Conversely, with the homeostasis of *c*^*EI*^ or *µ*^*E*^, nodes with higher inputs are more likely to engage in sustained oscillations (Fig S15), enhancing network instabilities. In short, the bifurcation modulation effect afforded by the homeostasis of excitation and the slope of intrinsic excitability is an important feature of local cortical circuits, ensuring that the target firing rate can be maintained across the network.

**Fig. S1.**
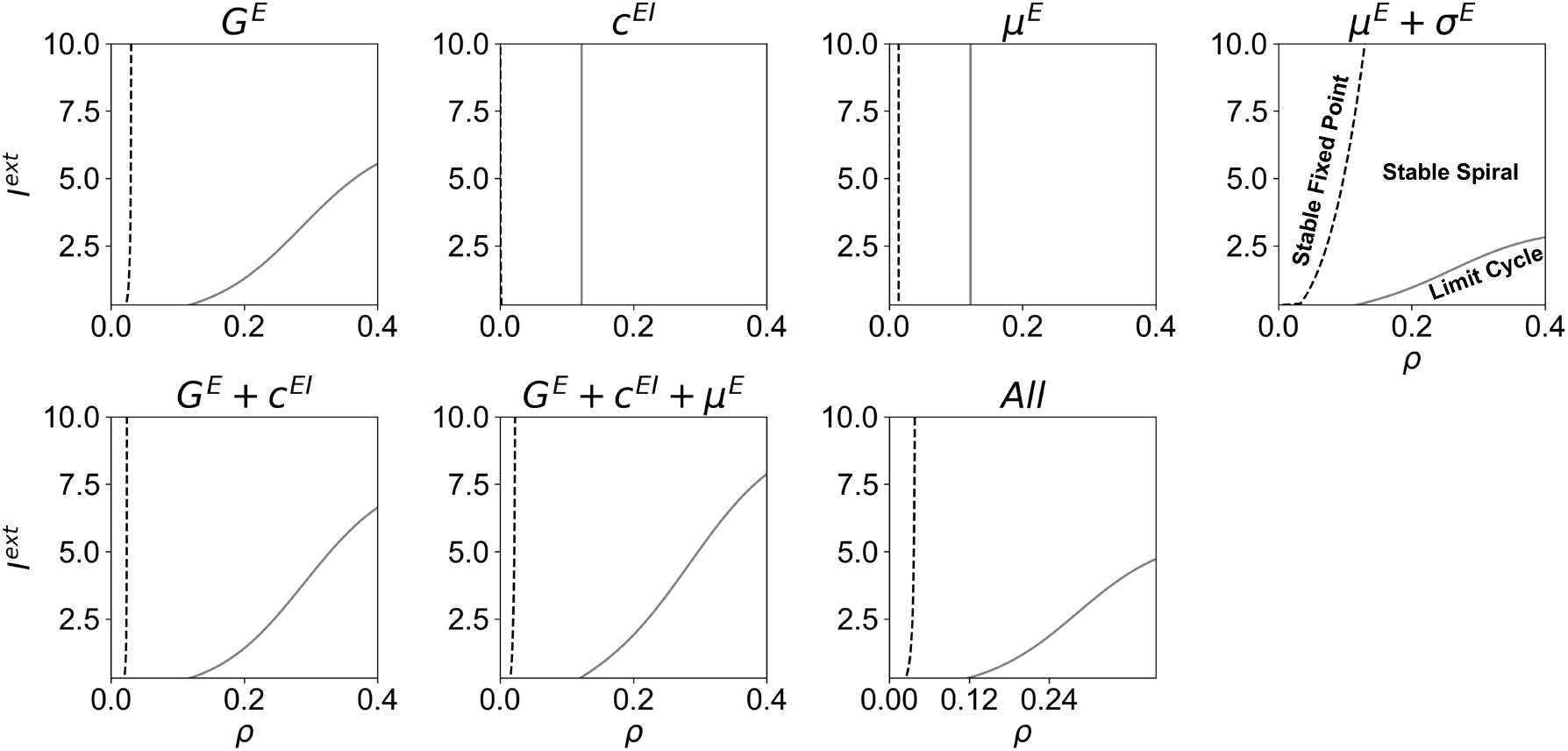
Phase portraits of the Wilson-Cowan model under different mechanisms of homeostasis. Dashed lines represent the transition between the stable fixed point and stable spiral regimes and solid lines represent the Andronov-Hopf bifurcation between damped and sustained oscillations (i.e. limit cycle). Refer to (1) for more detail on the formulation of each mechanism of homeostasis and the analysis of the dynamics of Wilson-Cowan nodes under the distinct mechanisms of homeostasis.

**Fig. S2.**
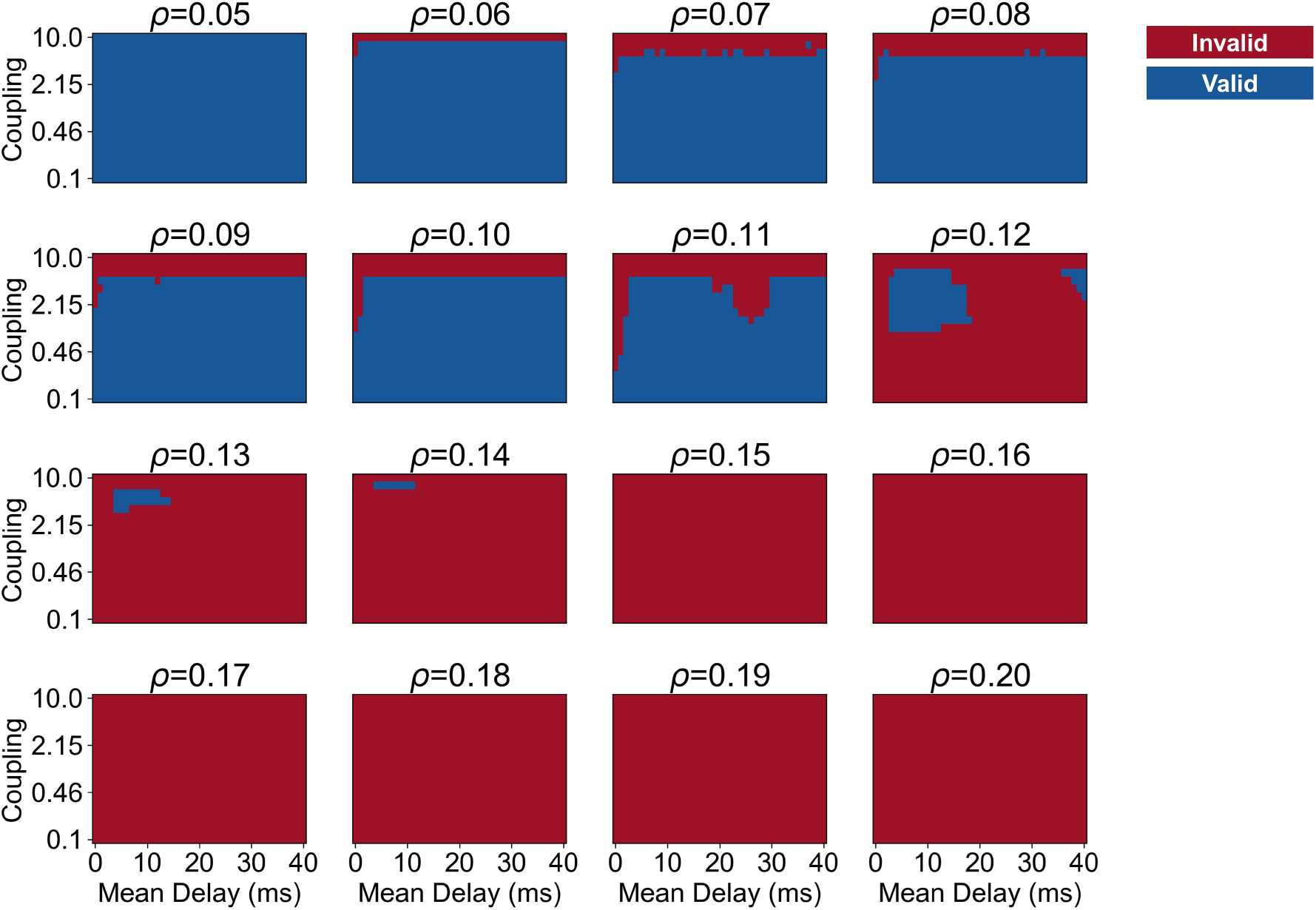
Model validity under different combinations of parameters for *G*^*E*^ Homeostasis. Red represents invalid models, where the mean activity of at least one of the nodes differed from the target firing rate *ρ* by more than 1%. Conversely, blue represents models where all nodes have mean activity close to the target.

**Fig. S3.**
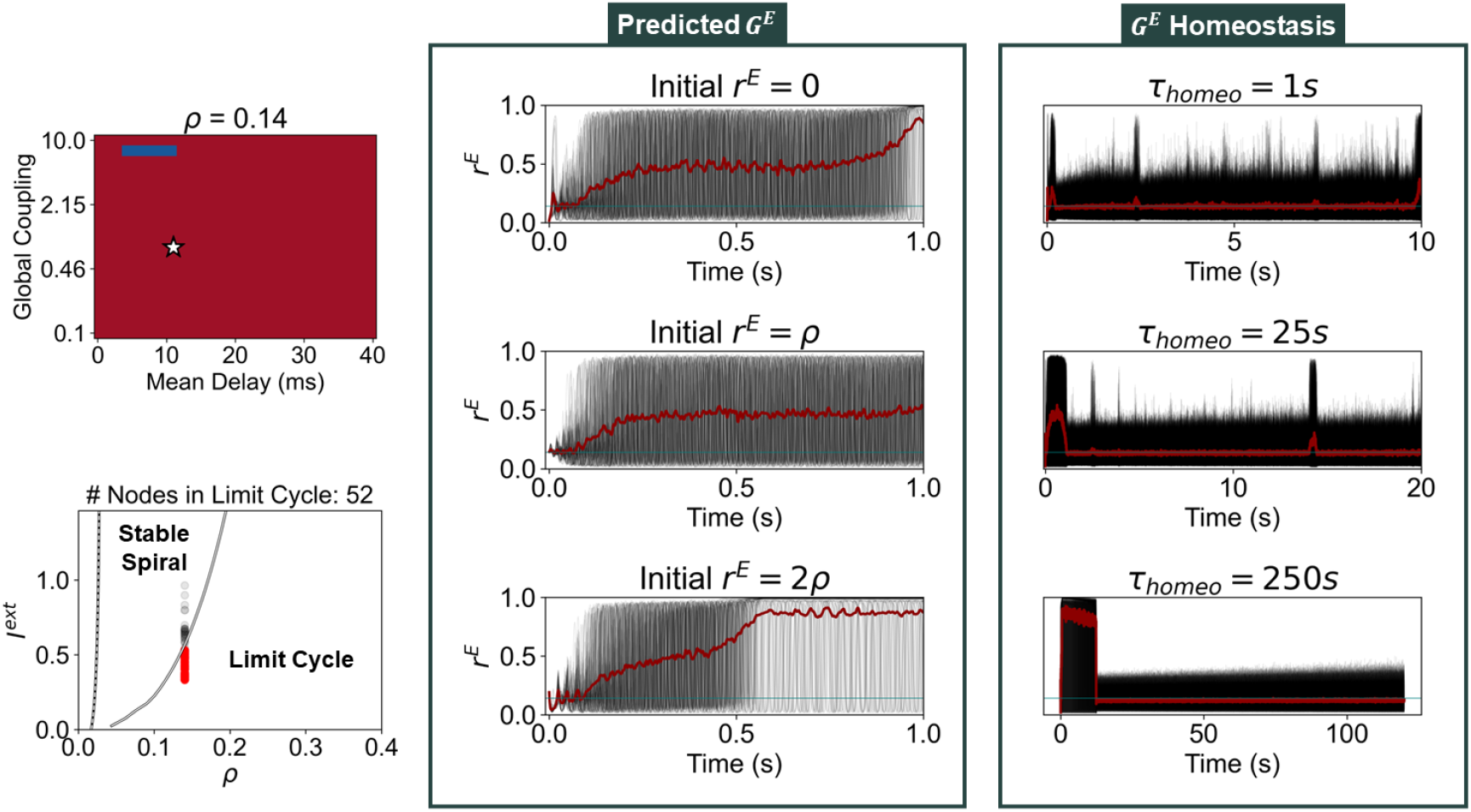
Network Behavior for *C* = 0.75, *ρ* = 0.14 and mean delay 10 ms. On the top left, we present the stability parameter space for *ρ* = 0.14, where the star represents the current parameters. On the bottom left, we present the dynamical portrait for *G*^*E*^ homeostasis at node level, with red and black dots representing network nodes in the limit cycle and stable spiral regimes, respectively. On the right, we present the activity of networks with predicted *G*^*E*^ values, for different initial conditions, and of networks with dynamical *G*^*E*^ homeostasis with different time constants. Black lines represent *r*^*E*^ of each node, red lines the average across nodes and blue lines the target firing rate *ρ*.

**Fig. S4.**
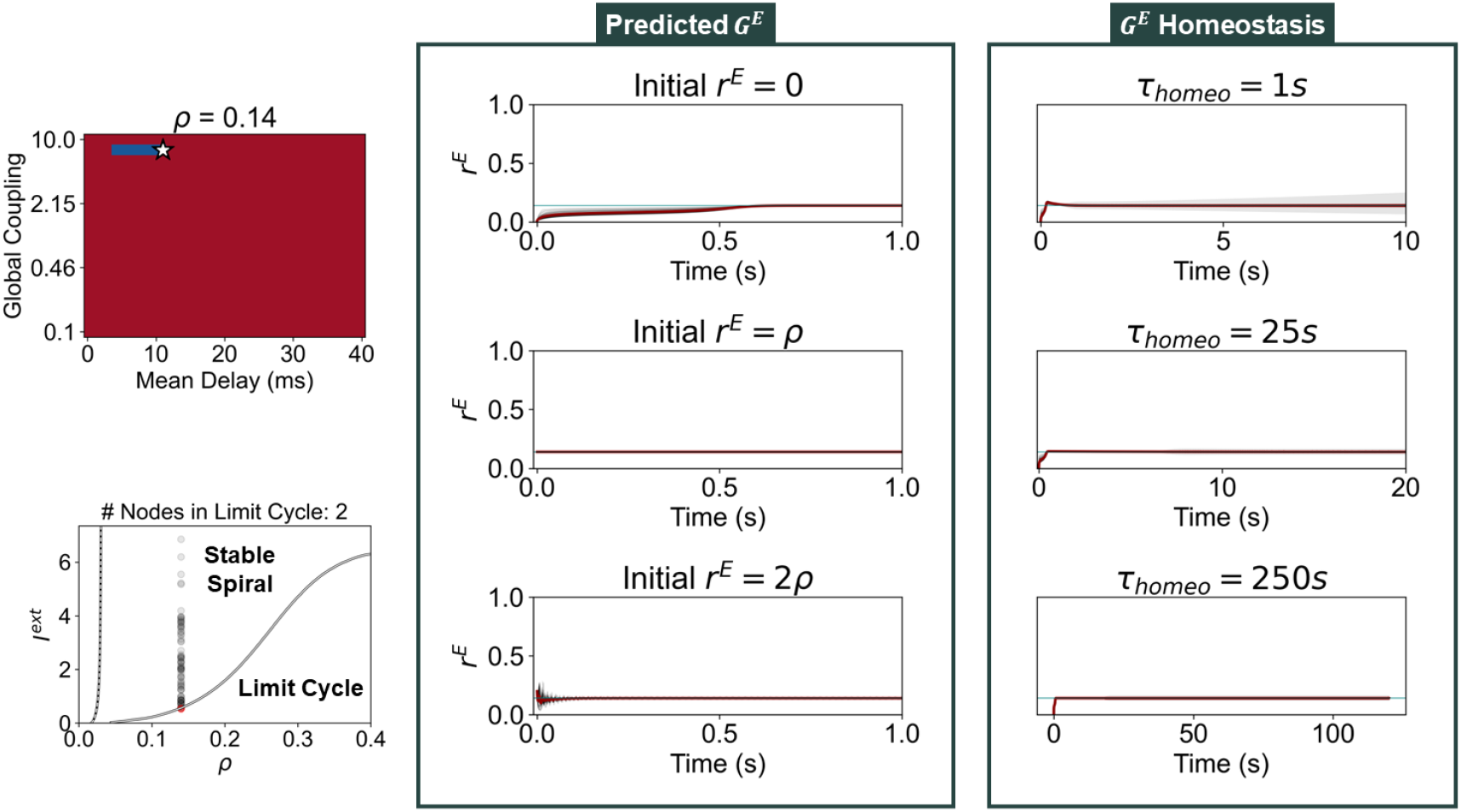
Network Behavior for *C* = 7.5, *ρ* = 0.14 and mean delay 10 ms.

**Fig. S5.**
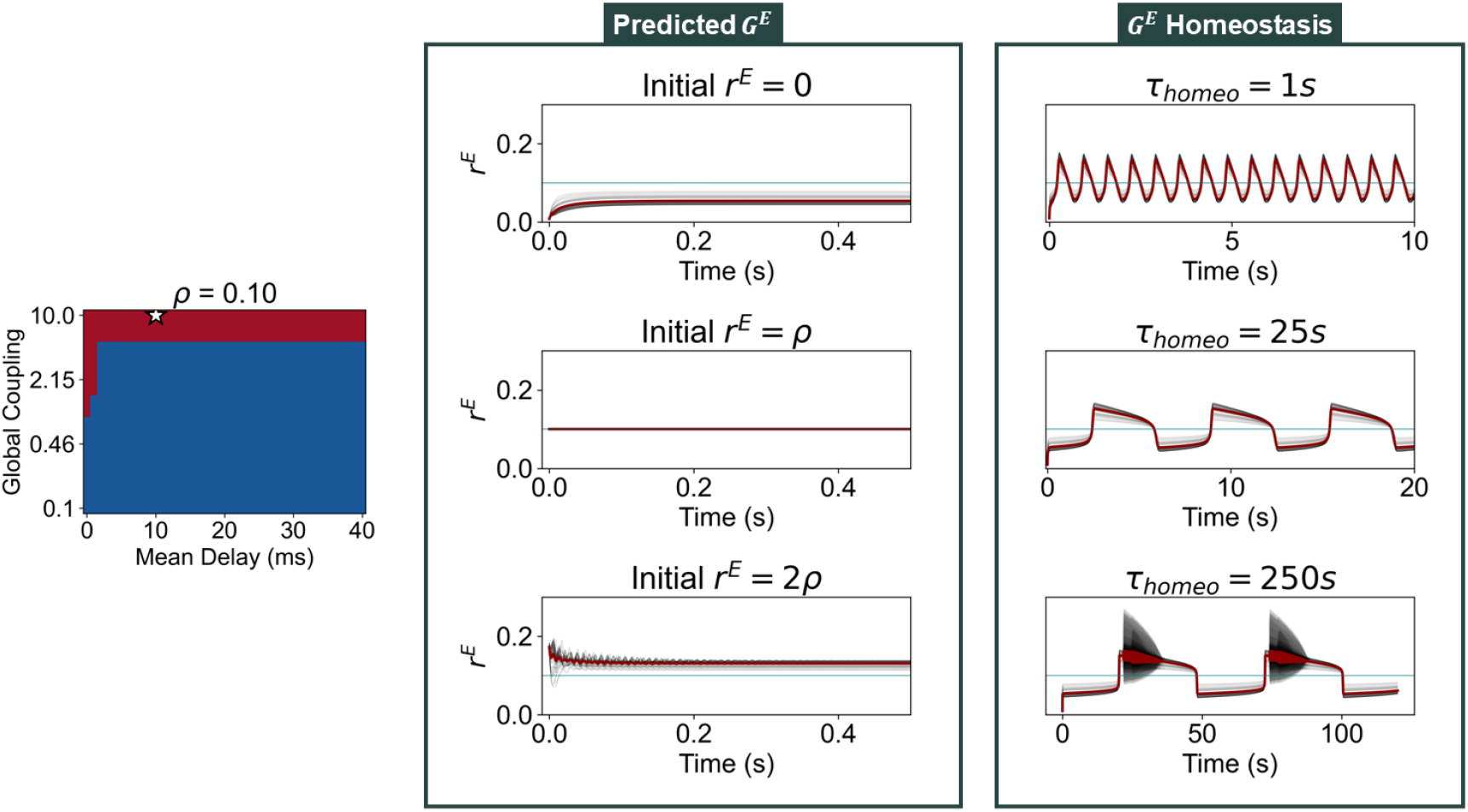
Network Behavior for *C* = 10, *ρ* = 0.10 and Mean Delay 10 ms.

**Fig. S6.**
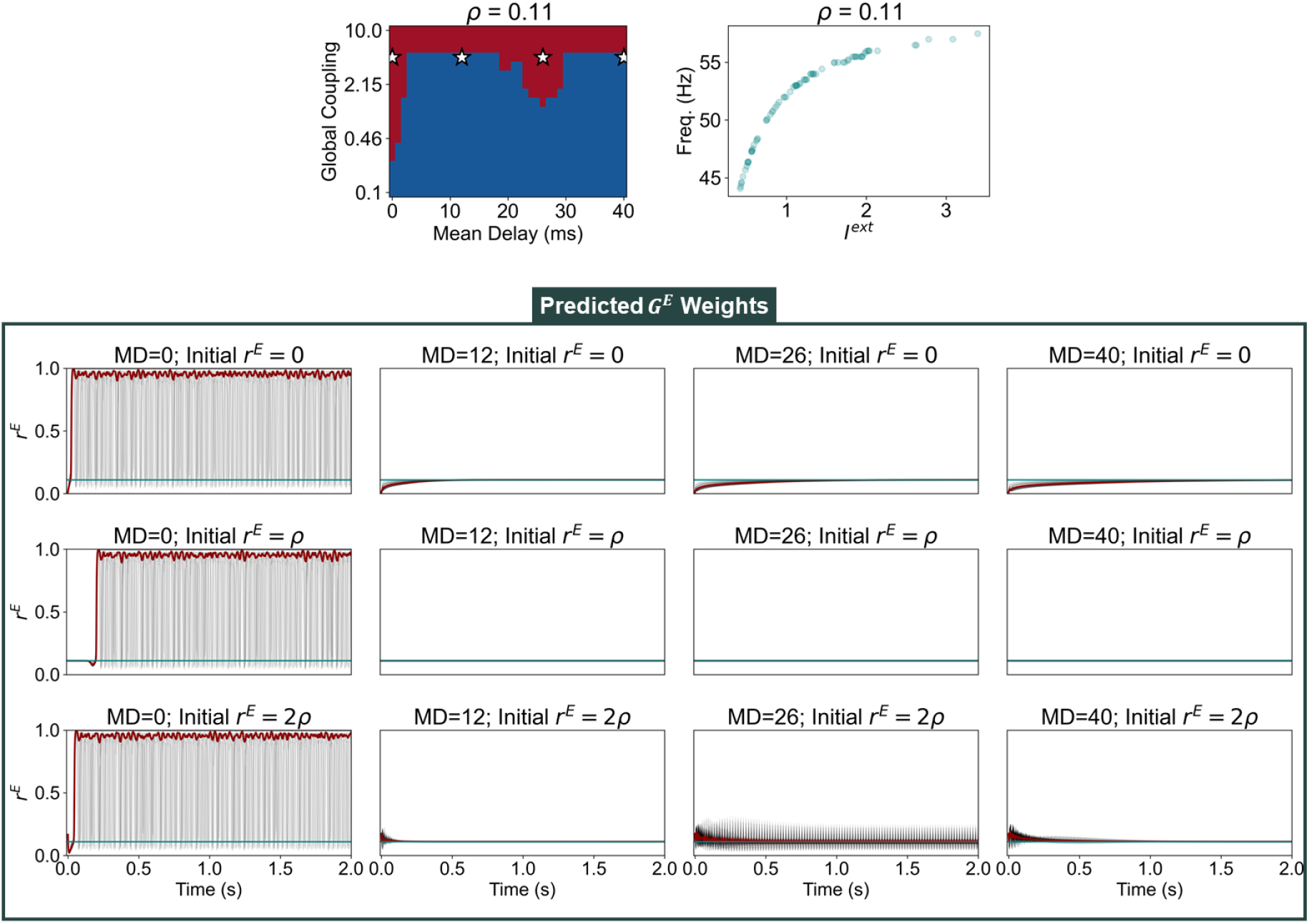
Network behavior for *C* = 4.5, *ρ* = 0.11 and different mean delays. On the top left, we present the validity parameter space of the model for *ρ* = 0.11. On the top right, we present the natural frequency of oscillation of nodes in the network as a function of the average external input they receive. On the bottom, we display network activity for models with mean delay 0, 12, 26 and 40 ms, and with different initial conditions. Black lines represent *r*^*E*^ of each node, red lines the average across nodes and blue lines the target firing rate *ρ*.

**Fig. S7.**
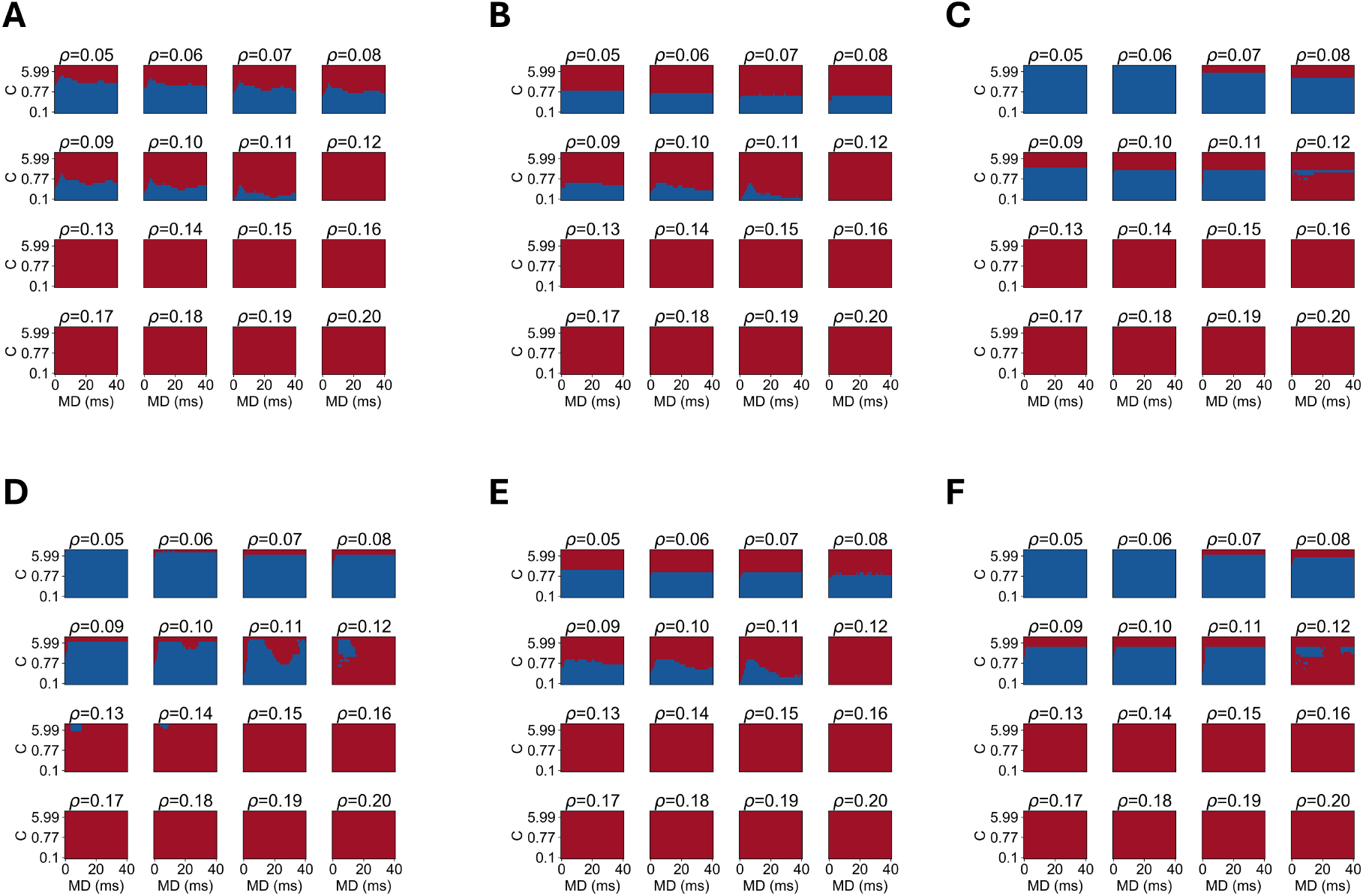
Validity of models with different mechanisms of homeostasis. (A) *c*^*EI*^ Homeostasis (B) *µ*^*E*^ Homeostasis (C) *µ*^*E*^ + σ^*E*^ Homeostasis (D) *G*^*E*^ + *c*^*EI*^ Homeostasis (E) *G*^*E*^ + *c*^*EI*^ + *µ*^*E*^ Homeostasis (F) *G*^*E*^ + *c*^*EI*^ + *µ*^*E*^ + σ^*E*^ Homeostasis. Blue and red colors represent valid and invalid simulations, respectively.

**Fig. S8.**
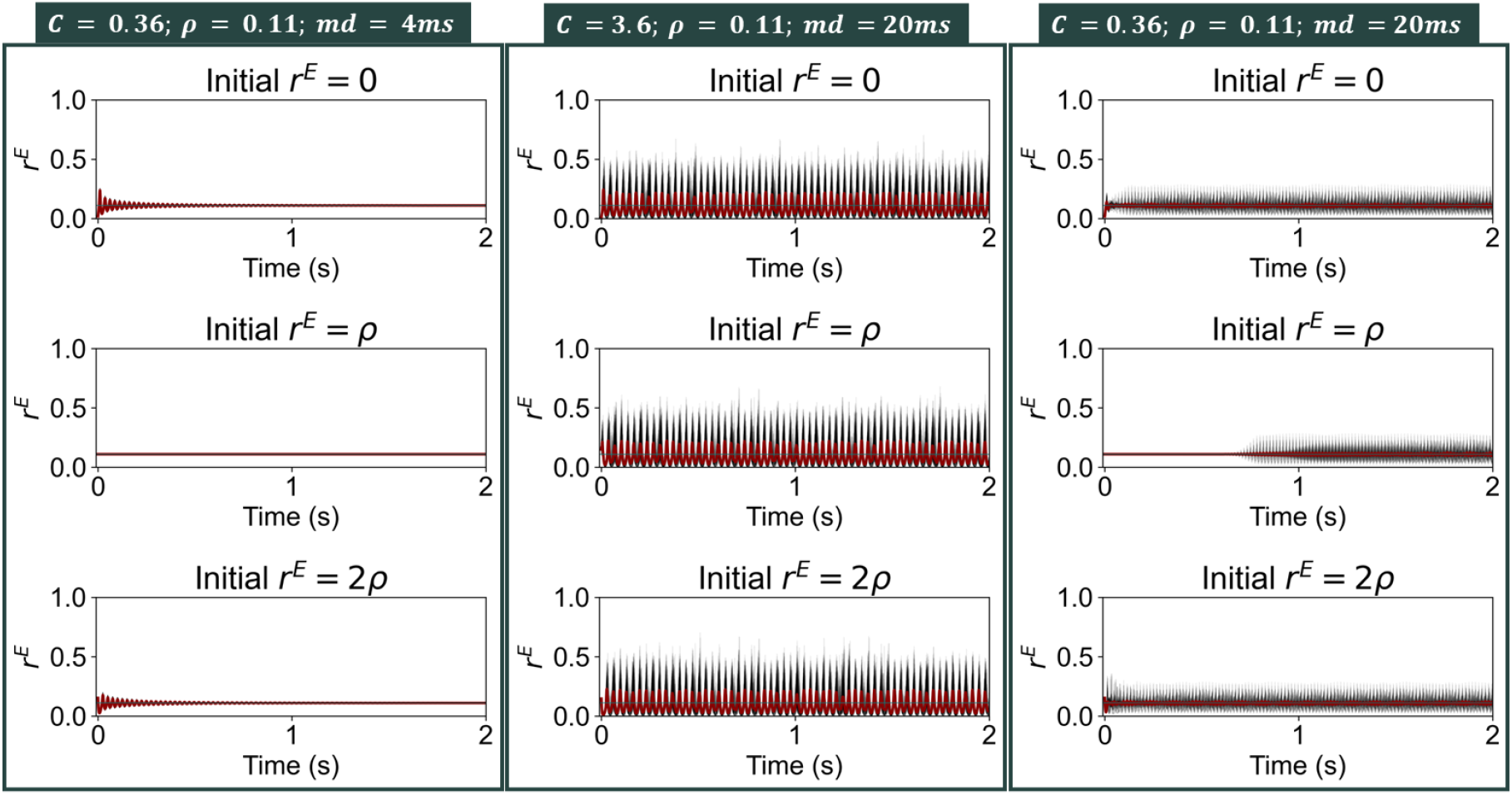
Behavior of the network with homeostasis of *c*^*EI*^. We present the activity of models with predicted *c*^*EI*^ values and different initial conditions: *r*^*E*^ = 0 (Top), *r*^*E*^ = *ρ* (Middle) and *r*^*E*^ = 2*ρ* (Bottom)

**Fig. S9.**
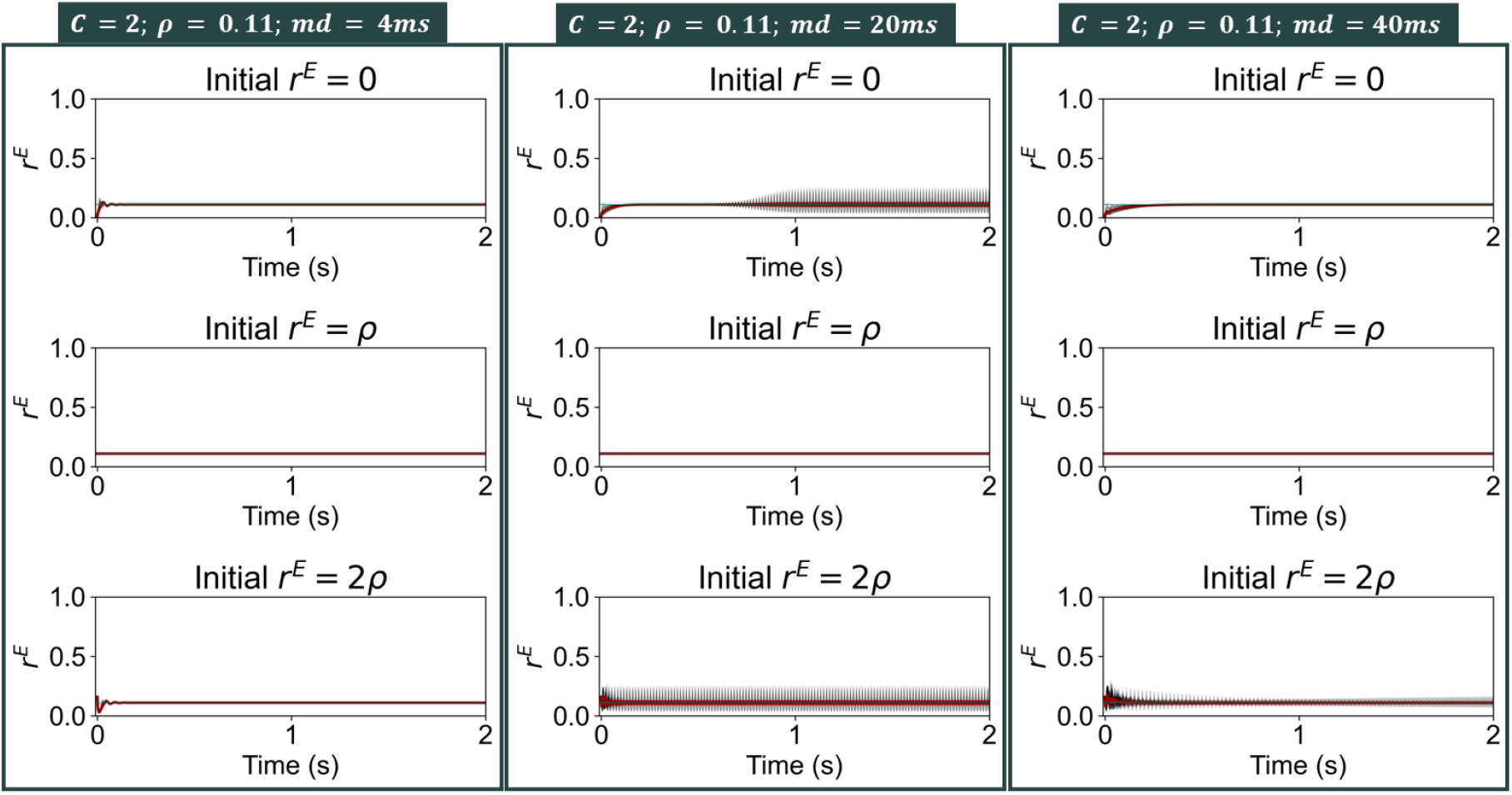
Behavior of the network with homeostasis of *G*^*E*^ and *c*^*EI*^. We present the activity of models with predicted parameters and different initial conditions: *r*^*E*^ = 0 (Top), *r*^*E*^ = *ρ* (Middle) and *r*^*E*^ = 2*ρ* (Bottom)

**Fig. S10.**
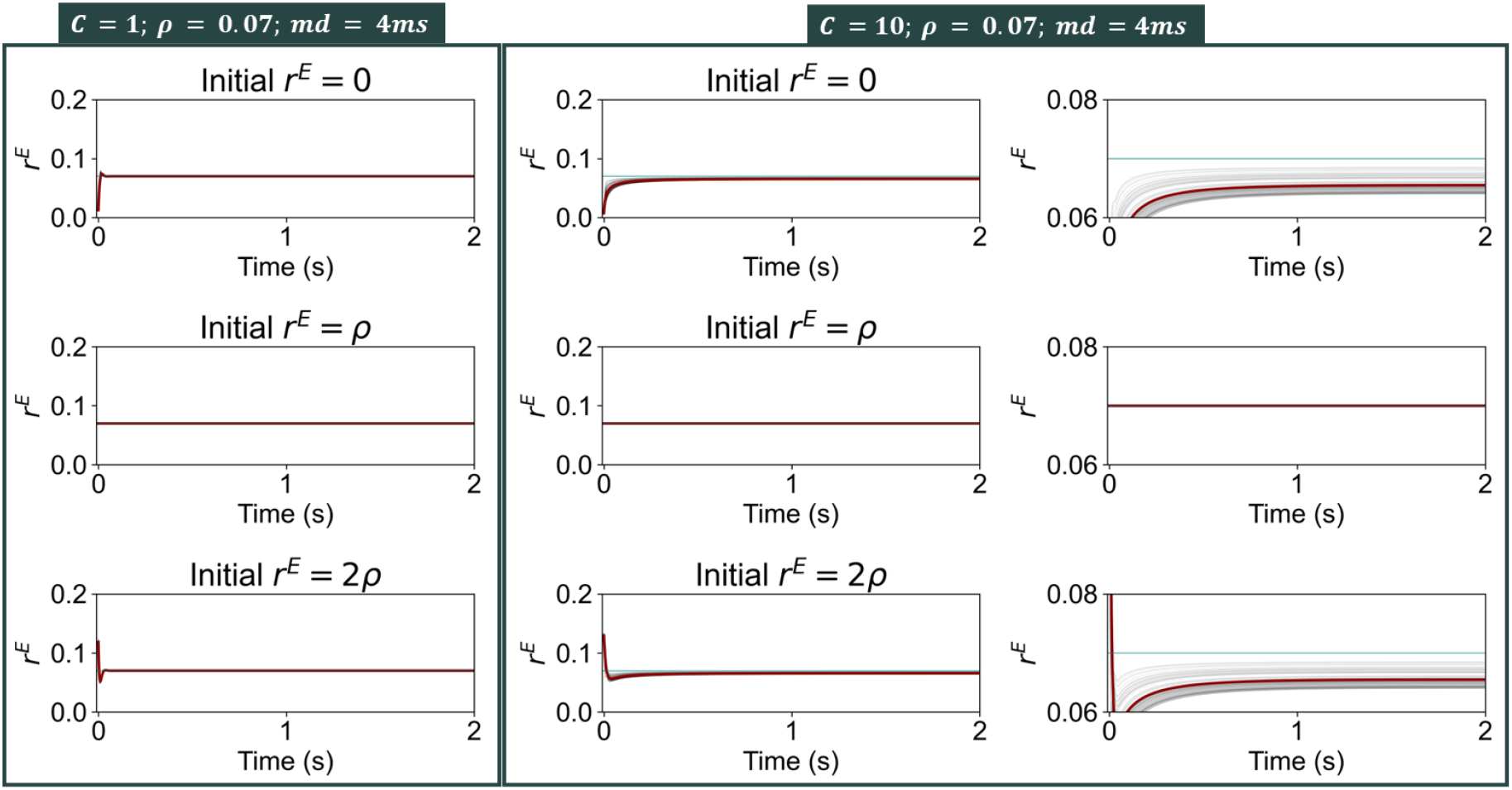
Behavior of the network with homeostasis of *G*^*E*^ and *c*^*EI*^. We present the activity of models with predicted parameters and different initial conditions: *r*^*E*^ = 0 (Top), *r*^*E*^ = *ρ* (Middle) and *r*^*E*^ = 2*ρ* (Bottom)

**Fig. S11.**
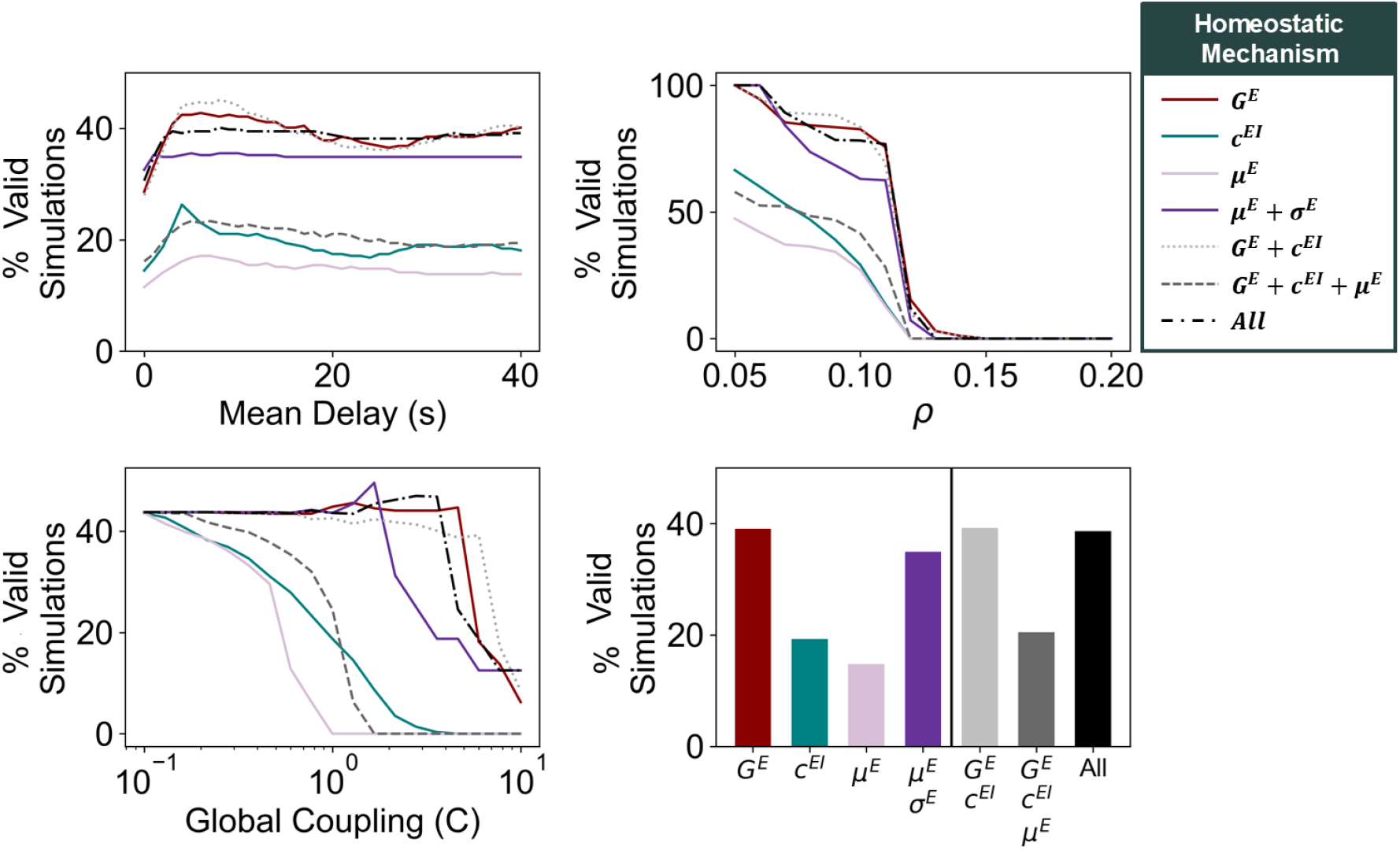
Model validity for different mechanisms of homeostasis. (A) Percentage of valid simulations as a function of the mean delay. Each line represents a different mechanism of E-I homeostasis. (B) Percentage of valid simulations as a function of *ρ* (C) Percentage of valid simulations as a function of *C*. (D) Percentage of valid simulations across all combinations of parameters for each mechanism of homeostasis

**Fig. S12.**
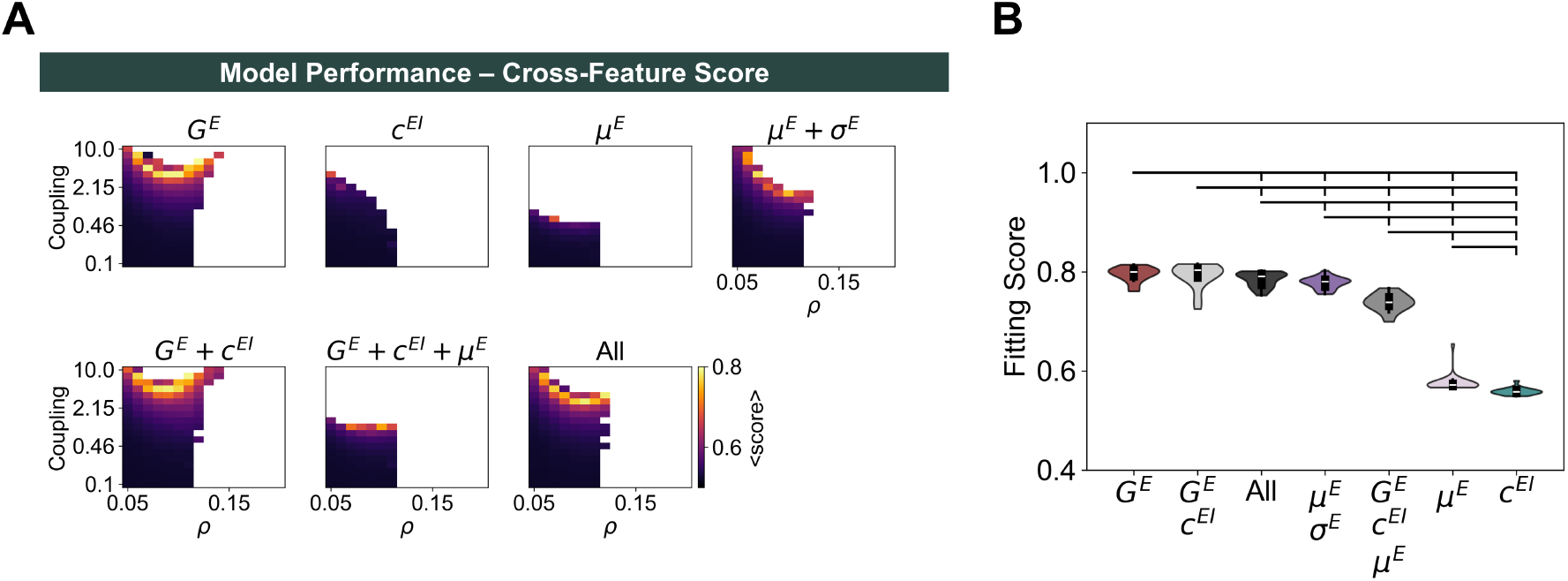
Cross-feature model performance for different homeostatic mechanisms. (A) Parameter spaces representing the cross-feature score for each combination of *C* and *ρ*, averaged across mean delays. Blank spaces represent combinations of *C* and *ρ* for which the homeostatic set point is not valid (i.e. mean firing rates differ from *ρ* by more than 1% in at least one cortical area) (B) Comparison between cross-feature fitting scores at the optimal point of each mechanism of homeostasis. Distributions correspond to simulations with optimal *C* and *ρ* and all values of mean delay yielding a valid network solution. Brackets indicate a significant difference, with p<0.05 from a Mann-Whitney U-test. All p-values were FDR-corrected. All scores in this figure were computed using the following formula: 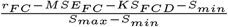. This score represents the norm of a 3D vector where each component represents the performance of the model in representing one of the features of interest, normalized between 0 and 1. *MSE*_*max*_ represents the maximum possible mean squared error between our empirical FC matrix and a given FC matrix.

**Fig. S13.**
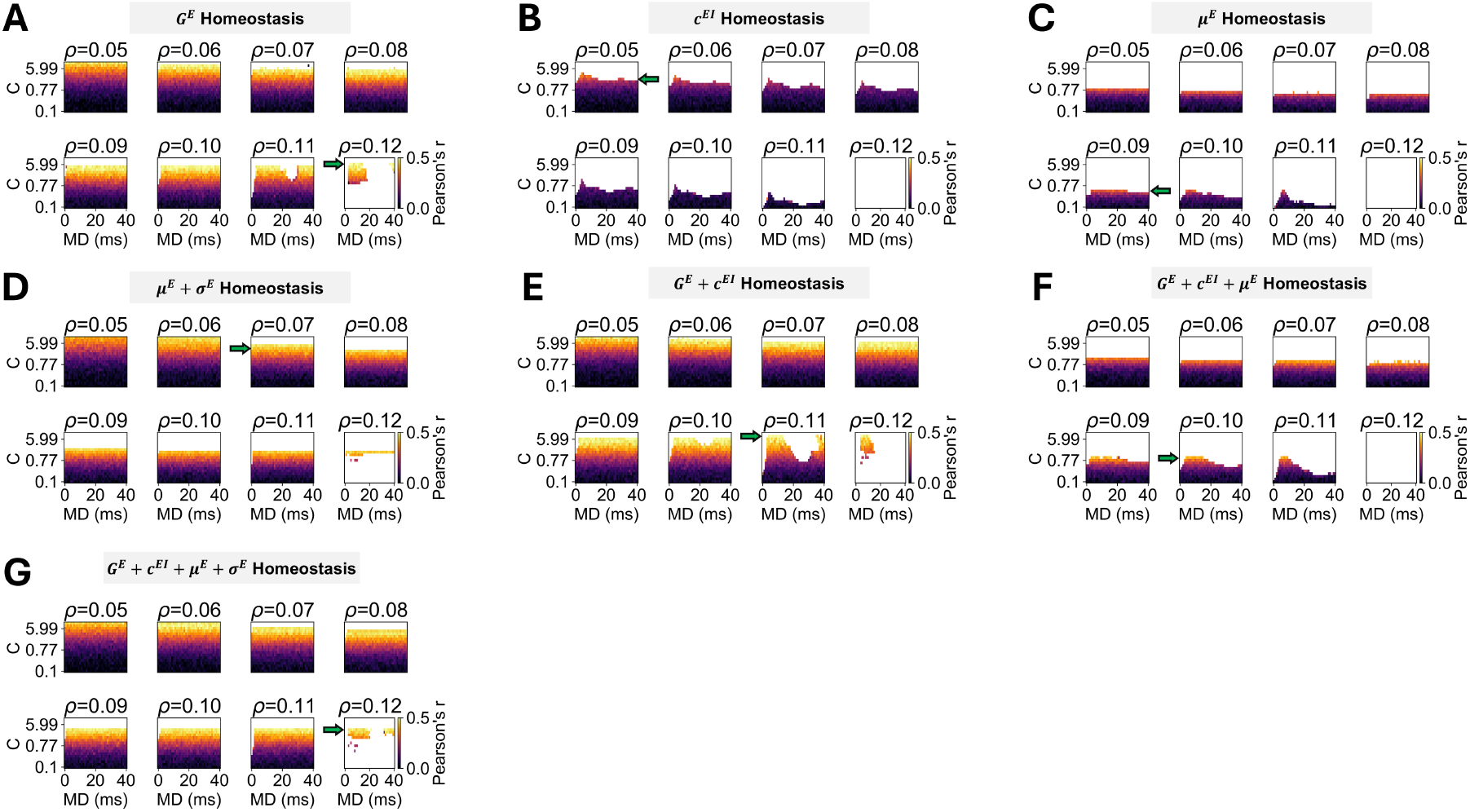
FC fitting of models with different mechanisms of homeostasis. (A) *G*^*E*^ Homeostasis (B) *c*^*EI*^ Homeostasis (C) *µ*^*E*^ Homeostasis (D) *µ*^*E*^ + σ^*E*^ Homeostasis (E) *G*^*E*^ + *c*^*EI*^ Homeostasis (F) *G*^*E*^ + *c*^*EI*^ + *µ*^*E*^ Homeostasis (G) *G*^*E*^ + *c*^*EI*^ + *µ*^*E*^ + σ^*E*^ Homeostasis. Colors represent the correlation coefficient between empirical and simulated FC matrices for each combination of *C* and *ρ*.

**Fig. S14.**
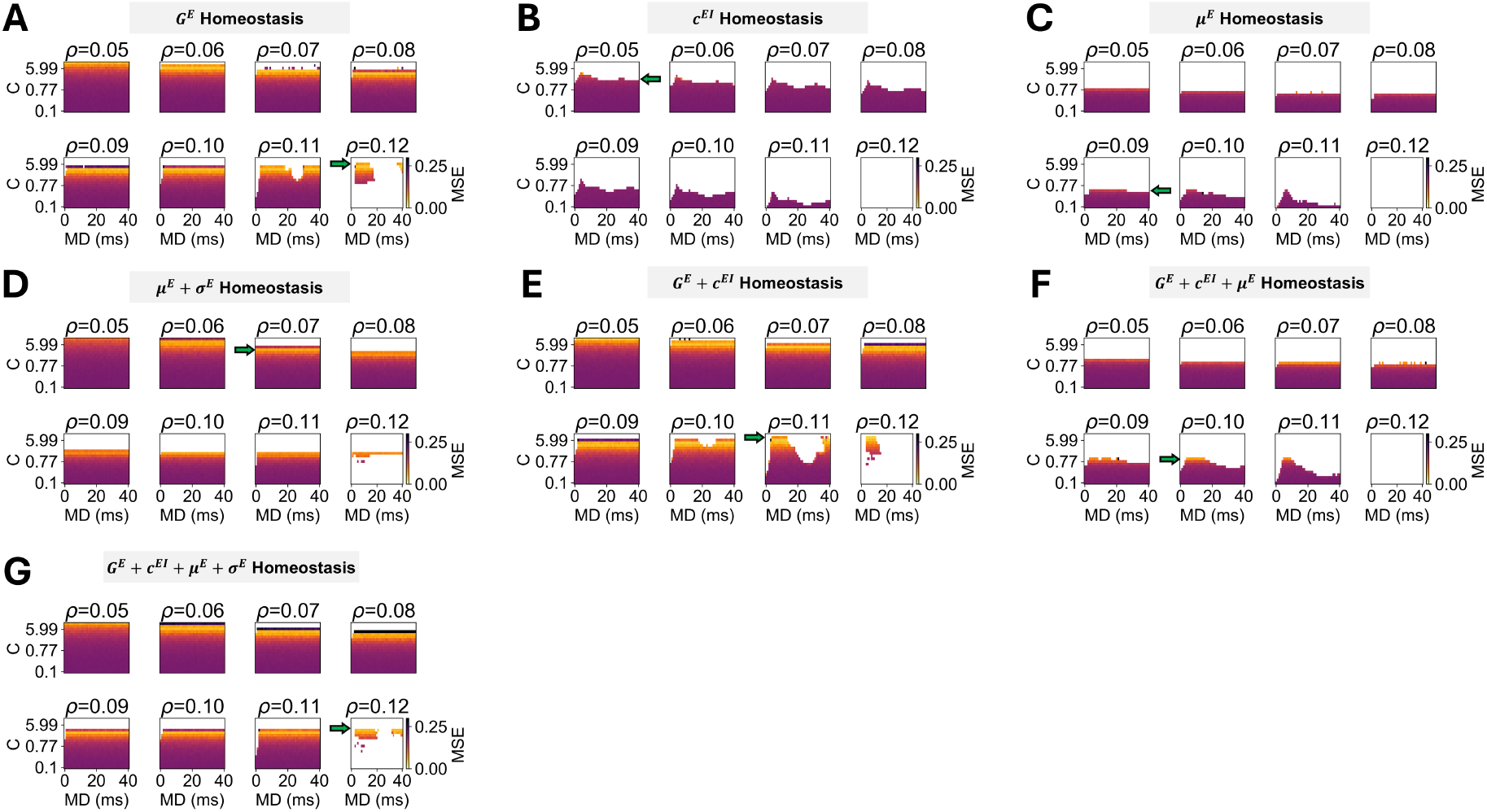
FC fitting of models with different mechanisms of homeostasis. (A) *G*^*E*^ Homeostasis (B) *c*^*EI*^ Homeostasis (C) *µ*^*E*^ Homeostasis (D) *µ*^*E*^ + σ^*E*^ Homeostasis (E) *G*^*E*^ + *c*^*EI*^ Homeostasis (F) *G*^*E*^ + *c*^*EI*^ + *µ*^*E*^ Homeostasis (G) *G*^*E*^ + *c*^*EI*^ + *µ*^*E*^ + σ^*E*^ Homeostasis. Colors represent the mean squared error between empirical and simulated FC matrices for each combination of *C* and *ρ*.

**Fig. S15.**
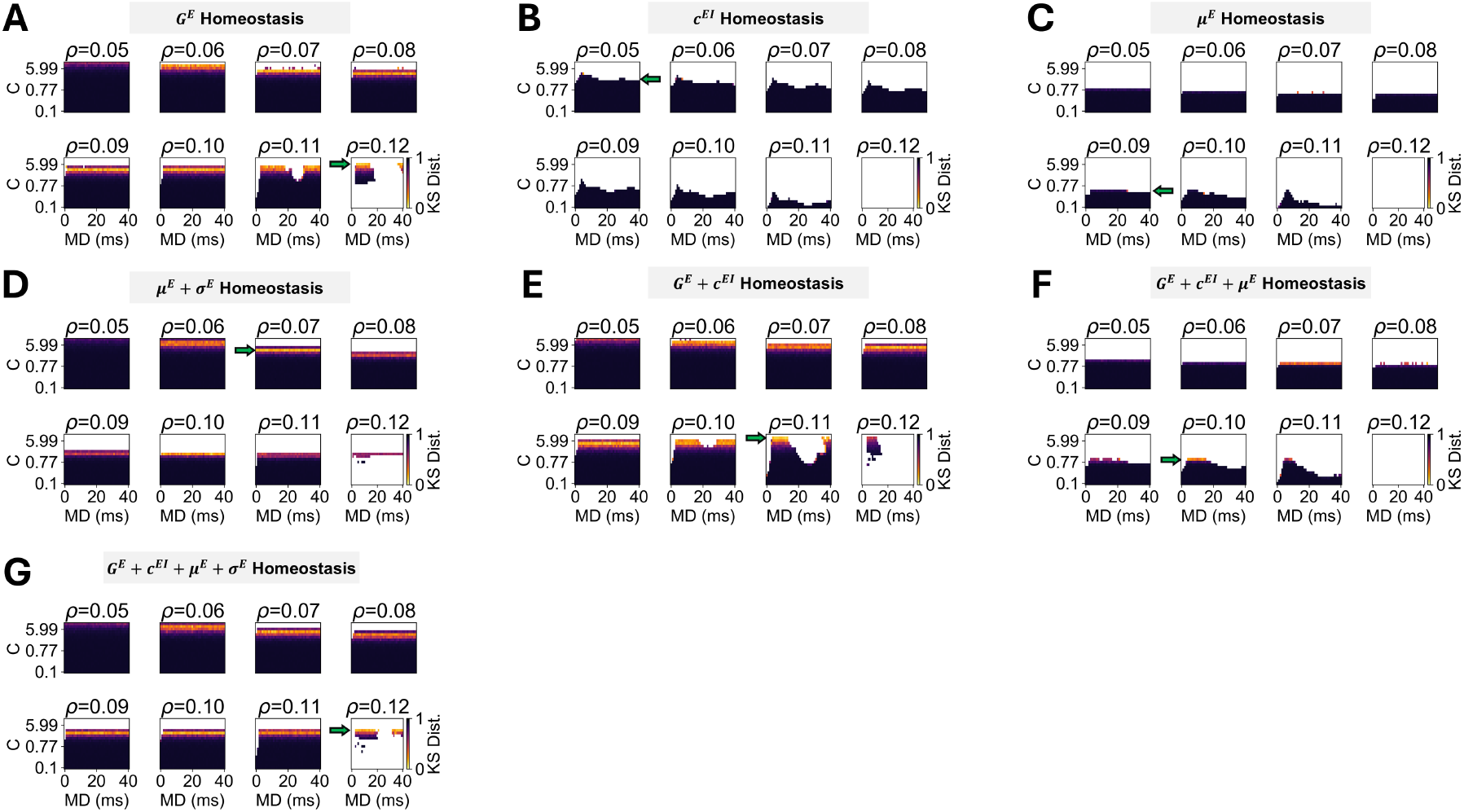
FCD fitting of models with different mechanisms of homeostasis. (A) *G*^*E*^ Homeostasis (B) *c*^*EI*^ Homeostasis (C) *µ*^*E*^ Homeostasis (D) *µ*^*E*^ + σ^*E*^ Homeostasis (E) *G*^*E*^ + *c*^*EI*^ Homeostasis (F) *G*^*E*^ + *c*^*EI*^ + *µ*^*E*^ Homeostasis (G) *G*^*E*^ + *c*^*EI*^ + *µ*^*E*^ + σ^*E*^ Homeostasis. Colors represent the Kolmogorov-Smirnov distance between empirical and simulated FCD distributions for each combination of *C* and *ρ*.

**Fig. S16.**
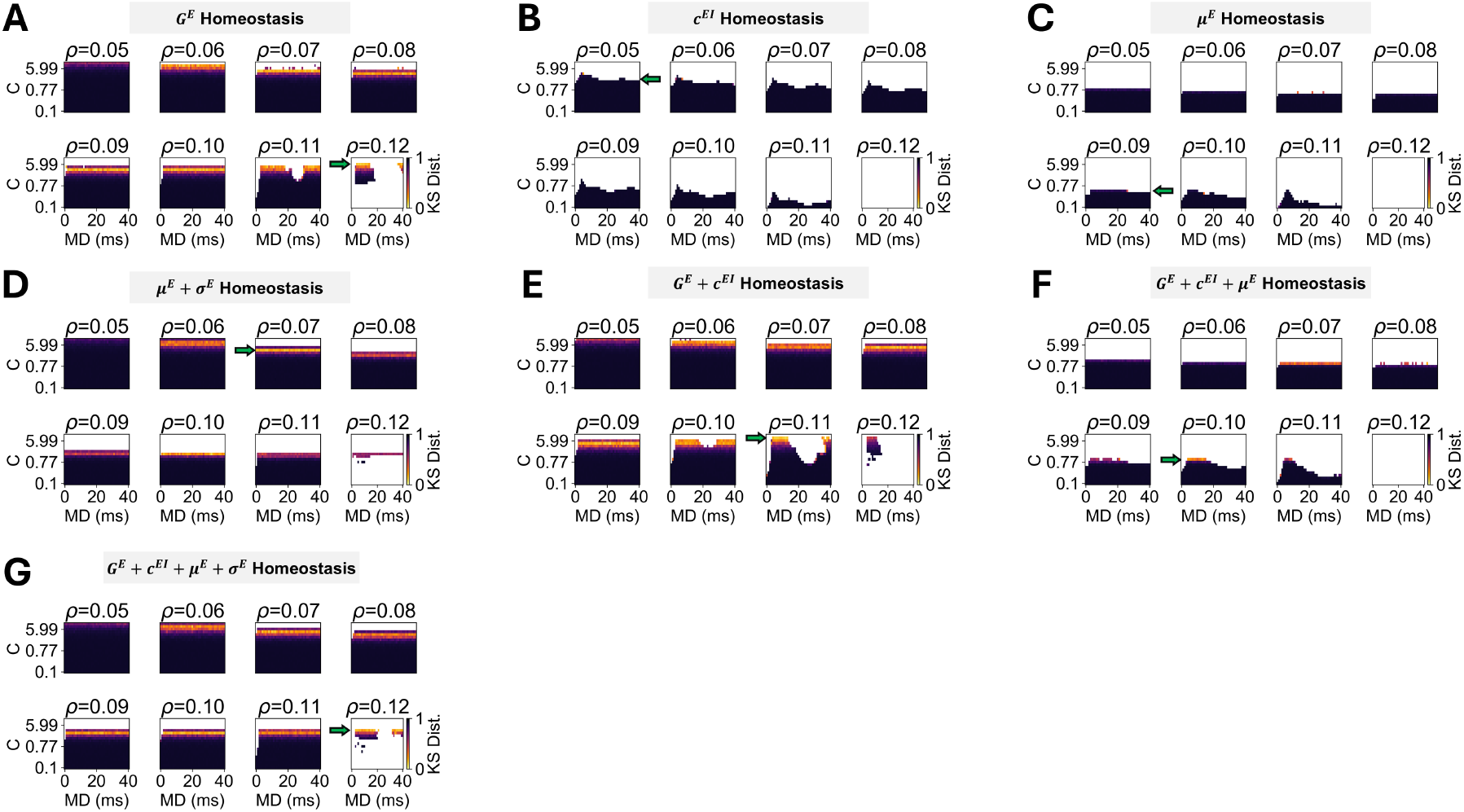
Fitting performance of models with different mechanisms of homeostasis. (A) *G*^*E*^ Homeostasis (B) *c*^*EI*^ Homeostasis (C) *µ*^*E*^ Homeostasis (D) *µ*^*E*^ + σ^*E*^ Homeostasis (E) *G*^*E*^ + *c*^*EI*^ Homeostasis (F) *G*^*E*^ + *c*^*EI*^ + *µ*^*E*^ Homeostasis (G) *G*^*E*^ + *c*^*EI*^ + *µ*^*E*^ + σ^*E*^ Homeostasis. Colors represent the fitting score (*r*_*FC*_ − *MSE*_*F C*_ − *KS*_*FCD*_) for each combination of *C* and *ρ*.

**Fig. S17.**
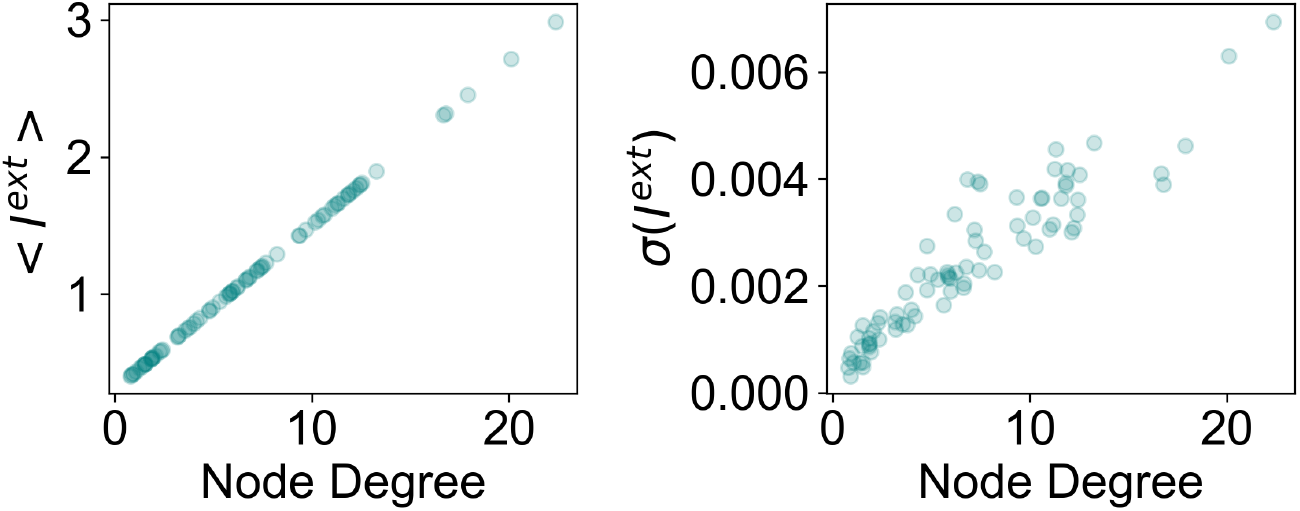
Relationship between node degree, average inputs and input fluctuations. (A) Relationship between node degree (i.e sum of connectivity weights of a node) and the average input received by nodes. (B) Relationship between node degree (i.e sum of connectivity weights of a node) and fluctuations (i.e. standard deviation) of inputs received by a node. These results were obtained from a 60s simulation of the model with homeostasis of *G*^*E*^, *c*^*EI*^ and *µ*^*E*^ + σ^*E*^ at the optimal point (*C* = 3.59, *ρ* = 0.12) and with a mean delay of 40 ms.

**Fig. S18.**
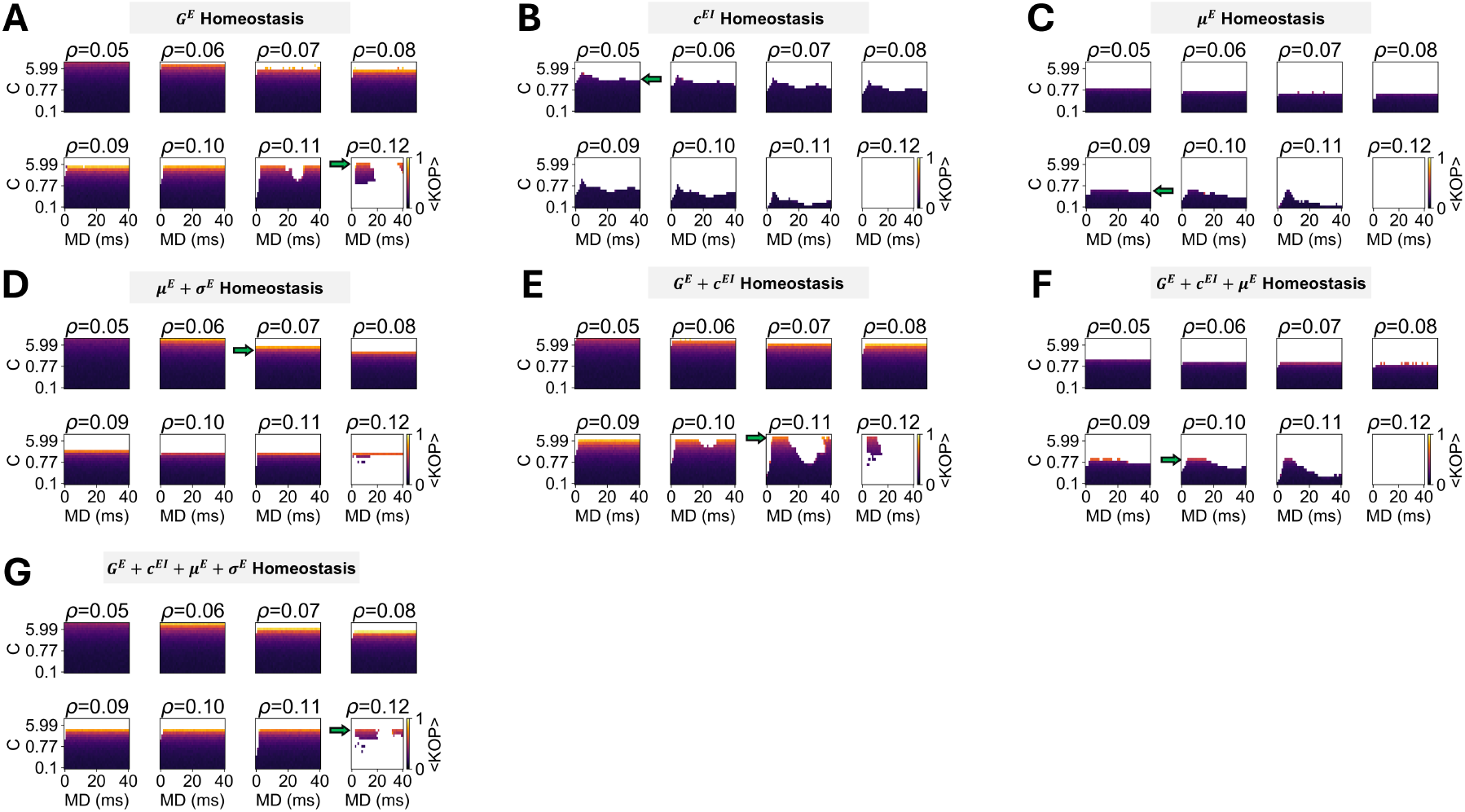
Synchrony of models with different mechanisms of homeostasis. (A) *G*^*E*^ Homeostasis (B) *c*^*EI*^ Homeostasis (C) *µ*^*E*^ Homeostasis (D) *µ*^*E*^ + σ^*E*^ Homeostasis (E) *G*^*E*^ + *c*^*EI*^ Homeostasis (F) *G*^*E*^ + *c*^*EI*^ + *µ*^*E*^ Homeostasis (G) *G*^*E*^ + *c*^*EI*^ + *µ*^*E*^ + σ^*E*^ Homeostasis. Colors represent the mean of the Kuramoto Order Parameter (i.e. synchrony) for each combination of *C* and *ρ*.

**Fig. S19.**
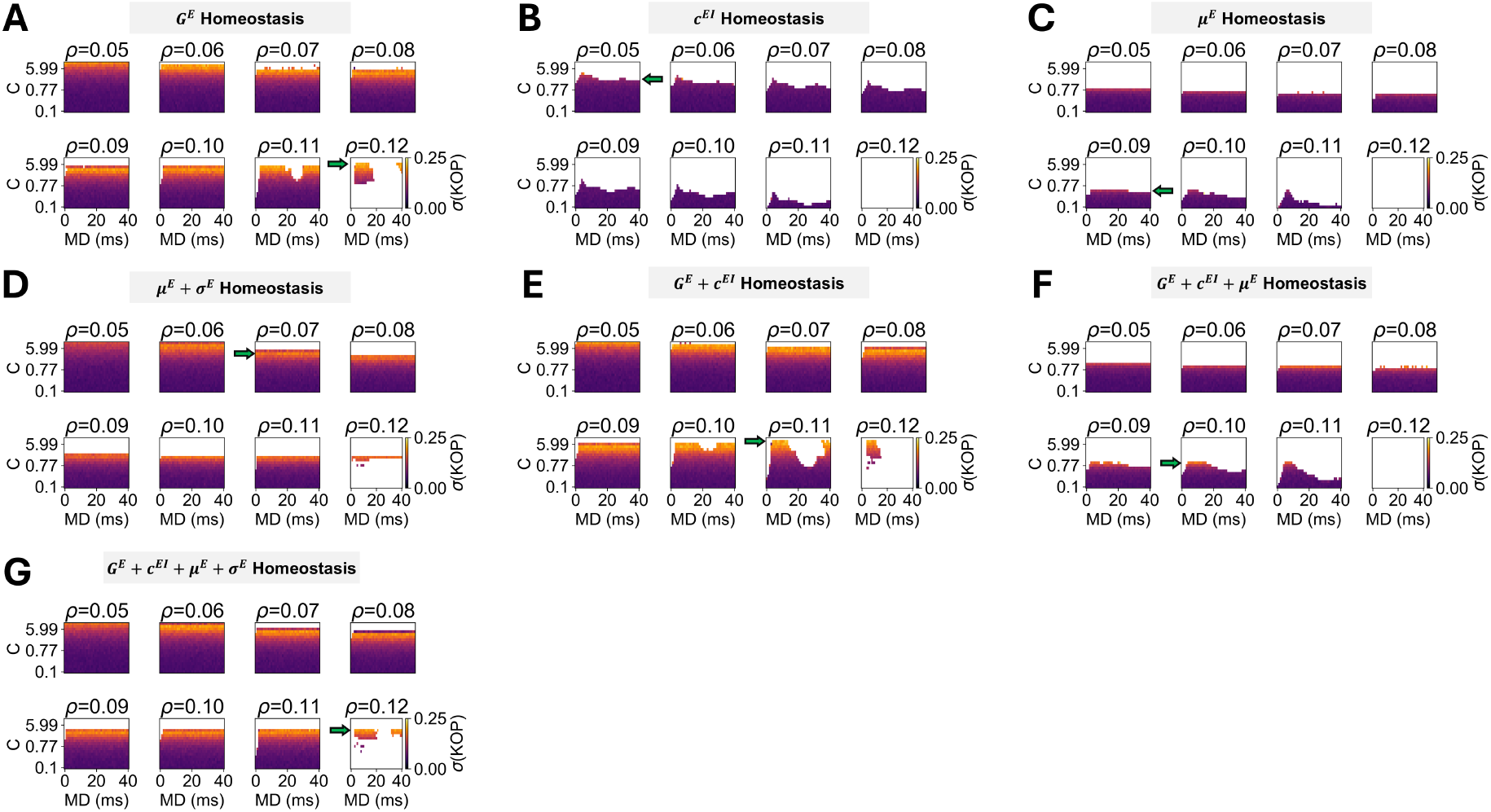
Metastability of models with different mechanisms of homeostasis. (A) *G*^*E*^ Homeostasis (B) *c*^*EI*^ Homeostasis (C) *µ*^*E*^ Homeostasis (D) *µ*^*E*^ + σ^*E*^ Homeostasis (E) *G*^*E*^ + *c*^*EI*^ Homeostasis (F) *G*^*E*^ + *c*^*EI*^ + *µ*^*E*^ Homeostasis (G) *G*^*E*^ + *c*^*EI*^ + *µ*^*E*^ + σ^*E*^ Homeostasis. Colors represent the standard deviation of the Kuramoto Order Parameter (i.e. metastability) for each combination of *C* and *ρ*.

**Fig. S20.**
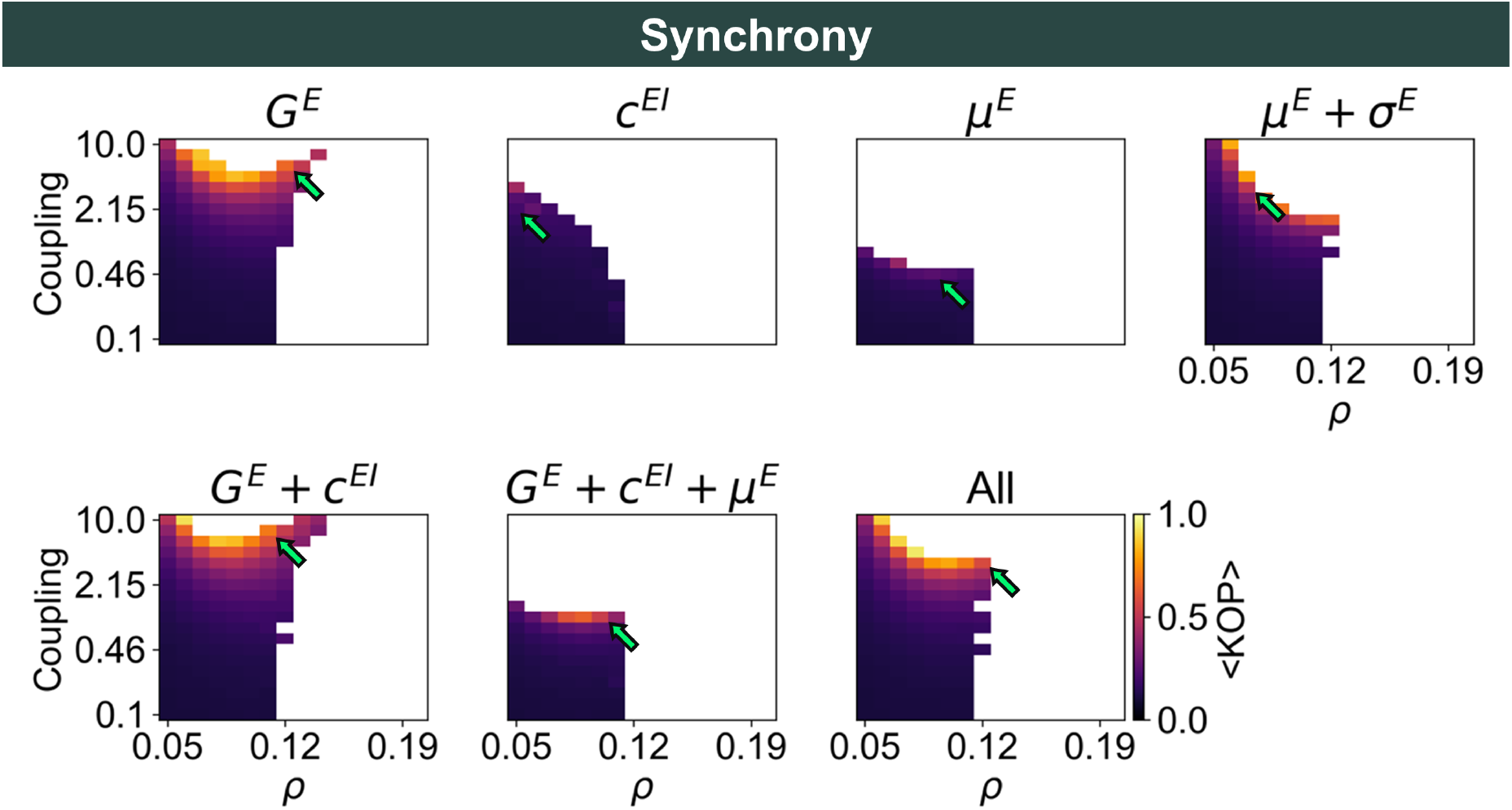
Parameter spaces representing network synchrony (average Kuramoto Order Parameter across time) for each combination of *C* and *ρ*, averaged across mean delays, for models with distinct mechanisms of homeostasis.

**Fig. S21.**
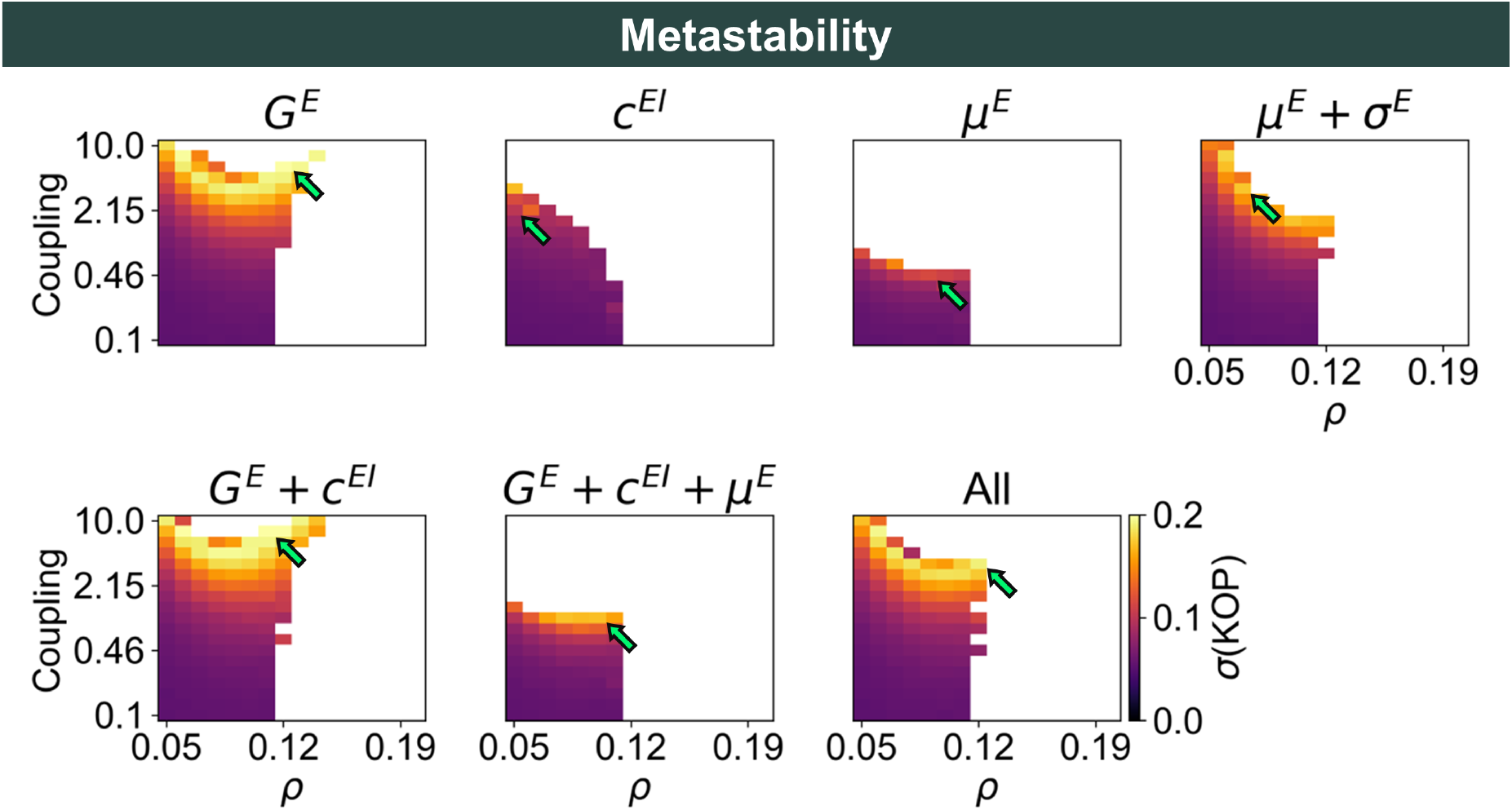
Parameter spaces representing network metastability (standard deviation of Kuramoto Order Parameter across time) for each combination of *C* and *ρ*, averaged across mean delays, for models with distinct mechanisms of homeostasis.

**Fig. S22.**
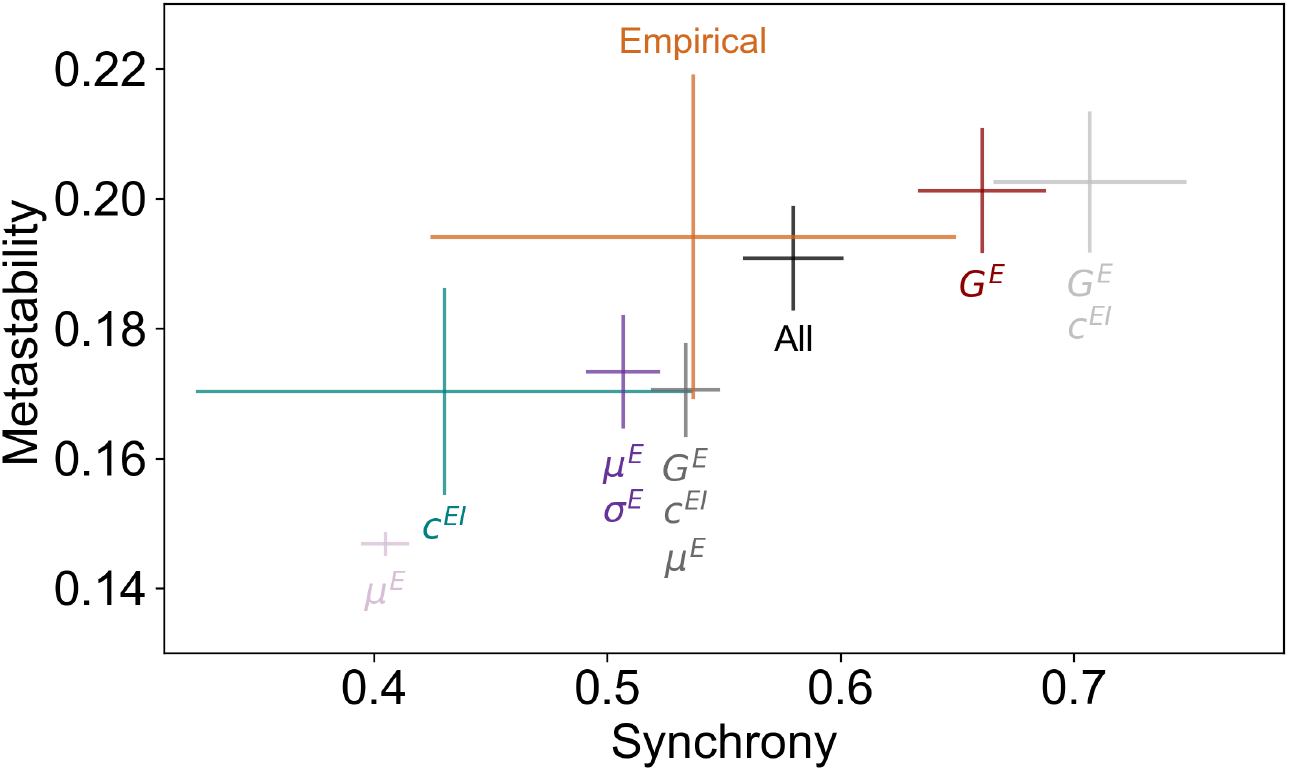
Synchrony and metastability in models with distinct mechanisms of homeostasis and empirical data. The center of the crosses represents the mean synchrony and metastability in models at the optimal working point (*C* and *ρ*) of each mechanism of homeostasis. Conversely, horizontal and vertical bars represent the standard deviation of synchrony and metastability, respectively. Note that the model with all mechanisms of homeostasis is the one that better approximates empirical levels of both synchrony and metastability.

**Fig. S23.**
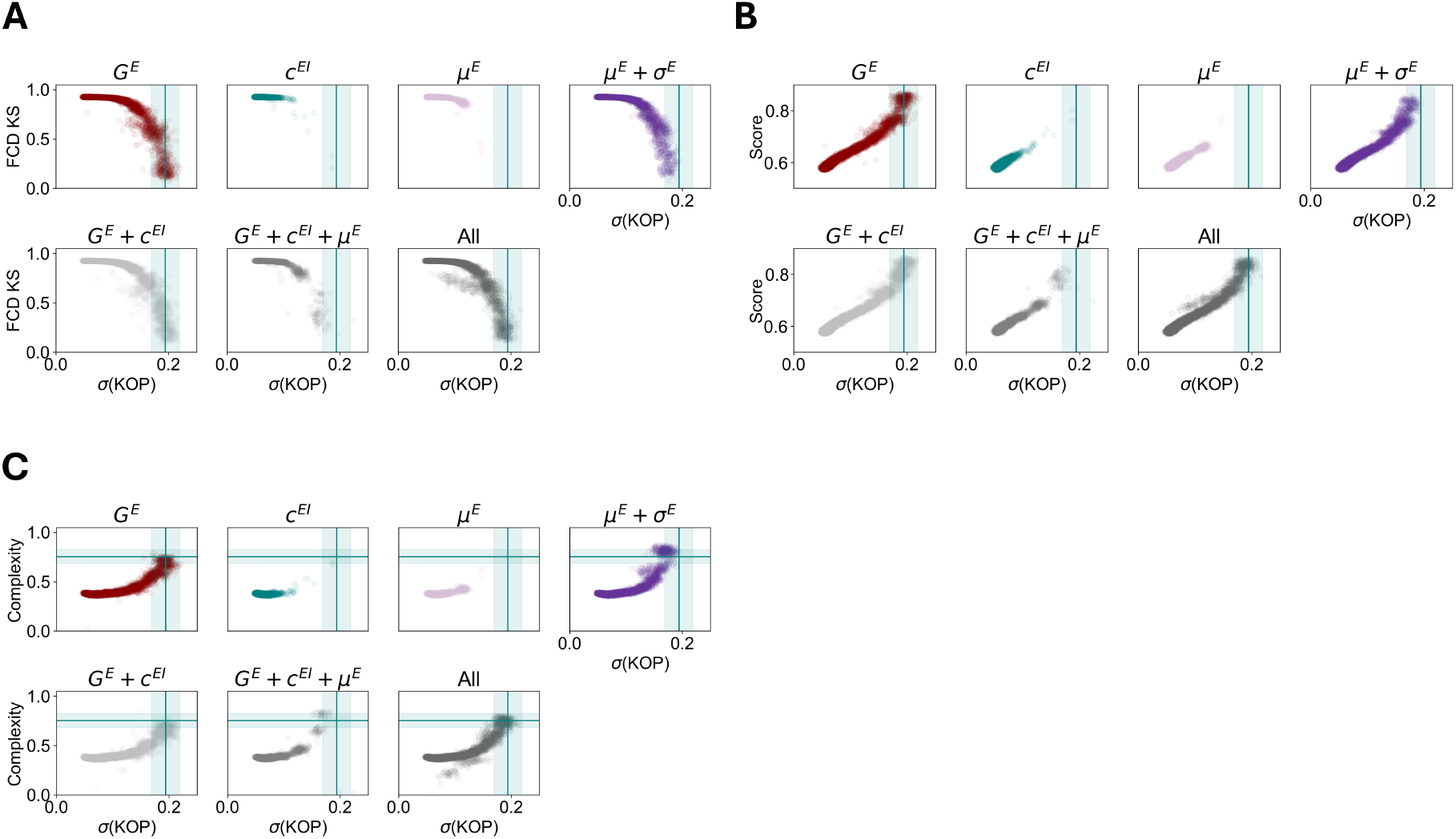
Relationship between metastability, model performance and complexity for all mechanisms of homeostasis. (A) Relationship between metastability and the KS distance between empirical and simulated FCD distributions (B) Relationship between metastability and fitting scores (C) Relationship between metastability and functional complexity. Each data point corresponds to a single simulation. Blue lines and shaded areas represent the mean and standard deviation of metastability and complexity in empirical data.

**Fig. S24.**
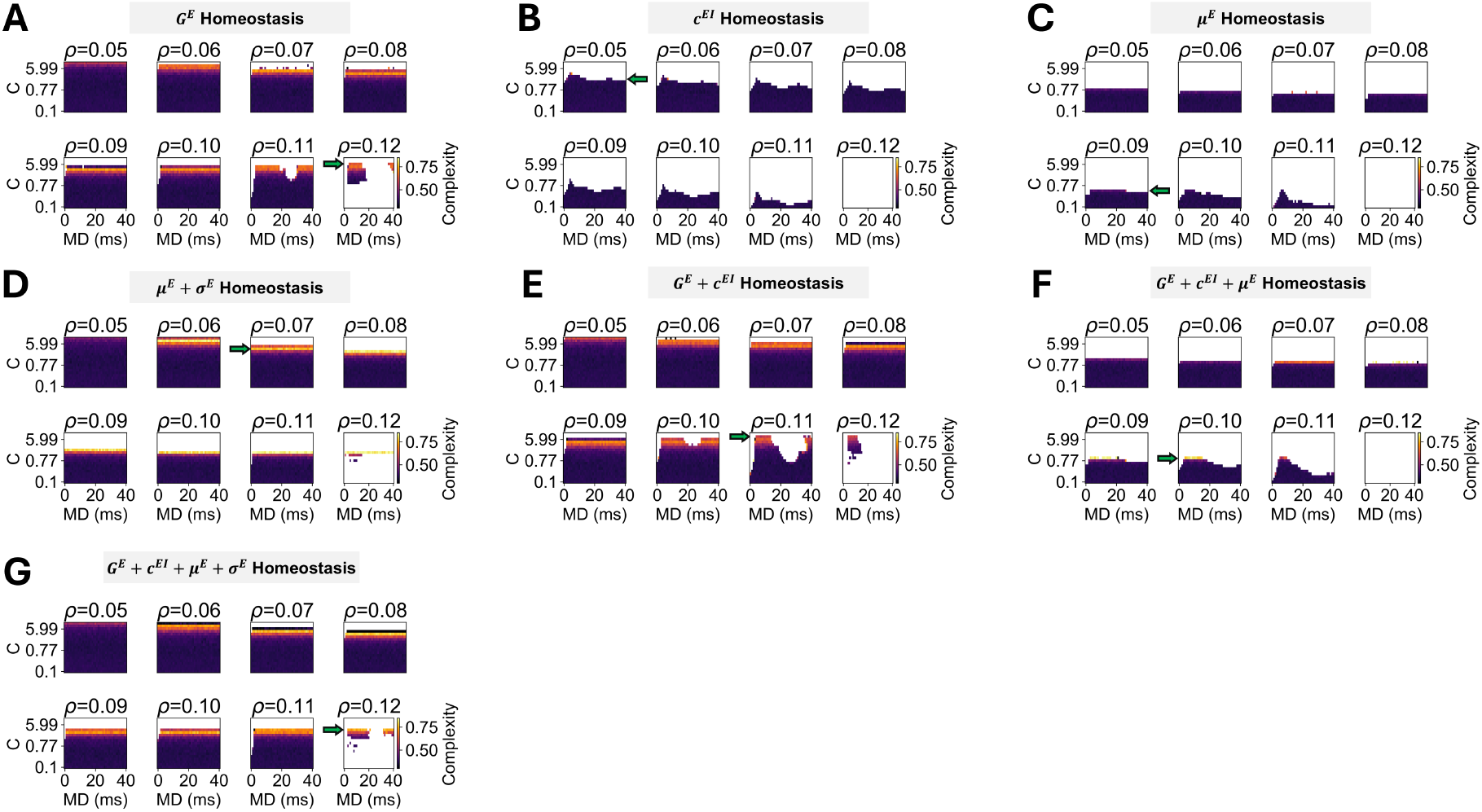
Functional complexity of models with different mechanisms of homeostasis. (A) *G*^*E*^ Homeostasis (B) *c*^*EI*^ Homeostasis (C) *µ*^*E*^ Homeostasis (D) *µ*^*E*^ + σ^*E*^ Homeostasis (E) *G*^*E*^ + *c*^*EI*^ Homeostasis (F) *G*^*E*^ + *c*^*EI*^ + *µ*^*E*^ Homeostasis (G) *G*^*E*^ + *c*^*EI*^ + *µ*^*E*^ + σ^*E*^ Homeostasis. Colors represent the complexity of FC matrices for each combination of *C* and *ρ*.

**Fig. S25.**
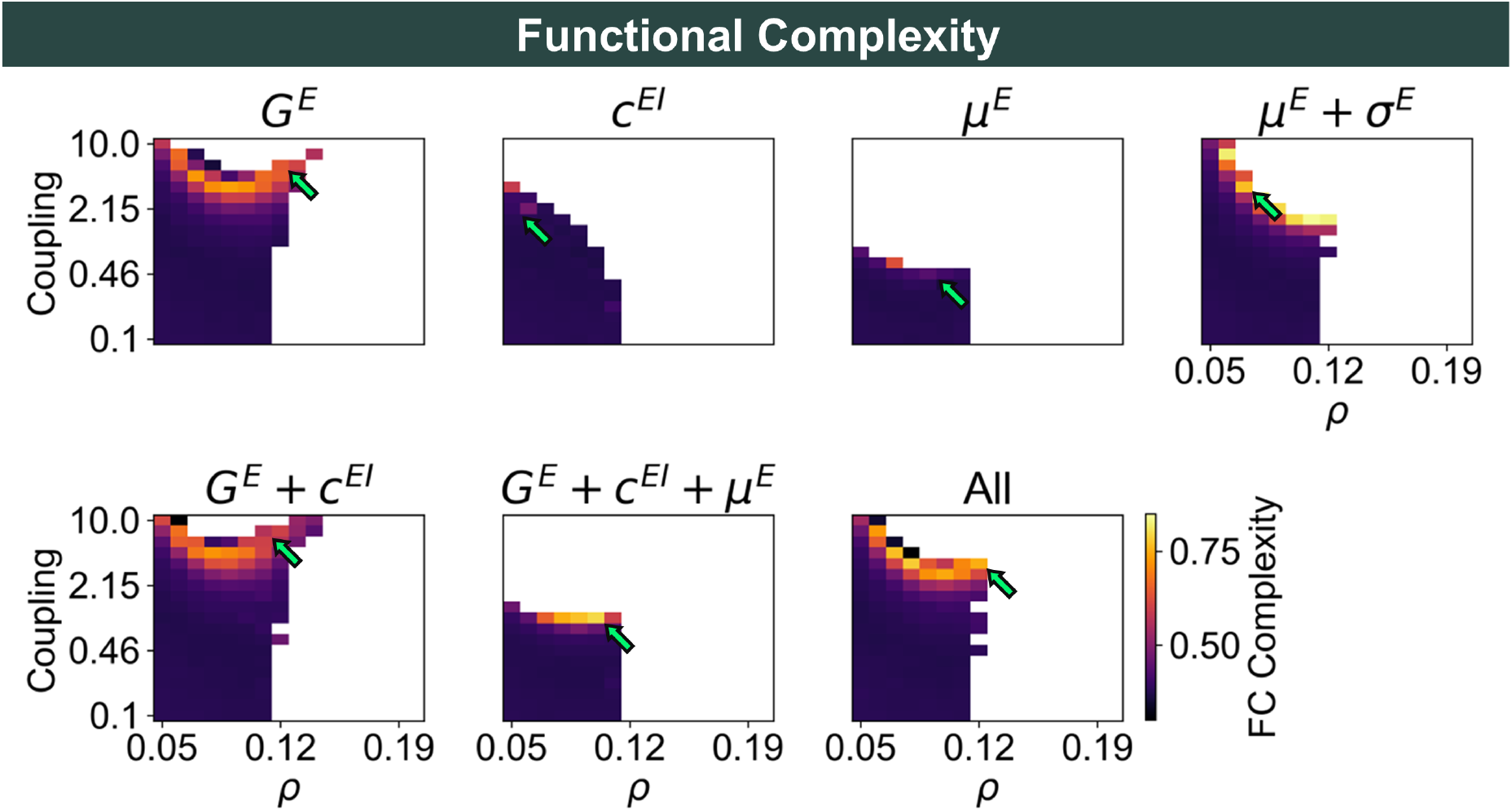
Parameter spaces representing functional complexity for each combination of *C* and *ρ*, averaged across mean delays, for models with distinct mechanisms of homeostasis.

**Fig. S26.**
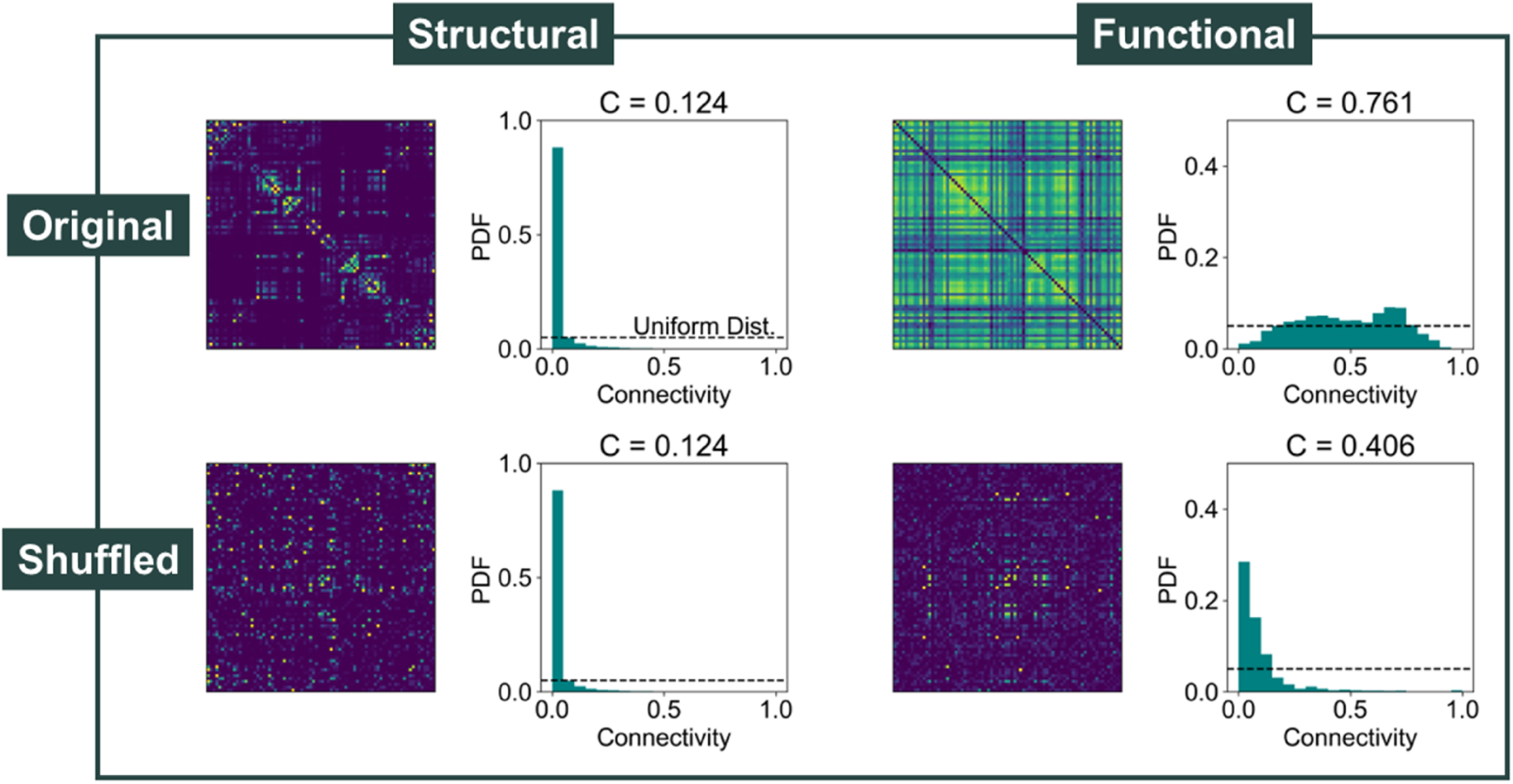
Functional complexity in models with the original and shuffled connectome. (Left) Structural connectivity matrices and respective weight distributions and complexity, corresponding to the original and shuffled connectomes. (Right) Functional connectivity matrices generated by models with the homeostasis of *G*^*E*^, *c*^*EI*^, *µ*^*E*^, and σ^*E*^ with the original and shuffled connectomes and respective weight distributons. We used the optimal models (*C* = 3.59, *ρ* = 0.12) with a mean delay of 40 ms.

**Fig. S27.**
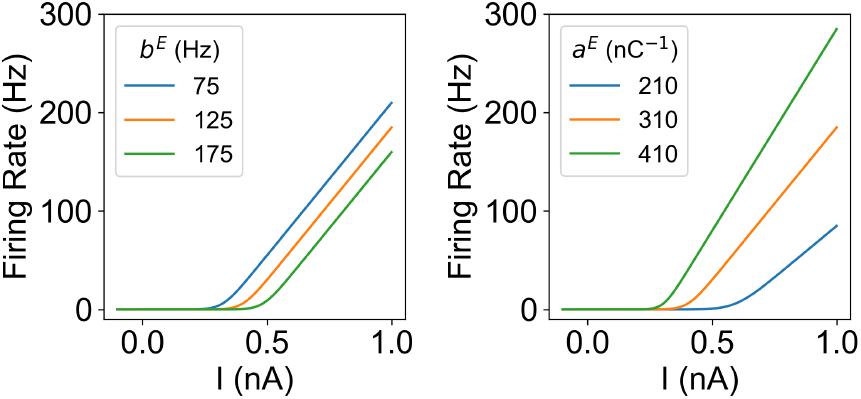
Input-Output function of the Wong-Wang model with different parameters. Value of *F*^*E*^ (*x*) in the Wong-Wang model for different values of the firing threshold *b*^*E*^ (Left) and slope *a*^*E*^ (Right)

**Fig. S28.**
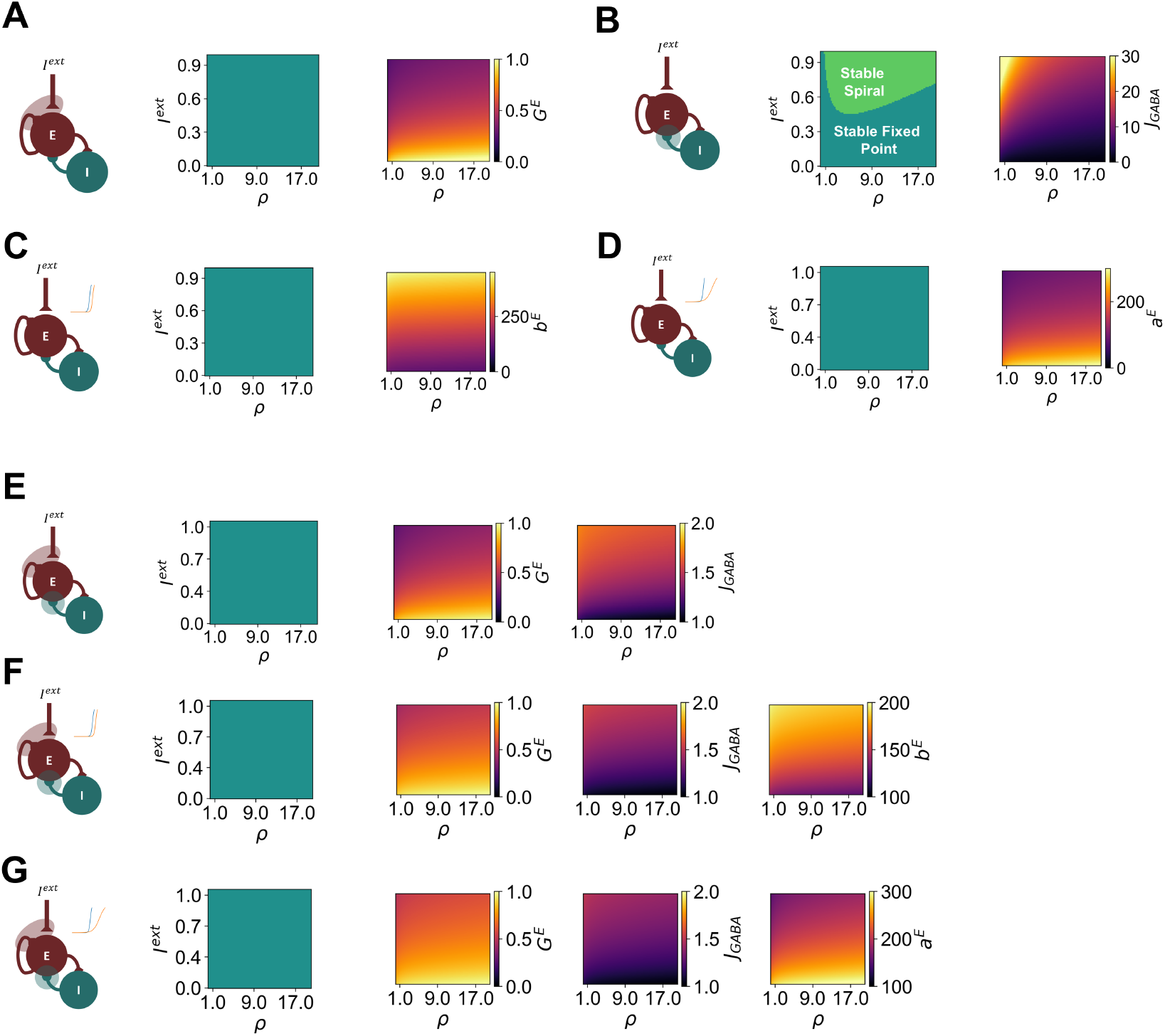
Dynamics of local Wong-Wang circuits under different mechanisms of E-I homeostasis. We present results for circuits under the homeostasis of excitation (A), inhibition (B), firing threshold (C), firing slope (D), excitation and inhibition (E), excitation, inhibition and firing threshold (F), and excitation, inhibition and firing slope (G). In each subplot, we present the result of linear stability analysis on the left, with colors representing the different dynamical regimes. In addition, we present the values of the parameters being modulated by E-I homeostasis. Note that, in the Wong-Wang model, E-I homeostasis does not have a strong effect on local dynamics, since they remain in a stable regime for all tested combinations of parameters and homeostatic mechanisms. As opposed to the Wilson-Cowan model, there is no Hopf-bifurcation.

**Fig. S29.**
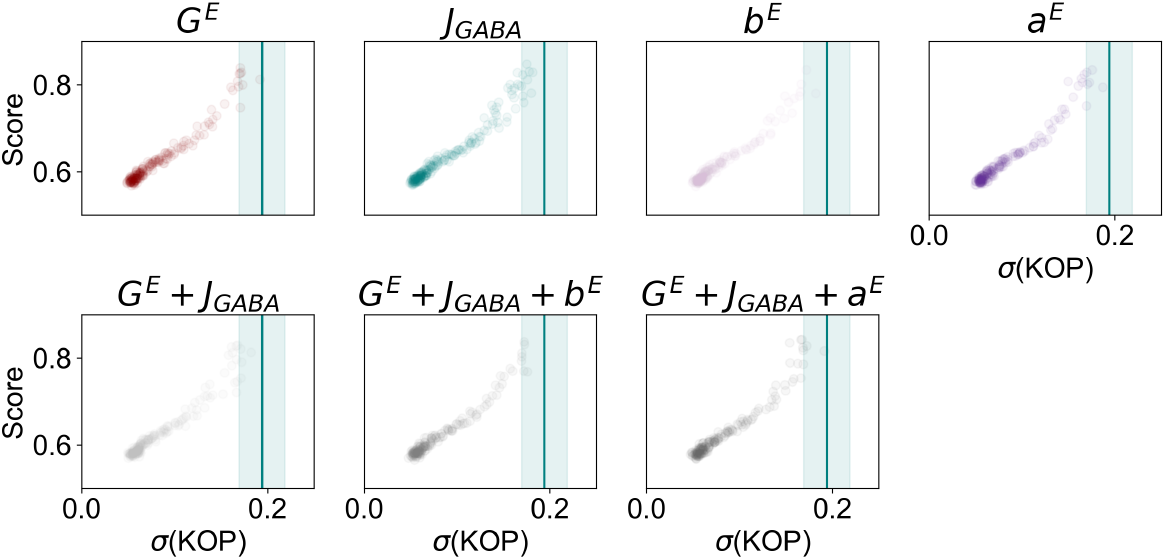
Relationship between metastability and model performance for all mechanisms of homeostasis in the Wong-Wang model. Blue lines and shaded areas represent the mean and standard deviation of metastability in empirical data.

**Fig. S30.**
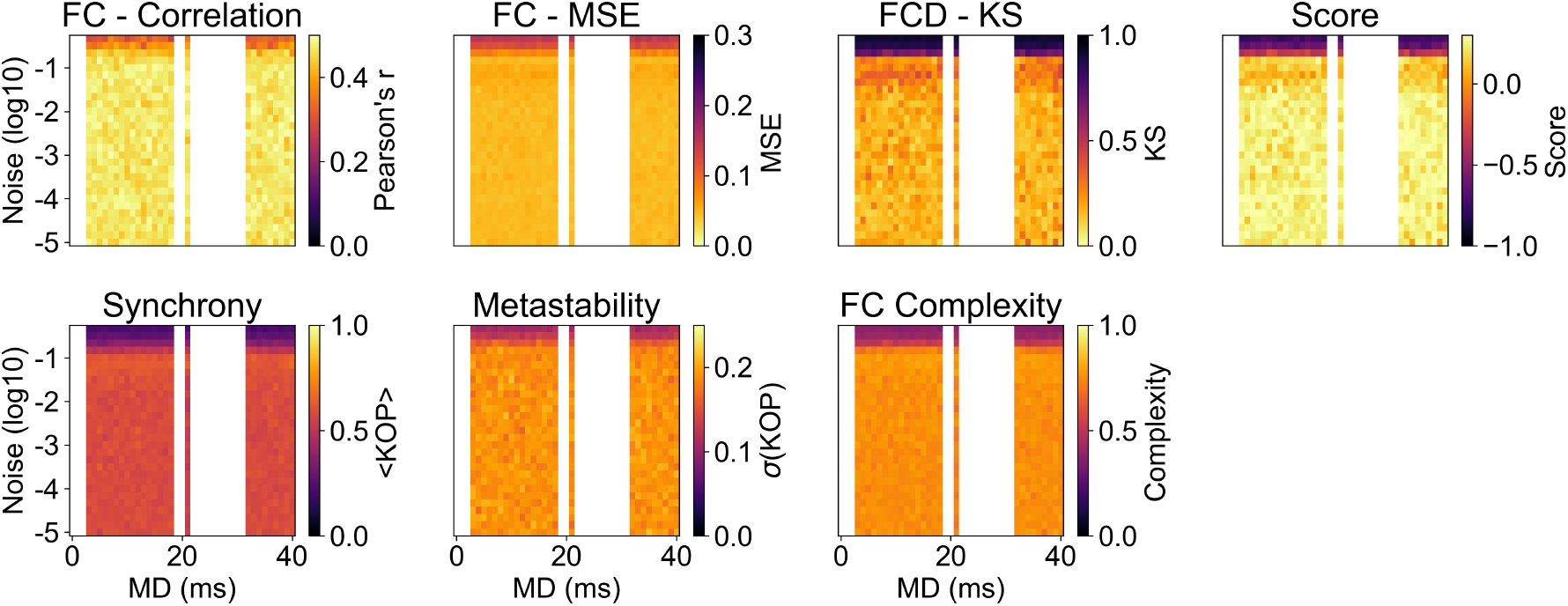
Parameter space of model with all mechanisms of homeostasis with different noise levels. We present the results for all combinations of mean delay and noise variance for models with *C* = 3.59 and *ρ* = 0.12 and across all mean delays yielding a valid solution of the model. Note that the empty areas in the parameter space relate to mean delays for which |⟨*r*^*E*^ ⟩ − *ρ*|*/ρ >* 0.01 in at least one node.

**Fig. S31.**
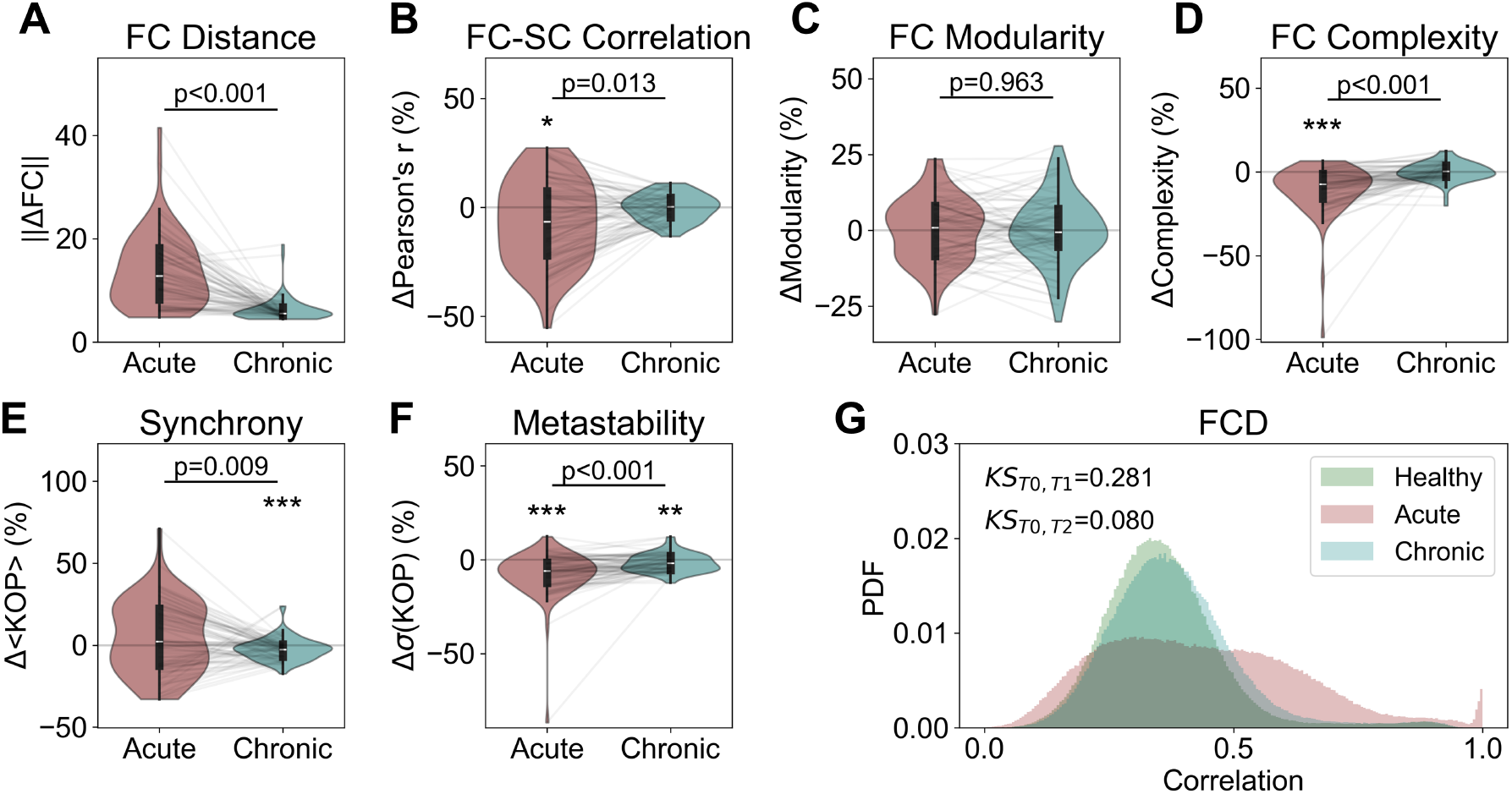
Disruption and recovery of network properties after focal lesion in models with homeostasis of excitation (*G*^*E*^) (A) Distance from baseline of FC matrices in acute and chronic simulations (B) Change from baseline (in percentage) of FC-SC correlation in the acute and chronic simulations (C) Same as B, for FC Modularity (D) Same as B, for FC complexity (E) Same as B, for synchrony (D) Same as B, for metastability (F) Distributions of FCD correlations in all stages of our lesion simulations. In addition, we present the Kolmogorov-Smirnov distance between distributions at baseline and acute/chronic simulations. P-values represent the result of a Wilcoxon Ranked-sum test. For 1-sample tests, asterisks represent the significance of a Wilcoxon Ranked sum test (* p<0.05; ** p<0.01; *** p<0.001). All p-values were FDR-corrected.

**Fig. S32.**
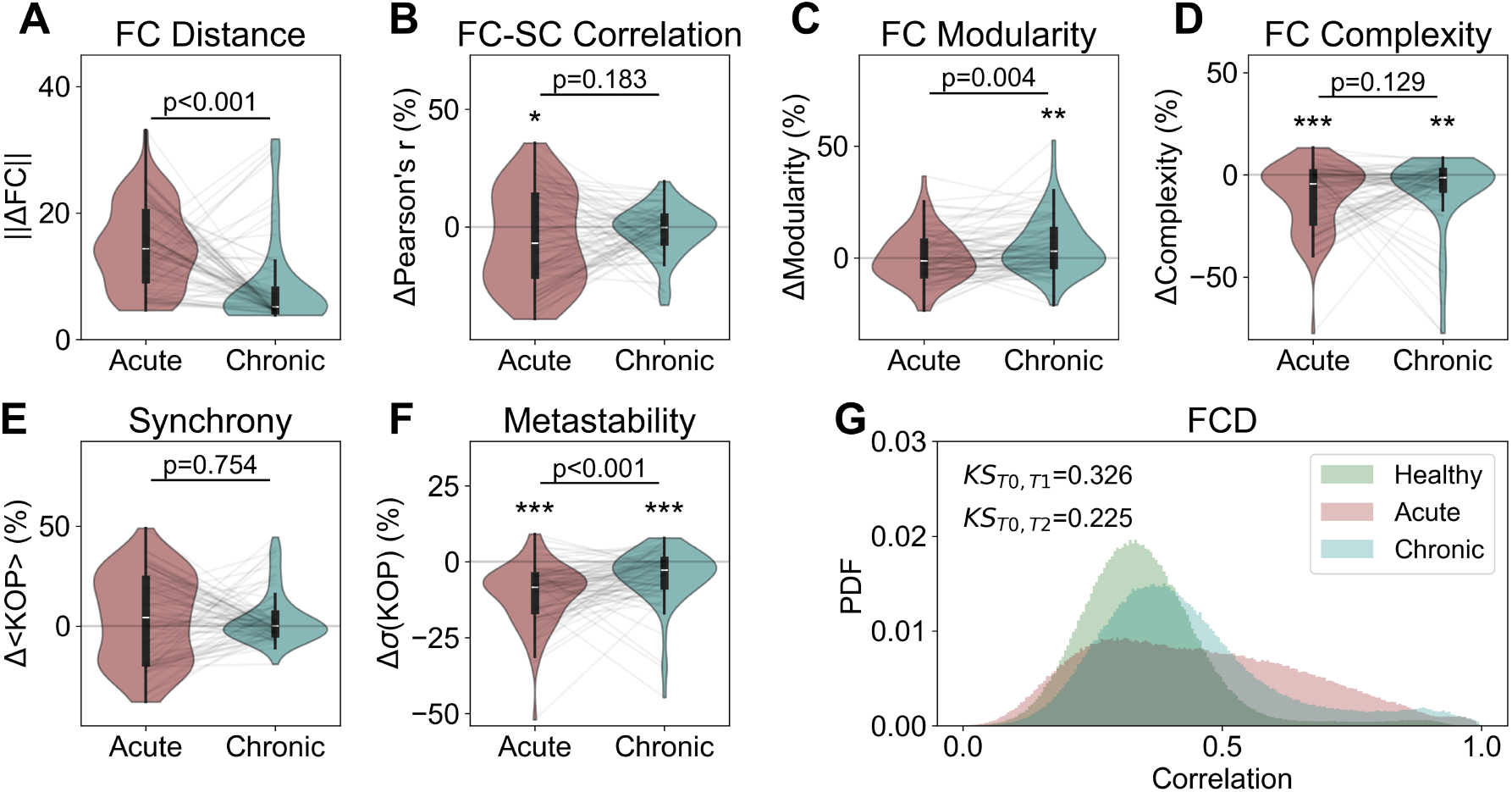
Disruption and recovery of network properties after focal lesion in models with homeostasis of excitation (*G*^*E*^) and inhibition (*c*^*EI*^) (A) Distance from baseline of FC matrices in acute and chronic simulations (B) Change from baseline (in percentage) of FC-SC correlation in the acute and chronic simulations (C) Same as B, for FC Modularity (D) Same as B, for FC complexity (E) Same as B, for synchrony (D) Same as B, for metastability (F) Distributions of FCD correlations in all stages of our lesion simulations. In addition, we present the Kolmogorov-Smirnov distance between distributions at baseline and acute/chronic simulations. P-values represent the result of a Wilcoxon Ranked-sum test. For 1-sample tests, asterisks represent the significance of a Wilcoxon Ranked sum test (* p<0.05; ** p<0.01; *** p<0.001). All p-values were FDR-corrected.

**Table S1.** Optimal parameters, fitting scores, and metastability in the Wong-Wang model for each mechanism of E-I homeostasis in comparison to empirical data. Fitting scores correspond to 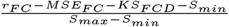, where *r*_*FC*_ is the Pearson’s correlation between empirical and simulated FC matrices, *MSE*_*F C*_ their mean squared error, *KS*_*F CD*_ the Kolmogorov-Smirnov distance between FCD distributions and *S*_*max*_*/S*_*min*_ the maximum/ minimum of *r*_*FC*_ − *MSE*_*F C*_ − *KS*_*F CD*_. Values are presented as mean±sd over 10 simulations. Values in bold represent a significant difference

| Homeostatic Mechanism | $C$ | $\rho$ (Hz) | Score | $r_{FC}$ | $MSE_{FC}$ | $KS_{FCD}$ | Metastability |
| --- | --- | --- | --- | --- | --- | --- | --- |
| $G^E$ | 0.125 | 4.0 | 0.831±0.007 | 0.41±0.01 | 0.066±0.002 | 0.19±0.04 | **0.170±0.005** |
| $J_{GABA}$ | 0.261 | 5.0 | 0.826±0.010 | 0.43±0.01 | 0.063±0.002 | 0.21±0.03 | **0.169±0.006** |
| $b^E$ | 0.178 | 10.0 | 0.825±0.006 | 0.39±0.01 | 0.068±0.002 | 0.20±0.03 | **0.165±0.006** |
| $a^E$ | 0.261 | 3.0 | 0.833±0.007 | 0.41±0.01 | 0.067±0.003 | 0.18±0.03 | **0.172±0.006** |
| $G^E + J_{GABA}$ | 0.178 | 6.0 | 0.820±0.012 | 0.41±0.02 | 0.073±0.004 | 0.24±0.05 | **0.164±0.006** |
| $G^E + J_{GABA} + b^E$ | 0.681 | 1.0 | 0.833±0.008 | 0.43±0.01 | 0.068±0.005 | 0.20±0.04 | **0.174±0.006** |
| $G^E + J_{GABA} + a^E$ | 0.215 | 4.0 | 0.828±0.007 | 0.40±0.01 | 0.067±0.002 | 0.19±0.02 | **0.171±0.008** |
| Empirical | - | - | - | - | - | - | 0.194±0.025 |

**Table S2.** Default Parameters of the Wilson-Cowan Model.

| Model Parameters |  |
| --- | --- |
| $c^{EE}$ | 3.5 |
| $c^{IE}$ | 3.75 |
| $c^{EI}$ | 2.5 |
| $G^E$ | 1 |
| $\mu^E/\mu^I$ | 1 |
| $\sigma^E/\sigma^I$ | 0.25 |
| $\tau^E$ | 2.5 ms |
| $\tau^I$ | 5.0 ms |

**Table S3.**
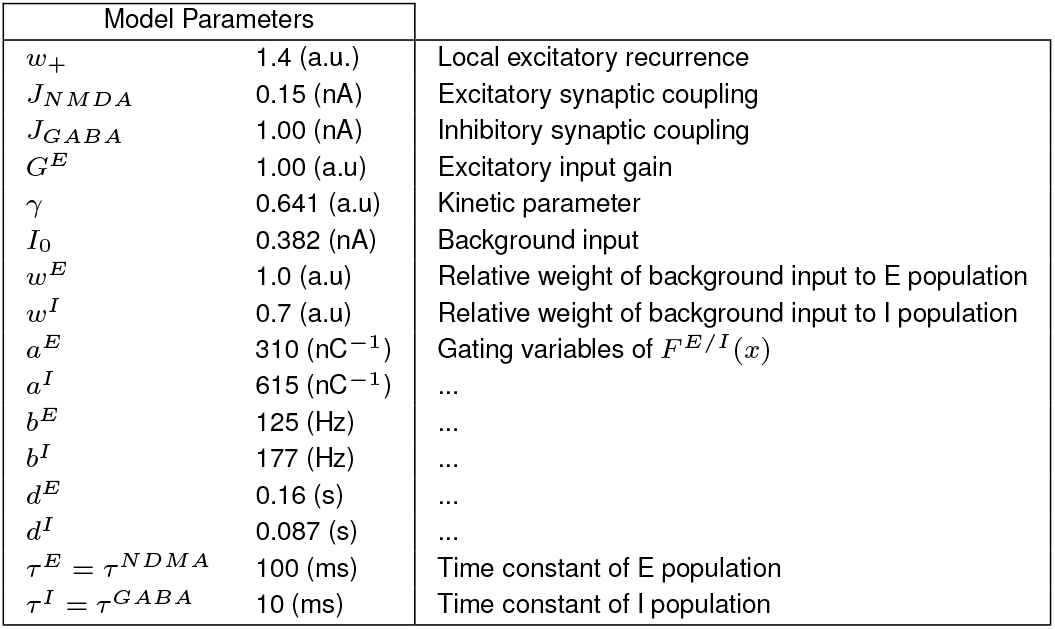
Default Parameters of the Wong-Wang Model.

## Notes

https://gitlab.com/francpsantos/wc_network_homeostasis

### Competing Interest Statement

FPS was employed by Eodyne Systems SL. PFMJV is the founder and shareholder of Eodyne Systems S.L., which brings scientifically validated neurorehabilitation and education technologies to society.

### Summary of Updates

- A new subfigure 1B, illustrating the effects of different homeostatic mechanisms on the local dynamics of our model - New results comparing the performance of our model and the reduced Wong-Wang model, which has slower local dynamics and no gamma oscillations, together with the respective methods and discussion. - New fitting score, normalized between 0 and 1, facilitating its interpretation - More stringent constraints on the choice of optimal 𝑪 and 𝝆, requiring that simulations with such parameters are valid for more than 10 different values of mean delay.

